# Epigenetic predictors of species maximum lifespan and other life history traits in mammals

**DOI:** 10.1101/2023.11.02.565286

**Authors:** Caesar Z. Li, Amin Haghani, Qi Yan, Ake T. Lu, Joshua Zhang, Zhe Fei, Jason Ernst, X. William Yang, Vadim N. Gladyshev, Ken Raj, Andrei Seluanov, Vera Gorbunova, Steve Horvath

## Abstract

Maximum lifespan is an intrinsic characteristic of a biological species and is defined as the longest time an individual of a species has been reported to survive. By analyzing 15K samples derived from 348 mammalian species representing 25 taxonomic orders we previously identified CpG methylation sites associated with maximum lifespan. Here we present accurate DNA methylation-based (DNAm) predictors of maximum lifespan (r=0.89), average gestation time (r=0.96), and age at sexual maturity (r=0.85). Our DNAm maximum lifespan predictor indicates a potential innate longevity advantage for females over males in 17 mammalian species such as humans, red deer, and cattle. The DNAm maximum lifespan predictions do not vary significantly by caloric restriction and partial reprogramming. Genetic disruptions in the somatotropic axis, which includes growth hormone, IGF-1, and their related receptors, have an impact on DNAm maximum lifespan only in select mouse tissues. Cancer mortality rates in major mammalian orders show no correlation with our epigenetic estimates of life history traits. The DNAm maximum lifespan predictor does not detect variation in lifespan between individuals of the same species, such as between the breeds of dogs. We also present the first prototypes of accurate pan mammalian DNAm predictors of sex and tissue type.

Collectively, our findings indicate that maximum lifespan is determined, at least in part, by an epigenetic signature that is an intrinsic property of each species and is distinct from the signatures that relate to individual lifespan, which is unaffected by interventions influencing the mortality risk of individuals.

## INTRODUCTION

Maximum lifespan varies dramatically across mammalian species: the cinereus shrew lives less than 1.9 years while bowhead whales can live for at least 211 years (*1*). The species appear to exhibit a maximum lifespan – an intrinsic characteristic of a biological species defined as the longest time an individual of a species has been reported to survive. However, the molecular mechanisms that determine it remain poorly understood (*2, 3*), despite prior studies correlating maximum lifespan with specific molecular processes and life history strategies (*4-6*). Many have suggested that epigenetic mechanisms play a role in determining lifespan (*7-15*). However, prior studies of cross-species variation in methylation patterns suffer from low sample size and heterogeneity in data acquisition methods.

To facilitate rigorous methylation studies of life history traits, the Mammalian Methylation Consortium generated an unprecedented and homogeneous data set of DNA methylation at well conserved loci across 348 mammals using a tailor-made DNA methylation measurement platform (*16*). Other reports by the Consortium have described pan-mammalian age-related methylation changes, epigenetic aging clocks, phylo-epigenetic trees and unsupervised machine learning approaches that were brought to bear on the analyses of this dataset (*17, 18*). In recent publications by our Mammalian Methylation Consortium, we released a DNA methylation dataset (n=15,456 tissue samples) (*17, 18*). These prior investigations uncovered individual cytosines and modules that correlate with maximum lifespan, gestation time, and age at sexual maturity.

In this study, we pivot our analytical approach. Rather than seeking individual CpGs tied to maximum lifespan and other life-history traits, we develop regularized multivariate regression models that estimate maximum lifespan and other characteristic traits of species. Drawing on statistical terminology, our previous work focused on univariate analysis (specifically, the selection of CpGs) and CpG modules (*18*). In contrast, here we utilize multivariate regression models to predict maximum lifespan (the dependent variable), based on highly conserved cytosines (the independent variables or covariates) simultaneously. Using this approach, we successfully developed methylation-based predictors of time-related life history traits: maximum lifespan, gestation time, and age at sexual maturity across mammalian species. Next, we characterized these new epigenetic biomarkers with regards to a variety of conditions ranging from demographic characteristics (age, sex, human mortality risk), and interventions that modulate murine lifespan.

## RESULTS

### DNA methylation data from 348 mammalian species

Leveraging our publicly accessible data from the Mammalian Methylation Consortium, we focused on highly conserved cytosine methylation profiles from n = 15K DNA samples. These samples spanned 59 unique tissue types and originated from 348 distinct mammalian species across 25 taxonomic orders. In total, the Mammalian Consortium profiled 25 of the 26 mammalian taxonomic orders as catalogued in the Mammal Diversity Database (Version 1.8, 2022), with marsupial moles being the only exception.

These methylation profiles were obtained using the mammalian methylation array, a tailor-made DNA array developed for the consortium’s objectives (*16*). This array efficiently gauges the methylation levels of roughly 36,000 highly conserved CpG sites. These CpGs are flanked by 50 base-pair DNA sequences that are remarkably conserved across various mammalian species.

### Universal predictors of sex and tissue type

The mammalian array-generated DNA methylation data proves highly effective in accurately classifying sample species, sex, and tissue. This is supported by our random forest predictors, which boast an out-of-bag accuracy rate of over 99% (**table 1**). Importantly, we crafted universal sex predictors grounded in CpG methylation levels that are applicable to all mammalian species, barring marmosets (**table 1**). It’s widely recognized that mosaicism in marmosets hinders the creation of methylation-based sex predictors for them (*19*). We previously postulated that the inability to build methylation-based predictors of sex in marmosets is due to their nature as hematopoietic chimeras. Specifically, littermates in marmosets exchange stem cells through placental anastomoses during development, as discussed in (*19*).

**Table 1.** Sex and pan tissue predictors performance. The table summarizes test set prediction results for regularized-regression-based predictors and out-of-bag prediction results for random-forest-based predictors. Test sets are randomly partitioned into equal 10 folds of the entire data set. At each iteration, within the 90% training set, 10-fold-validation was employed to select the penalization parameter for the regularized regression (sex predictor).

| Predicted Outcome | Predictor Framework | Method | Test Set / Out-of-bag Prediction Accuracy |
| --- | --- | --- | --- |
| Sex (Female = yes/no) | Elastic Net | 10 fold validation | 98.53% |
| Species | *RF Predictor | *100 trees | 99.94% |
| Tissue | *RF Predictor | 100 trees | 98.22% |
| Taxonomic Order | *RF Predictor | 100 trees | 99.97% |

Note: \*RF: Random Forest predictor; \*100: each random forest was calibrated to use this many decision trees with a reasonable run time; random forest unbiased prediction accuracy estimate is calculated as follows; first, summarize by calculating the median of each category's out-of-bag prediction errors, subtracted by unity, across the 500 trees; second, use these category mean accuracies to find the overall median-of-means accuracy.

Our universal tissue predictors, based on methylation, are likely influenced by species variations, potentially making them less precise than the universal sex predictors. While we offer these tissue predictors to the community as potential tools for identifying human platemap errors, we advise users to be aware of the potential species-related confounding factors associated with these predictors.

### Multivariate predictors of life history traits

Since we aimed to focus on species traits, we first removed the confounding effect of sex and tissue type by averaging across these variables. Specifically, we calculated the mean methylation value for each CpG within each species, producing a summarized dataset in which each data point corresponds to a species’ average methylation level per CpG (**table S1**). In addition to this overarching dataset, we curated two more specialized datasets: one stratified by both species and tissue type and another that exclusively focuses on younger samples that were derived from animals that are both not yet sexually mature and under 5 years of age.

We employed three distinct penalized regression models to predict the log-transformed values of maximum lifespan, gestation time, and age at sexual maturity for each species. The trait values for these species were derived from the latest version of the anAge database (*2, 18*). For the convenience of our readers, we have included these values in **table S1** and **table S2**. The resultant epigenetic predictors showcased high accuracy as evidenced by the leave-one-species-out (LOSO) or leave-one-clade-out (LOCO) cross-validation. For instance, the predicted log maximum lifespans aligned closely with those recorded in anAge, exhibiting a Pearson’s correlation of R = 0.89 (see **Fig. 1a,b**). An alternative method for assessing predictive precision entails dividing the data into training and test subsets. Utilizing our 70%-30% training-test random partitioning of species, we observed comparably robust correlations for the log maximum lifespan in both subsets (training set, R = 0.98, **Fig. 2a**; test set, R = 0.88, **Fig. 2a,b**).

**Fig. 1.**
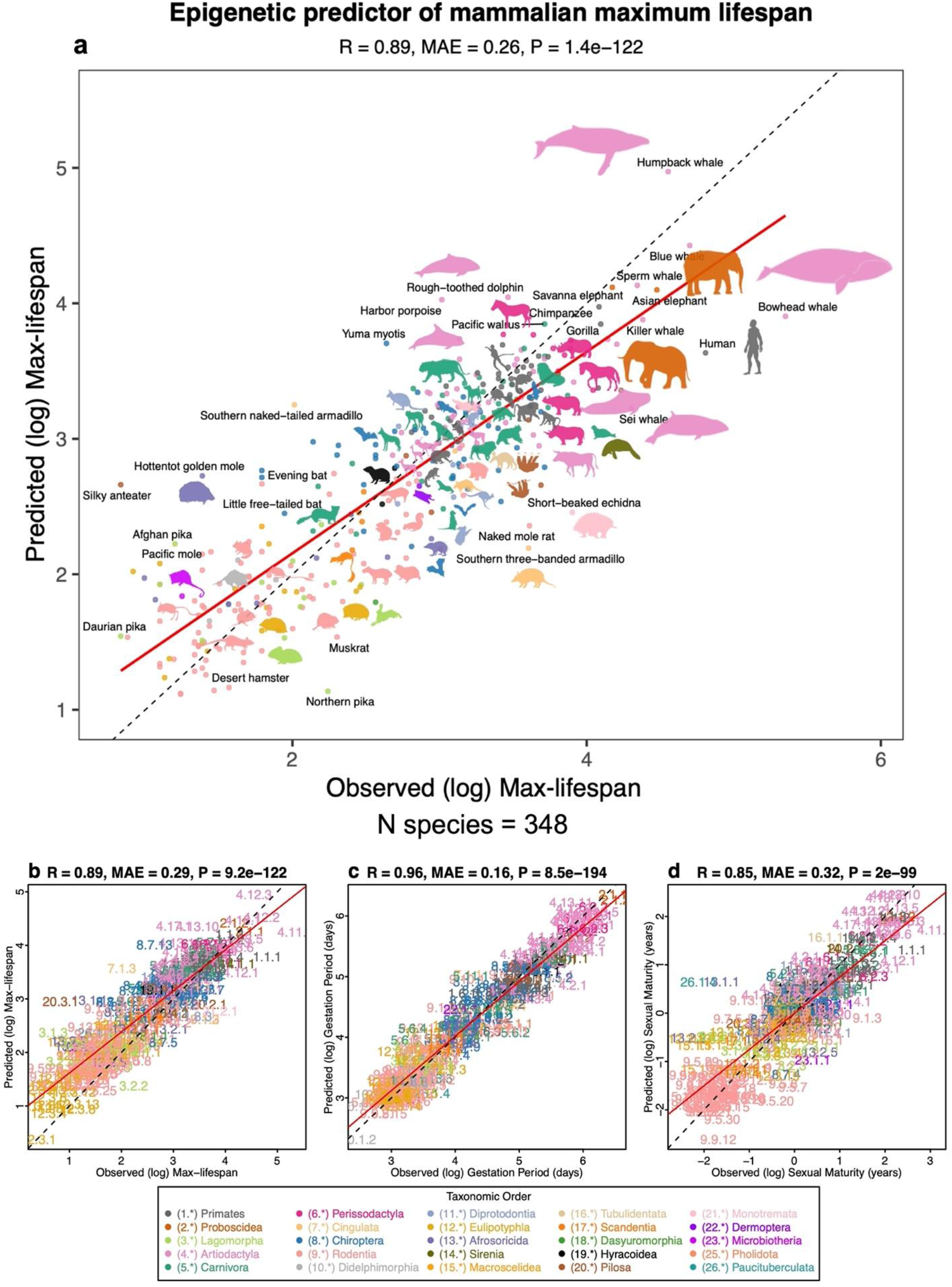
Multivariate analysis of life history traits using epigenetic predictors. This figure delves into the Leave-One-Species-Out (LOSO) cross-validation analysis of epigenetic predictors. It focuses on the log-transformed (base e) estimates for various life history traits, including: **a,b**, Maximum lifespan (in years) **c**, Gestation time (in days) **d**, Age at sexual maturity (in years). Each species in the scatter plot panels is symbolized by a specific number. The whole number (integer) part of this numeric representation corresponds to its taxonomic order. These numbers, color-coded by their respective taxonomic orders, link to distinct species. For detailed numeric values, refer to **table S4**. The title atop each panel provides essential statistical data: the Pearson correlation coefficient, median absolute error (MAE), and a two-sided uncorrected p-value. Consistency in color representation for taxonomic orders is maintained throughout this and other related figures. To comprehend the species-specific numeric designations in the scatter plots, readers can refer to the accompanying figure legends that annotate the common names and taxonomic orders (**fig. S8**). A dotted line within the scatter plots illustrates the line of perfect prediction, while the solid red line portrays the fitted linear regression. Animal silhouettes featured are sourced from the Phylopic database (https://www.phylopic.org/).

**Fig. 2.**
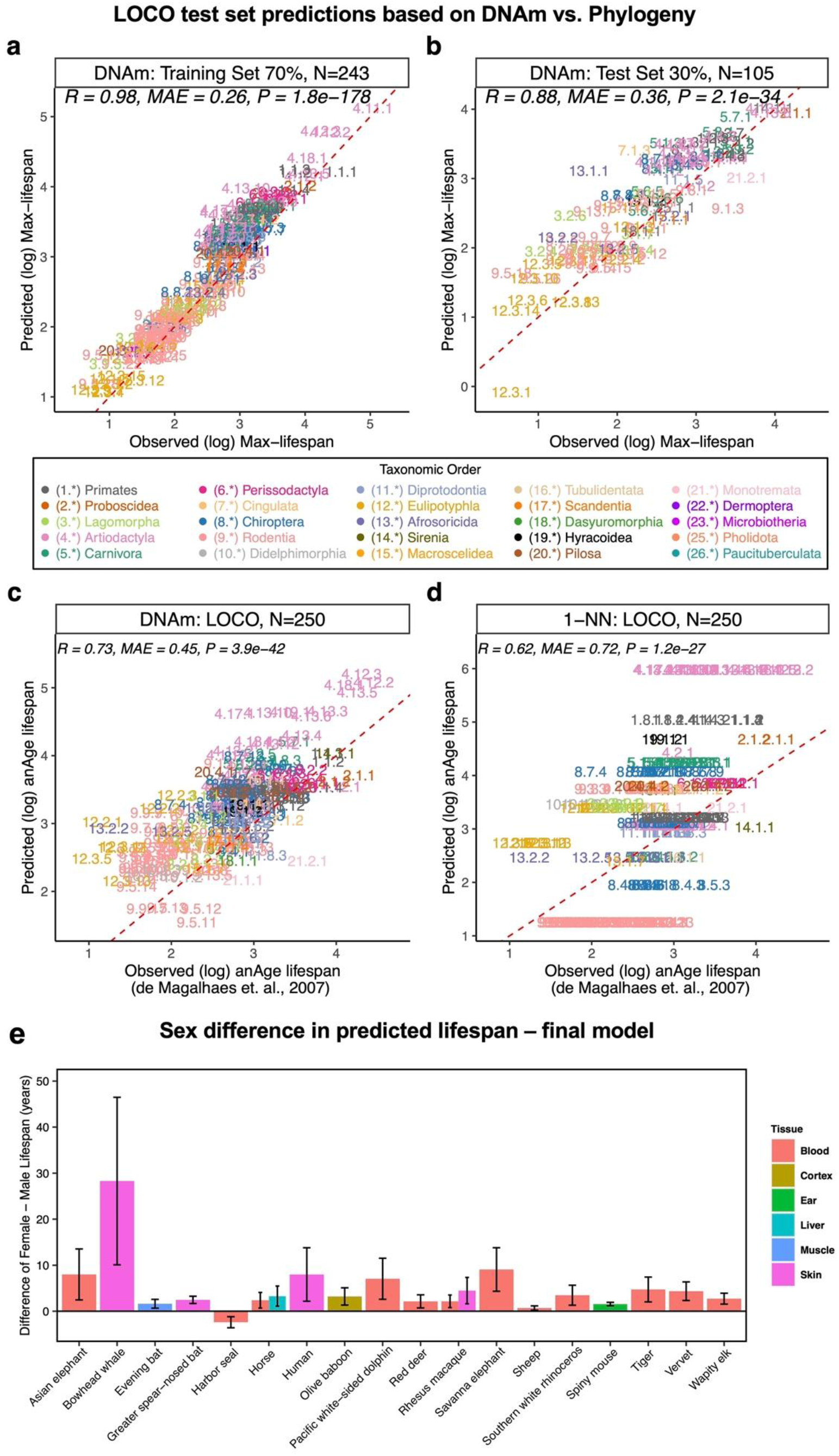
Comparison of DNAm lifespan predictor, phylogeny-based predictor, and sex-related differences in predicted lifespan. **a, b**, Evaluation of the multivariate predictor of maximum lifespan based on cytosine methylation in training data (panel **a**) and test data (panel **b**), encompassing 70% and 30% of species, respectively. In panels a and b, each data point symbolizes a unique species, differentiated by its taxonomic order color-coding. The dotted red line indicates the fitted linear regression. **c, d**, Leave-one-clade-out (LOCO) cross-validation analyses concentrate on the log-transformed (base e) maximum lifespan predictions. Given that several species’ missing lifespan observations were filled using neighboring species, lifespan estimates naturally favor k-NN. To mitigate this bias, this analysis only includes 250 species from the original anAge database (*2*) with actual maximum lifespan records. This analysis provides an unbiased assessment of the performance of the DNAm elastic net predictors (panel **c**) with the 1-Nearest-Neighbor (k-NN) predictor (panel **d**), which uses distances from the Mammalian phylogenetic TimeTree (*54*). **e**, Bar plots emphasize the differences in lifespan predictions between females and males, specifically highlighting species that exhibits uniformity across tissues with statistically interesting (two-sided unadjusted Wilcoxon rank sum test p-value =< 0.01) female-male differences. This means in all statistically interesting tissue groups, females are consistently predicted to have longer DNAm lifespan. Only species with DNAm lifespan predictions that significantly differ between sexes are reported, based on a two-sample T-test with a p-value less than 0.01. Error bars outline the 95% confidence interval of these differences. Bars throughout the figure are colored by tissue type, as detailed in the accompanying legend.

Shifting our focus to other life history traits, the actual log gestation time—which is inherently more straightforward to determine than maximum lifespan—manifested an even higher correlation with its predicted counterpart (R = 0.96, **Fig. 1c**). Intriguingly, the epigenetic prediction of (log-transformed) age at sexual maturity presented a somewhat lower correlation of R = 0.85 with recorded data (**Fig. 1d**). This discrepancy might stem from the fact that the age at sexual maturity is considerably more variable than gestation time, being influenced by factors like food availability and varying ecological conditions.

We will refer to the predicted maximum lifespan, expressed in log years, as either the epigenetic maximum lifespan or DNA methylation (DNAm) maximum lifespan. Analogous naming conventions will apply to other DNAm-derived estimates of life history traits. The final life history predictor coefficients, which were trained on all available samples, and the corresponding CpG annotations are summarized in **table S5-S7** (also available as an R package on Github: caeseriousli/MammalianMethylationPredictors).

### Chronological age versus epigenetic maximum lifespan

We carried out two analyses to study the relationship between the life history traits and chronological age of the individuals of species sampled. First, we built a separate maximum lifespan predictor using only samples obtained from animals that were younger than their species’ average age of sexual maturity and younger than 5 years, and this had a considerable correlation with predicted maximum lifespan (R = 0.68, Fig. S1), even though the restriction of age resulted in fewer species (n = 122) being available for this analysis. The predictor’s remarkable accuracy in long-lived species (for instance, those with a maximum lifespan exceeding 20 years) indicates that the determinants of maximum lifespan can be discerned from DNA samples obtained even from relatively young individuals.

Second, we utilized the finalized lifespan predictor model on individual animal samples. While the predictor was designed to estimate species-level lifespan on a logarithmic scale, we used the coefficients to predict the lifespan of individual samples. Our findings indicate that the predicted maximum lifespans for individual samples can vary and, in certain species such as the naked mole rat skin, human blood, sheep ear, and cat blood, correlate significantly with chronological age (**Fig. S2**). In a similar vein, gestation duration and age of sexual maturity correlate significantly with age in select species tissue strata (**Fig. S3, Fig. S4**).

**Fig. 3.**
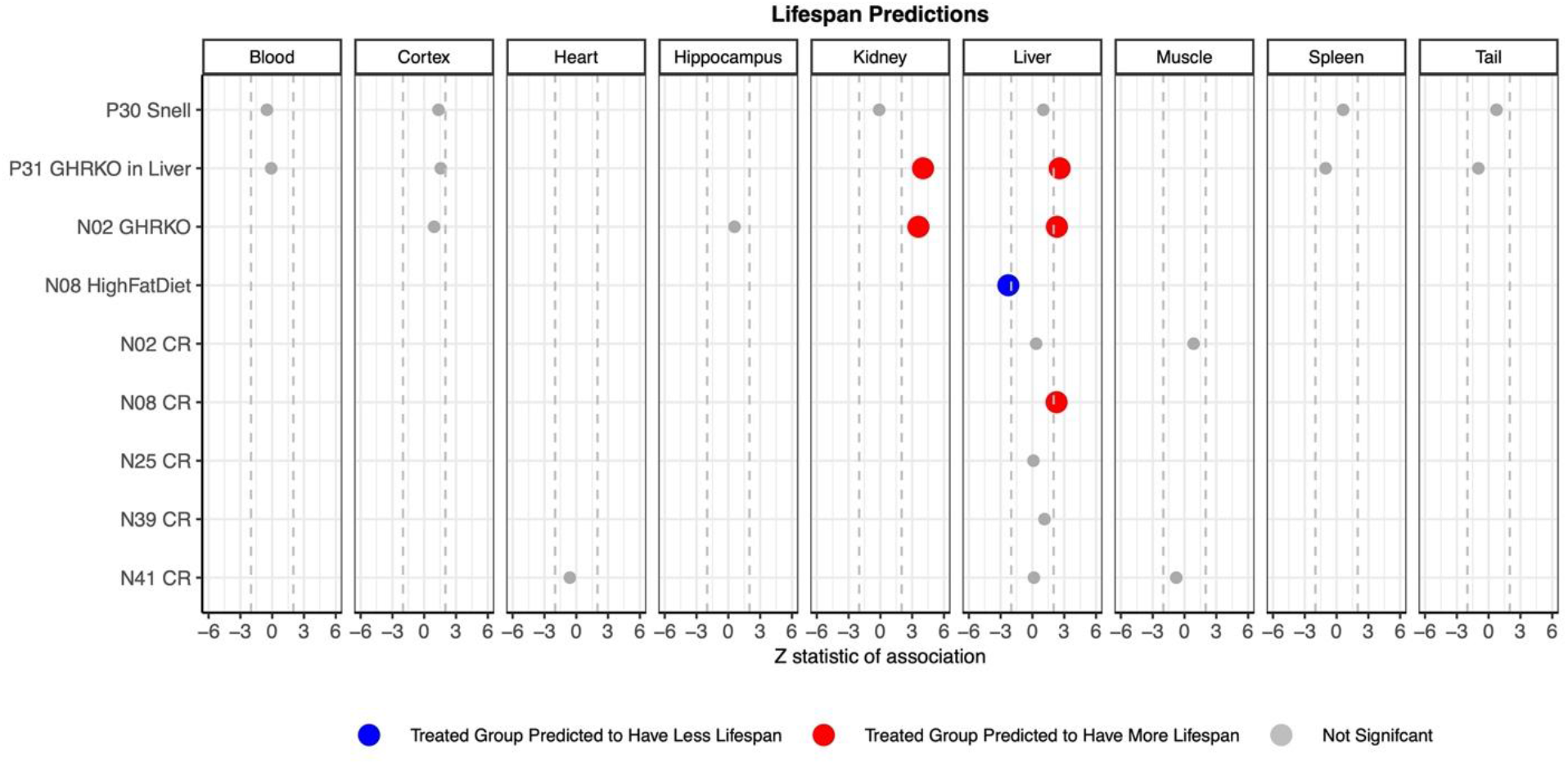
Predicted lifespan across murine experimental treatment groups. This figure presents the predicted lifespan from our final model based on murine perturbation experiments. Each row corresponds to a specific experiment, and columns stratify these results by tissue type. The experimental treatment groups, from top to bottom, are as follows: Snell Dwarf mice, liver-specific growth hormone knock-out mice, full-body growth hormone knock-out mice, high-fat diet, and five separate caloric restriction experiments. The prefixes in the rows, like P30 for “project 30” and N08 for “number 8”, denote distinct data sets. Empty cells signify the absence of samples for the corresponding tissue in the experiment. Grey dots represent associations that are not statistically significant. Red and blue markers highlight significant associations (p<0.05) that align with our expectations. We found no significant associations that deviated from our expectations. The x-axis reports Wald test statistics that follow a standard normal distribution under the null hypothesis. Dashed lines represent the critical Z statistic values when assessing a two-sided T-test with Type I error controlled at ALPHA=0.05.

**Fig. 4.**
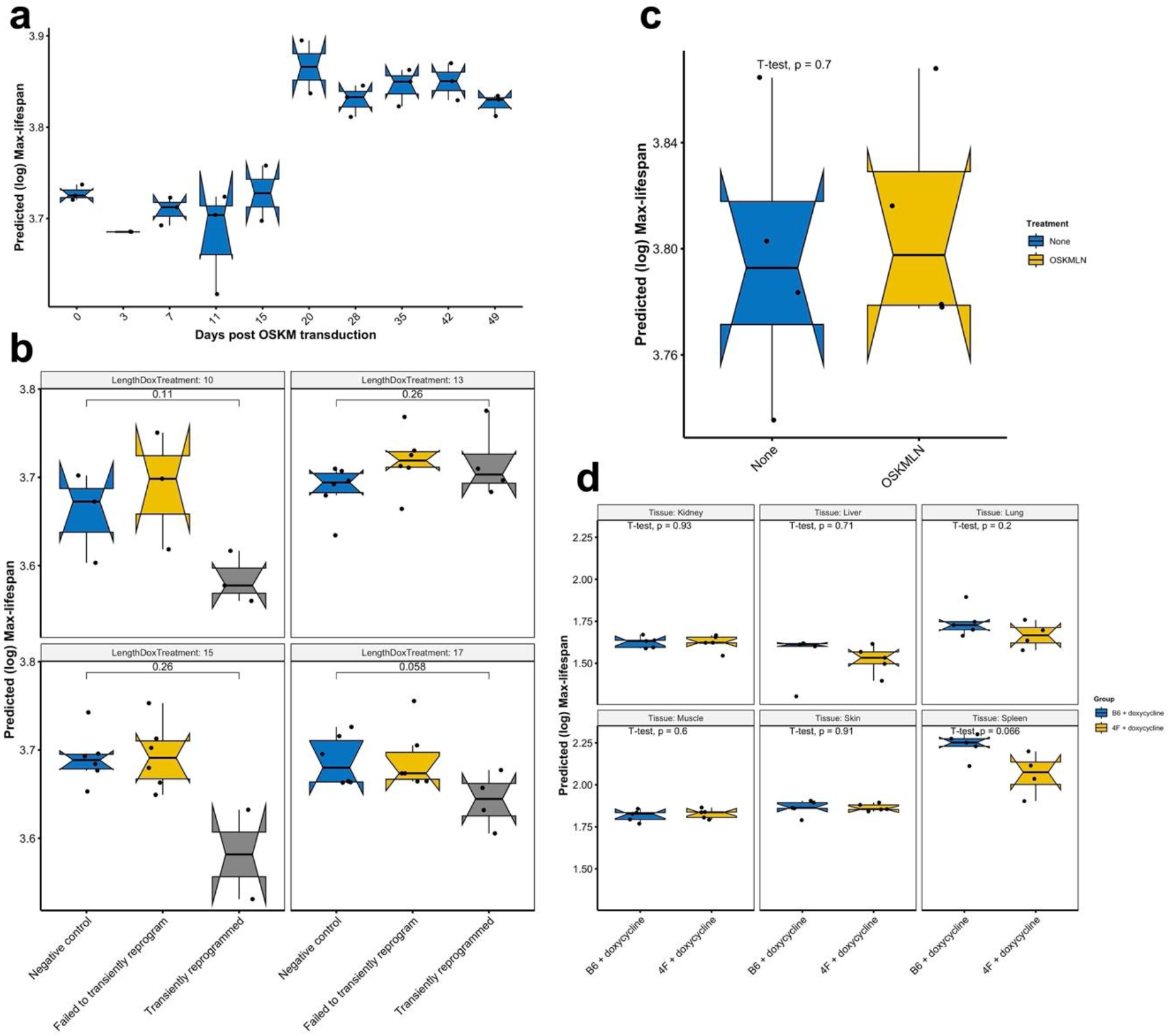
Partial or full OSKM reprogramming versus epigenetic maximum lifespan. Panels show, **a**. Predicted max lifespan in a 49-day full reprogramming time course of human dermal fibroblasts (HDFs) resulting in iPS cells (Kruskal Wallis test *p-value* = 0.0086) (*55*). Y axis: log(max lifespan) calculated from DNA methylation arrays from the following cell populations: day 0 (HDFs), day 3 (OSKM expressing EGFP (+) HDFs), day 7 to 28 (TRA-1–60 (+) cells at intermediate stages of reprogramming), and iPSCs after days 35. **b.** Predicted max lifespan of human dermal-fibroblasts (HDFs) after transient reprogramming (GSE165179) (*37*). Different lengths of transient reprogramming were separated into sub-panels. Negative control cells, transiently reprogrammed cells (CD13− SSEA4+) and cells that failed to transiently reprogram (CD13+ SSEA4−) were included in the plot. **c.** Predicted max lifespan of HDFs with transient expression of OSKMLN (GSE142439) (*56*). OSKMLN was daily transfected for 4 consecutive days, and DNA methylation was measured 2 days after the interruption. **d.** Predicted max lifespan in various tissues of 4F mice after 7 months of treatment (GSE190665) (*35*). B6 or 4F mice were given doxycycline in drinking water for 2 d followed by 5 d of withdrawal. The treatment started at 15 months of age and continued until 22 months of age (7- month treatment). B6 mice: WT mice; 4F mice: mice with the OSKM polycistronic cassette.

Overall, our analysis reveals that epigenetic indicators of life history traits, when confined to a specific species and tissue, do not have a consistent correlation with age.

### Tissue type can play a role

In the preceding section, we introduce epigenetic predictors for life history traits, derived from mean methylation levels averaged across species and encompassing all available tissue types. As these predictors disregarded specific tissue types, we term these as tissue-agnostic life-history predictors.

To delve deeper into the influence of tissue type on lifespan predictions, we applied our epigenetic predictors—specifically for maximum lifespan, gestation duration, and age of sexual maturity—to selected species with data from various tissues (**Fig. S5**, **Fig. S6**, and **Fig. S7**).

**Fig. 5.**
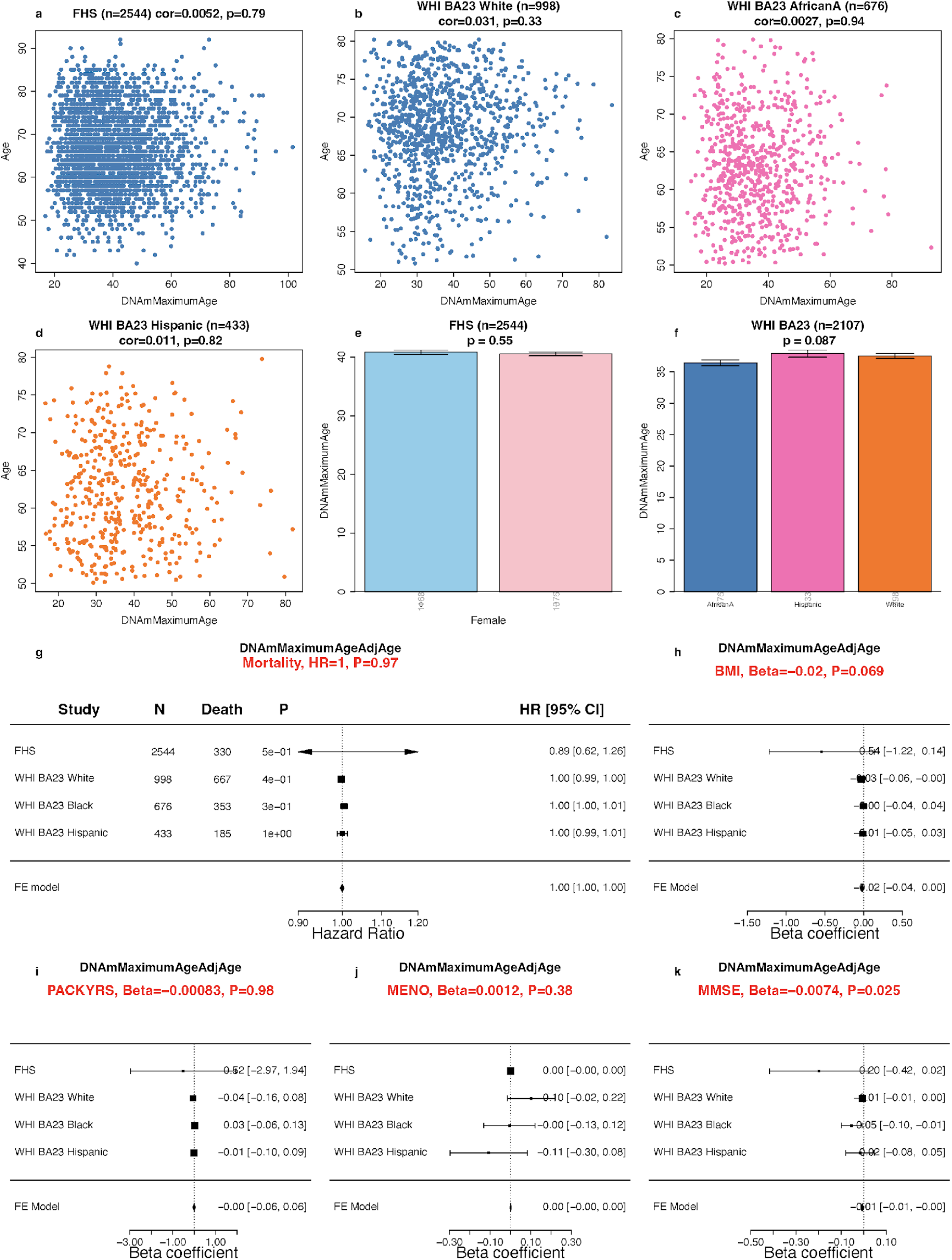
Methylation-based estimate of maximum lifespan in human cohorts. Panels **a-d** present scatter plots of the predicted maximum lifespan, transformed from log-years back to years (DNAmMaximumLifespan, x-axis), against chronological age (y-axis). These panels depict data from a) n=2,544 Caucasians of European ancestry in the Framingham Heart Study Offspring Cohort (FHS) and **b-d**) 2,107 women from the Women’s Health Initiative Cohort Broad Agency Award 23 (WHI BA23). This data is further categorized by three racial/ethnic groups: European ancestry, African American ancestry (AfricanA), and Hispanic ancestry. Each data point symbolizes an individual and is color-differentiated based on ethnic group. Titles indicate the sample size and furnish the Pearson correlation coefficients accompanied by their respective p-values. Panel **e** contrasts DNAm maximum lifespan with sex in the FHS, while **f** relates to ancestry. Panel **g** is a forest plot summarizing a meta-analysis of Cox regression models for time-to-death (due to all causes), based on the age-adjusted version of DNAmMaximumAge. This analysis spans various study-ethnic groups, with each row detailing the hazard ratio [95% confidence interval] for a one-year elevation in DNAmMaxLifespanAdjAge. The title reports the meta P-value, derived using inverse variance-weighted fixed-effect models. Forest plots showcase the correlation between age-adjusted DNAmMaxLifespan and the following variables: **h**, human body mass index (BMI), **i**, self-reported smoking pack-years, **j**, age at female menopause, **k**, Mini-Mental State Examination (MMSE) scores. The analysis, which spans different study-ethnic groups, outlines in each row the correlation coefficient [95% CI] corresponding to a one-year increase in DNAmMaxAgeAdjAge. All p-values are two-sided and are presented in their nominal form, without adjustment for multiple comparisons.

**Fig. 6.**
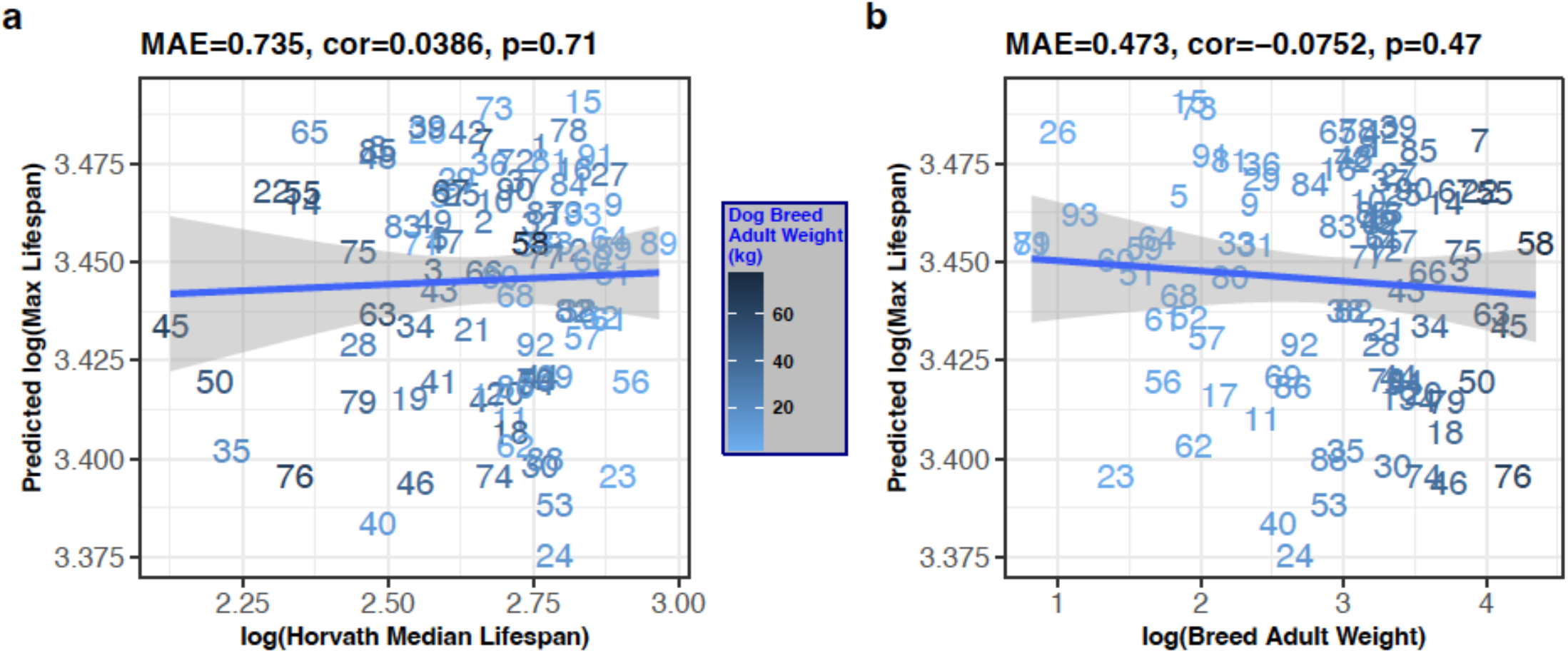
Mammalian lifespan prediction applied to blood methylation data from 90 different dog breeds. All quantities are log transformed (base e). Predicted log transformed maximum lifespan (y-axis) versus characteristics of dog breeds. **a.**, actual maximum lifespan of the breed (x-axis). Maximum age of the dog breed was estimated as the product of 1.33 times the median lifespan of the breed from Horvath et al 2022, (*41*). **b.,** Average adult weight of the dog breed. Each integer label corresponds to a different dog breed (*41*).

The epigenetic maximum lifespan estimates do reveal disparities between certain tissues. For instance, in human samples, distinct epigenetic lifespan predictions emerge (**table S3**): Blood and epidermis yield elevated lifespan predictions of 98.1 and 94.6 years, respectively, while skin and cerebral cortex produce estimates of 79.1 and 51.1 years, respectively. In contrast, embryonic stem cells (34.4 years), iPSC cells (25.6 years), endothelial cells (23.9 years), and skeletal muscle (35.4 years) present lower lifespan predictions (**table S3**).

Interestingly, the trend of blood samples reflecting the highest epigenetic maximum lifespan is consistent across various species (**Fig. S5**). For instance, in species ranging from humans to brown rats, blood samples consistently indicate elevated epigenetic maximum lifespan predictions.

In horses, we’ve observed that blood results in elevated lifespan predictions, whereas the ovaries and adrenal cortex yield lower estimates (**Fig. S5**). In mice, blood, LSK Progenitor Hematopoietic Stem cells, and bone marrow macrophages stand out with elevated predictions, whereas other tissues align closely. Both beluga whales and rhesus macaques show elevated lifespan estimates in blood (**Fig. S5**). In summary, blood samples consistently yield higher epigenetic maximum lifespan predictions across a variety of species. A detailed overview is available in **table S3**. The biological significance of these disparities warrants further investigation.

We briefly describe a strategy for building epigenetic predictors of life history traits that mitigate the confounding influence of tissue types. A predictor for maximum lifespan can be built based on mean methylation levels in strata formed by species and tissue type, termed tissue-aware life history predictors. In this setup, every species is represented through multiple data points corresponding to different tissues collected from the same species. Notably, these predictors, rooted in species-tissue aggregated data, are highly accurate (**Fig. S8**). In addition, maximum lifespan, gestation time, and age at sexual maturity predictors produce similar tissue-stratified results (**Fig. S9, Fig. S10, Fig. S11**). The predictor coefficients and corresponding CpG annotations are summarized in **table S8-S10**.

In our subsequent discussions and the remainder of the article, we will focus on tissue-agnostic predictors for life history traits.

### Superior Performance of DNAm-Based Predictors Over Phylogeny-Based Models

While DNA methylation levels are influenced by genetics, our DNAm-based lifespan predictor seems to transcend mere DNA sequence variation influenced by phylogenetic relationships. This assertion is supported by two distinct analyses.

First, we employed elastic net regression models to predict maximum lifespan using both CpG methylation data and taxonomic order indicators. Interestingly, the model exclusively selected CpGs, indicating their superior explanatory power over taxonomic variables in lifespan variation. Second, we compared the accuracy of the epigenetic lifespan predictor against k-nearest neighbor (k-NN) regression models, which base predictions on phylogenetic tree branch lengths. At its simplest, with K=1, the k-NN model predicts a species’ lifespan based on its closest taxonomic neighbor. Upon evaluating the correlation between predicted and actual values, the phylogeny-driven k-NN model slightly trails the DNAm predictor, especially under a leave-one-species-out evaluation. This is primarily because many mammalian species in our dataset exhibit lifespans akin to their taxonomic neighbors (**Fig. S12a,b, table S1**). This trend is also pronounced at the taxonomic family level (**Fig. S12c,d**). However, the k-NN model’s performance diminishes under a more rigorous leave-one-clade-out (LOCO) evaluation, which tests the model’s ability to predict lifespan of taxonomically diverse species. While k-NN models (with K=1) achieved a moderate correlation of R=0.62 (**Fig. 2d, Fig. S13**), they lag behind the methylation-based predictor, which boasts a correlation of R=0.73 (**Fig. 2c, Fig. S12**). k-NN models with K=2 and K=3 neighbors yielded correlations of R=0.62 and R=0.57, respectively. A detailed examination of the residuals highlights the k-NN model’s tendency to make generalized predictions for larger taxonomic orders, often deviating significantly from actual values (**Fig. S13**).

In conclusion, when assessed through LOCO cross-validation, DNAm-based predictors distinctly outshine their phylogeny-based counterparts. The DNAm predictor’s capability to accurately estimate lifespan across diverse taxonomic orders underscores its potential to capture aspects of mammalian lifespan that transcend mere phylogenetic relationships.

### Sex differences in predicted lifespan

We aimed to investigate any potential disparities in maximum lifespan predictions across sexes. Using our final regression model, based on average methylation data per species and designed to predict species-level lifespan on a logarithmic scale, we predicted individual sample lifespans. Predictions from female tissues showed a striking alignment with those from male tissues, with a strong correlation of R = 0.99 on a log scale. Most species showed consistent epigenetic estimates of maximum lifespan in female and male samples (**table S4**, column “Female – Male Significant Tissues,” where a “+” denotes Female minus male mean predicted DNAm lifespan is positive with an unadjusted p-value =< 0.01, “-” vice versa, and “.” denotes a p-value > 0.01). Stratifying by tissue type, we observed statistically interesting consistent sex difference in epigenetic maximum lifespan (a more conservative two-sided unadjusted Wilcoxon rank sum test p-value < 0.01) (*20*) in only 18 species (**Fig. 2e**). This means we only consider it statistically interesting when, 1. at least one tissue group within the species exhibits statistically interesting (two-sided unadjusted Wilcoxon rank sum test p-value =< 0.01) female-male differences 2. these differences must be in the same direction, for example, the female mean DNAm lifespan being greater than that of the males. In other words, we look for species for which one sex is consistently predicted to have longer DNAm lifespan than the other. Females were predicted to have a longer maximum lifespan than males in 17 of the 18 species, including humans (**Fig. 2e**, **table S4**). The one exception was blood from harbor seals. Across all species, females have a 1.8% longer predicted epigenetic maximum lifespan than males of the same species.

### Adult weight is not a driver of prediction accuracy

Across species, there is a notable correlation between maximum lifespan and average adult weight (body mass), as depicted in **Fig. S14a**. This correlation has been well-documented in prior studies (*2*). Given this, we evaluated whether the high accuracy of epigenetic lifespan predictors could be influenced by the average adult weight. Our findings from two distinct analyses suggest otherwise.

In the first analysis, we focused on small animals, specifically those with an average adult weight of less than 150 grams. Despite a negative correlation between adult weight and maximum lifespan in these species (R = -0.28, **Fig. S14c**), the epigenetic predictor of maximum lifespan still showed a strong correlation with observed values (Pearson correlation R = 0.43, P = 6.2×10-7, **Fig. S14b**). In the second analysis, encompassing all animals, a multivariate regression model (with the dependent variable being the log of maximum lifespan, indicated that (log transformed) adult weight (Wald test P = 1.3×10-6) is a less significant covariate than (log-transformed) epigenetic maximum lifespan (P < 2×10-16). This shows that adult weight only weakly mediates the effect of epigenetic maximum lifespan on actual maximum lifespan. This observation is reinforced by a correlation value (R) of 0.54 between our model’s predictions, after weight adjustments, and the actual maximum lifespan (**Fig. S14d**). In conclusion, both analyses consistently show that the epigenetic maximum lifespan provides predictive information that extends beyond adult weight.

### Cancer mortality risk across mammals

Distinct variations in cancer mortality rates across major mammalian orders have been documented (*21*). Notably, there exists a pronounced negative correlation between mammalian cancer risk and observed gestation time (Pearson r=-0.37, p=0.0031, **Fig. S15**). Considering the notable correlation among gestation time, maximum lifespan, and age at sexual maturity on a logarithmic scale (**Fig. S15a-b**), one might theorize that one or more of these life history traits could predict cancer mortality risk in mammals. However, this theory is challenged by the data: neither maximum lifespan (**Fig. S15d**) nor average age at sexual maturity (**Fig. S15e**) exhibits this anticipated relationship. The only significant correlation with cancer mortality risk is observed for gestation time and its epigenetic counterpart (r=-0.41, p=0.00092, **Fig. S15i**). Further, upon adjusting for observed values, no significant correlation was found between epigenetic predictions of life history traits and mammalian cancer risk (**Fig. S15k-m**). Collectively, these findings indicate that the epigenetic markers predicting life history traits, such as gestation time, do not inherently offer predictive information into mammalian cancer risk beyond the observed life history values. This result is consistent with the concept of Peto’s paradox where there is no correlation between cancer rates and either maximum lifespan or body mass (*21*).

### Weak effect of mutations in the somatotropic axis

The somatotropic axis, encompassing growth hormone, IGF-1 levels, and their respective receptors, is a focal point in aging and longevity research (*22*). Growth hormone receptor knock-out mice (dwarf mice) typically exhibit an extended maximum lifespan (*23, 24*). Intriguingly, a full-body growth hormone receptor knock-out (GHRKO) mouse holds the record of nearly reaching a lifespan of five years (*22*). In our study, we sought to determine if decreased GH/IGF-1 pathway activity influences the epigenetic estimates of maximum lifespan across three distinct mouse models. It should be noted that Snell dwarf mice and full-body GHRKO mice show extended maximum lifespans (*25-27*). On the other hand, liver-specific GHRKO mice, despite exhibiting reduced serum IGF1 levels, do not show a corresponding increase in maximum lifespan (*28, 29*).

Our observations indicate that both the full-body GHRKO and liver-specific dwarf mice show a notably extended epigenetic maximum lifespan, particularly in samples from liver and kidney (**Fig. 3**). However, such association was not observed in samples from blood, cerebral cortex, hippocampus, spleen, or tail. Similarly, we did not detect any significant association across tissues in Snell dwarf mice. Given these observations, two potential inferences emerge. Either manipulation within the somatotropic axis (comprising growth hormone, IGF-1 levels, and their associated receptors) has only a weak effect on epigenetic lifespan estimators in select tissues, or the epigenetic predictor of maximum lifespan is insufficiently precise when utilized in mouse studies.

### Equivocal effect of caloric restriction and high-fat diet on epigenetic lifespan

Caloric restriction has been documented to extend the maximum lifespan in approximately one-third of all mouse strains. We aimed to gauge the influence of caloric restriction on the epigenetic estimates of maximum lifespan from mouse liver samples. Surprisingly, in four of the five studies, no significant (when assessed with a relaxed, unadjusted Type I error rate control of 5%) impact on epigenetic maximum lifespan in murine liver was observed (**Fig. 3**). Only one study presented the expected association between caloric restriction and a prolonged maximum lifespan (**Fig. 3**).

On the other hand, high-fat diets have been identified as factors that both shorten murine lifespan and accelerate epigenetic aging (*30*). Consistent with this, our observations did confirm the anticipated link between a high-fat diet and a reduction in epigenetic maximum lifespan (**Fig. 3**). In sum, the outcomes from the application of epigenetic maximum lifespan indicators to mouse interventions, which inherently influence mouse longevity, are somewhat equivocal.

### Cellular reprogramming based on the Yamanaka factors

The Yamanaka factors, comprising Oct4, Sox2, KLF4, and Myc, are known for their role in full reprogramming (resulting in induced pluripotent stem cells) as well as in partial reprogramming of somatic cells (*31-36*). We tested whether reprogramming affects epigenetic maximum lifespan using publicly accessible data from both complete and partial reprogramming studies conducted on human and mouse cells.

Our findings (**Fig. 4**) show the maximum lifespan predictor outcomes for various cellular reprogramming treatment groups. Notably, human dermal fibroblasts subjected to a full reprogramming course based on OSKM transduction exhibited a slightly increased (and statistically significant p<0.05) epigenetic maximum lifespan after 20 days (see **Fig. 4a**). Meanwhile, in a partial reprogramming experiment (GSE165179) (*37*), the treatment group displayed a marginally reduced mean predicted maximum lifespan. However, the disparity between the groups did not reach a statistically significant level (**Fig. 4b**).

We note that our examination of tissue and cell types did not yield conclusive evidence indicating a significant divergence in the epigenetic maximum lifespan between embryonic stem cells or iPS cells and primary cells (see **fig. S5**). Our findings are somewhat inconclusive. Although full reprogramming in human dermal fibroblasts hints at an increase in epigenetic maximum lifespan after 20 days of OSKM administration, other experiments were unable to confirm this effect. We discuss caveats surrounding the measurement platform below.

### Human epidemiological cohort studies

We utilized methylation-based estimators to assess the maximum lifespan in blood samples sourced from participants of the Framingham Heart Study (n=2544) (*38*)and the Women’s Health Initiative (n=2107) (*39, 40*). Given that these samples were processed using a different methylation platform (the human Infinium 450K array), we employed the Array Converter software to convert values from the mammalian methylation probes (*18*). We observed no significant correlations between the predicted maximum lifespan and the actual age of participants across three distinct racial/ethnic groups (**Fig. 5a-d**). It’s important to highlight that this finding contrasts with our previous analysis, where we identified a correlation between age and epigenetic maximum lifespan in humans. These discrepancies likely arise from variations in measurement platforms. Our earlier analyses employed the mammalian array, whereas the epidemiological cohort studies utilized the human Illumina array.

Our analysis reveals no significant associations with other demographic variables: in blood samples, the DNAm-based maximum lifespan does not show a significant association with sex (p=0.55, **Fig. 5e**), racial/ethnic group (p=0.087, **Fig. 5f**), human mortality risk (**Fig. 5g**), body mass index (p=0.069), smoking pack years (**Fig. 5i**), or age at female menopause (**Fig. 5j**). The Mini-Mental State Examination (MMSE) is a diagnostic tool for cognitive impairment and dementia. The MMSE evaluates various cognitive domains. A higher score on the MMSE indicates better cognitive functioning. We noted a marginally significant negative correlation (p=0.025, **Fig. 5k**) between MMSE and age-adjusted DNAm lifespan. However, this significance disappears after accounting for multiple comparisons.

We delved into the relationship between our methylation-based lifespan estimators and several dietary and health-related biomarkers (**Fig. S16**). This comprehensive assessment covered 59 variables: 27 from self-reported dietary inputs, 9 from blood-based dietary measurements (including mean carotenoid levels, indicative of vegetable and fruit consumption), 17 clinical indicators including metabolic characteristics, central adiposity, inflammatory markers, leukocyte telomere length, cognitive performance, lung function. We also analyzed lifestyle and demographic variables (diet, exercise, education, income). Upon analysis, neither the epigenetic estimate of maximum lifespan nor its age-adjusted counterparts showed any significant association with the biomarkers after adjusting the analysis for multiple comparisons (**Fig. S16**). The inconclusive results suggest that lifestyle behaviors do not profoundly influence the maximum bounds of human lifespan, as measured by epigenetic predictors. However, it is essential to highlight a significant limitation in our analysis: the human data was sourced from a different methylation array platform and was heavily dependent on imputation methods. Future research should revisit these findings using data from methylation platforms that assess the highly conserved CpGs on the mammalian array.

### Evaluation in different dog breeds

Dog breeds display a remarkable variability in lifespan, with certain breeds outliving others by as much as two-fold. We assessed our epigenetic predictor of maximum lifespan using 742 individual blood samples sourced from 93 distinct dog breeds (*41*). However, our canine dataset presented two primary challenges. First, the representation of dogs within each breed was inconsistent, ranging from as few as 2 samples for the English Setter to as many as 95 for the Portuguese Water Dog. Second, there was a disparity in age distributions across breeds; for example, the relative ages R for the Otterhound breed spanned from 0.06 to 0.14, while for the Beagle, R ranged from 0.06 to 0.73 (*41*). To average out these inconsistencies, we took the average of maximum lifespan predictions for each breed. When applying the mammalian maximum lifespan predictor to blood samples from 90 diverse dog breeds (**Fig. 6**), we did not observe a significant correlation between the predicted mammalian maximum lifespan and either the breed’s average/maximum lifespan or its average weight. Overall, these results suggest that the epigenetic predictor of mammalian lifespan is not effective in predicting breed-specific lifespans in dogs.

Since the predicted maximum lifespan for various dog breeds showed only minor variation (**Fig. 6**), these results suggest that epigenetic maximum lifespan across dog breeds aligns closely with that of wolves. Thus, epigenetic maximum lifespan predictor remains unaffected by recent changes in dogs due to human selection and continues to reflect the lifespan of ancestral dog species.

## DISCUSSION

Drawing from the comprehensive dataset of our Mammalian Methylation Consortium, we developed multivariate predictors that adeptly discern maximum lifespan and associated life history traits. Notably, our epigenetic estimator demonstrated heightened precision for gestation duration (R = 0.96) compared to maximum lifespan (R = 0.89). This discrepancy might be attributed to the inherent challenges in procuring accurate maximum lifespan data across a diverse array of species.

In terms of sexual dimorphism in lifespan predictions, for the majority of species, there was a congruence in the predicted maximum lifespan between sexes. However, a distinct trend emerged in 17 species, including humans, where females displayed a longer predicted lifespan, with harbor seals being a notable exception. This observation resonates with previously published studies that underscores the longevity advantage of females (*42-44*).

Our epigenetic markers’ predictive prowess seems to transcend mere phylogenetic correlations, indicating their broader applicability. Interestingly, neither chronological age nor typical adult weight appeared to markedly sway the accuracy of our life history trait predictors. In numerous species, there was a conspicuous absence of correlation between chronological age and the epigenetic lifespan.

The actual maximum lifespan of humans, at 122.5 years, exceeds our epigenetic maximum lifespan estimates. For humans, the highest epigenetic lifespan values were observed in blood and epidermis samples, at 98.1 and 94.6 years respectively. This trend of elevated epigenetic lifespan in blood samples is consistent across various species, from humans to brown rats. We did not find definitive evidence suggesting that the epigenetic maximum lifespan of embryonic stem cells or iPS cells significantly diverges from that of somatic cells. The biological significance of cell type and tissue-specific variations in epigenetic lifespan predictions warrants further investigation.

Epigenetic maximum lifespan showed little variation across dog breeds indicating that it is not affected by recent genetic selection enforced on dogs and may represent the ancestral state of the dog as a species.

An independent analysis centered on CpG density revealed a compelling linkage with maximum lifespan (*10*). This underscores the combined importance of sequence information and methylation levels in predicting maximum lifespan.

Analysis of murine lifespan interventions showed that only growth hormone knockouts showed extended epigenetic lifespan in liver and kidney tissues, while other tissues and long-lived strains did not influence epigenetic maximum lifespan. strains. Similarly, caloric restriction did not affect epigenetic maximum lifespan.

Our analysis of human cohorts, despite its comprehensiveness, did not definitively determine the effects of lifestyle on epigenetic maximum lifespan. One possible constraint might arise from using different methylation array platforms for data gathering (specifically, the human Illumina array as opposed to the mammalian methylation array). For more accurate insights in future human epidemiological cohort studies, it would be beneficial to profile the highly conserved CpGs using the mammalian methylation array.

Taken together our results suggest that species maximum lifespan is determined, in part, by epigenetic signature that is largely independent of sex, body mass, calorie restriction, or other lifestyle factors. This signature may be an intrinsic property of each species that is difficult to change. Interestingly, only growth hormone knockout and full reprogramming had some effect on epigenetic maximum lifespan. It would be interesting to identify novel interventions that affect epigenetic maximum lifespan as they may be the key to achieving large lifespan differences observed between species.

## METHODS

### DNA methylation data

We used existing data from our Mammalian Methylation Consortium that were published previously (*18*). All data were generated using the mammalian methylation array (HorvathMammalMethylChip40) (*16*) which provides high sequencing depth of highly conserved CpGs in mammals. Nearly 36k probes (cytosines) on the array exhibit high levels of sequence conservation within mammalian species (*16*). The subset of species for which each probe is expected to work is provided in the chip manifest file which can be found at the NCBI Gene Expression Omnibus (GEO) as platform GPL28271, and on our Github webpage. The SeSaMe normalization method was used to define beta values for each probe and to calculate detection p-values (*45*).

### Data description

We analyzed methylation data from 348 mammalian species representing 25 out of 26 taxonomic orders (**table S2, Fig. 1**). The only order not represented was the marsupial order Peramelemorphia. DNA was derived from 59 different tissues and organs including blood, skin, liver, muscle, and brain regions (**table S1**).

### Life history traits and AnAge database

The high accuracy of the epigenetic estimator of maximum lifespan is a testament to the success of a decade-long effort of biologists and the anAge database (*2*) to establish this elusive phenotype. For several species, maximum lifespan was not available in anAge. For select species, we used a K=1 nearest neighbor predictor to impute values. Therefore, we limited our comparative analysis to species for which this value was available and did not require imputation. To enhance the reproducibility of our findings we include our updated version of the anAge database (*2*) (**table S1**).

### Multivariate estimators of maximum lifespan

The regression coefficients from the final predictor, that is, the full model trained on all available species-level data for extrapolation purposes, are reported in **table S5**. For most species, relatively few animals informed the determination of maximum lifespan, which may bias this life history trait (*46, 47*). To account for the fact that the maximum lifespan of humans and mice was established based on many studies while the maximum lifespan of other mammalian species was based on fewer animals, we corrected the maximum lifespan value of the remaining species by multiplying it by 1.3. This adjustment step assumes that each maximum lifespan estimate reported in anAge underestimates the true value by 30 percent in all species except for humans and mice. We applied the same adjustment step in our universal mammalian clock project (*18*). In addition, in the final model fitted to all species as a training set, we calibrated the predictor by the mean and standard deviation, similar to those of biomarker, to match those of the observed lifespan (*48*).

The empirical distributions of mammalian life history traits in general, as well as they are represented in our dataset, are highly skewed towards the larger values. This is due to the fact that few species (such as humans and bowhead whales) live much longer than the majority of mammal species. Therefore, to maintain statistical integrity regarding regression model fitting, and to counteract uneven species weighting by the numerous short-lived species, we employed a log-transformation step to all three life history traits studied here.

We used elastic net regression to build different multivariate predictors of maximum lifespan, gestation time, and age at sexual maturity (*49*). To build a model on the basis of CpGs that are present/detectable in most species, we restricted the analysis to CpGs with significant median detection p-values (false discovery rate<0.05) (*50*) in 85% of the species. This resulted in a lower-dimensional dataset consisting of 17,032 CpGs.

We employed three strategies for building maximum lifespan predictors. The first strategy ignored tissue type. Here, all tissue samples from a given species were averaged resulting in a single observation per species. To arrive at unbiased estimates of the predictive accuracy of lifespan and other predictors, we used a leave-one-species-out (LOSO) cross-validation analysis that iteratively trained the predictive model on all but one species. Next, the predictor was applied to the observations from the left-out species. By cycling through the species, we arrived at LOSO estimates for each species. The second strategy formed average values for each stratum defined by tissue type and species. For example, this analysis formed an average value for human blood (considered as one stratum). The second approach allowed us to study the influence of tissue type on lifespan predictions. This second strategy shows similar prediction correlations in all three life history traits (**Fig. S8**).

Third, we also conducted a leave-one-clade-out analysis as described in the following. Conducting a comprehensive leave-one-taxonomic order out cross validation presented challenges. The primary issue was the unequal distribution of animals across taxonomic orders; for instance, Rodentia comprised 27% of all species, while many orders had fewer than 3% (table S2). To circumvent this, we adjusted the leave-one-order-out analysis. In larger taxonomic orders with over 20 species (like Rodentia, Artiodactyla, Chiroptera, Primates, Carnivora, and Eulipotyphla), we left out all species except two, representing the minimum and maximum lifespan. These two species functioned as a benchmark, tasking the predictor to estimate the lifespan for the entire taxonomic order based on limited data. Conversely, smaller taxonomic orders were left out completely as test sets. For instance, orders such as Dasyuromorphia, Microbiotheria, Sirenia, and Tubulidentata were represented only by a single species (table S2). This modified approach was termed the leave-one-clade-out (LOCO) analysis. A predictor heavily influenced by neighboring species with close lifespans, like the tree-based k-NN, would likely struggle with this methodology. Notably, as we used k-NN for imputing missing lifespan observations for several species, lifespan estimates naturally favor k-NN. Therefore, for this specific analysis, we relied on the original anAge database (*2*) that was devoid of imputed values.

It became clear that, while the k-NN lifespan predictor showed a reasonable prediction correlation, it frequently provided static and deviant predictions for entire taxonomic orders (**Fig. 2b**). When faced with any test set, the algorithm often perceived the “nearest” species as the two specified in the LOCO training set, or occasionally species in a neighboring small order. This led to uniform estimates across a taxonomic order, making the algorithm less effective for diverse species or clades.

For assessing the sex difference in individuals’ DNAm maximum lifespan prediction results, we chose to conduct two-sided Wilcoxon rank sum tests (*20*) instead of Student T-tests, for the following considerations, 1. small sample sizes in some species’ tissue-sex strata, 2. weak normality assumption in these small sample sizes, 3. Wilcoxon rank sum test is a relatively more conservative test than a Student T-test (*51*), 4. both Wilcoxon rank sum test and Student T-test work in other strata in which normality can be assumed and larger sample sizes are present, and 5. to be consistent across all strata and species, Wilcoxon rank sum test was used for sex difference in DNAm lifespan predictions.

### Interventions in mice

We used existing mammalian methylation data from mouse studies (*18*). The mammalian array data were generated using two versions of the mammalian array: the original mammalian array (called “40K” array) and the expanded array (referred to as “320K”) that also includes mouse probes (*16*). Some CpG probes unique to each array required imputation. Methylation levels of CpG sites missing on the 320K array were imputed from median beta values of the training mouse samples (“40K” array). None of the samples from the murine anti-aging studies were incorporated into the training set. Our DNAmMaxAge was assessed using the following independent test datasets: 1) Snell dwarf mice (n=95), 2) GHRKO experiment 1 (n=71), 3) GHRKO experiment 2 (n=96), 4) Calorie restriction (n=95).

T-tests evaluated whether these conditions affected epigenetic maximum lifespan. The DNA methylation data from datasets (1) and (3) were collected using an Illumina 320k customized array (available in GSE223943 and GSE223944). Datasets (2, 4, and 5) are available at GSE223748. Below is a brief overview of the experiments. Comprehensive details can be found in the Supplementary Information of (*18*).

Snell Dwarf Experiment (n=95): We analyzed tissues from 47 Snell dwarfs and 48 age-matched wild-type control mice, aged around 6 months. Snell dwarf mice, known for an approximately 30- 40% extended lifespan, lack growth hormone, thyroid-stimulating hormone, and prolactin. Methylation profiling was conducted on blood, cerebral cortex, liver, kidney, spleen, and tail from these mice.

GHRKO Experiments: We analyzed tissues from full body (n=71) and liver-specific (n=96) GHRKO studies. The full-body GHRKO mice exhibited prolonged lifespan, while liver-specific GHRKO did not. DNA methylation profiles were created for various tissues, and age-matching was performed.

Calorie Restriction Study (n=95): This study involved analyzing liver samples from 95 male mice, 59 from the calorie-restricted group and 36 controls. All mice, sourced from UT Southwestern Medical Center, Dallas, were 1.57 years old and from the C57BL/6J strain.

### Cancer risk in different mammals

We sourced estimates of mammalian cancer risk from a recent study (*21*). Two key metrics were considered: First, cancer Mortality Risk (abbreviated as CMR) - This refers to the ratio of cancer-related deaths to the total number of individuals for whom postmortem pathological records exist. It is a measure that has been used in various comparative studies (*52, 53*). Second, cumulative Incidence of Cancer Mortality (abbreviated as ICM) - This metric gauges the risk of cancer mortality by eliminating potential biases from both left and right censoring. Notably, there is a strong correlation between CMR and ICM, with a Pearson correlation coefficient of r = 0.89 (*21*). However, neither of these metrics showed any correlation with epigenetic maximum lifespan.

### Mortality analysis in human epidemiological cohort studies

We estimated DNAm maximum age in blood methylation data from 4,651 individuals from (a) the Framingham Heart Study (FHS) offspring cohort (n=2544 Caucasians, 54% women) (*38*) and (b) Women’s Health Initiative cohort (*39, 40*) (WHI, n=2,107, 100% women). Since these data were generated on a different platform (the Ilumina 450K array), we applied the Array Converter algorithm to impute mammalian methylation array data (*18*). Although the epigenetic maximum lifespan estimates are not correlated with chronological age, we defined a measure of epigenetic age acceleration (AgeAccel) as the raw residual resulting from regressing DNAm maximum lifespan on chronological age. By definition, the resulting DNAmMaxLifespanAdjAge measure is not correlated with chronological age. We applied Cox regression analysis for time-to-death (as a dependent variable) to assess if individual variation in the predicted maximum lifespan is attributable to mortality risk. The analysis was adjusted for age at blood draw and sex in the FHS cohort. We stratified the WHI cohort by ethnic/racial groups and combined a total of four results across the FHS and WHI cohorts using fixed effect models weighted by inverse variance. The meta-analysis was performed using the metafor function in R.

### Dog breeds

We used existing methylation profiles from 742 blood samples, representing 93 distinct dog breeds (Canis lupus familiaris) (*41*). Breed weight and average lifespan data were compiled from multiple sources as outlined in (*41*). We formed consensus values by integrating information from the American Kennel Club and the Atlas of Dog Breeds of the World. Lifespan approximations were derived from averaging standard breed lifespans across sexes. This information was gathered from a myriad of publications, most of which are multibreed studies focusing on age and mortality causes from veterinary clinics, as well as extensive breed-specific research typically conducted by purebred dog associations. The specific sources for each breed’s median lifespan are cited in (*41*).

To derive a reliable estimate for the maximum lifespan of each breed, we based our calculations on the breed’s median lifespan. Specifically, we used the formula: MaxLifespan = 1.33 x MedianLifespan. Notably, our conclusions hold even when applying different multipliers beyond 1.33, as the log transformation converts these multipliers into additive shifts. Comprehensive data on the breeds can be found in (*41*). Among the 93 breeds studied, median lifespans varied between 6.3 years (Great Dane, with an average adult weight of 64 kg) and 14.6 years (Toy Poodle, average adult weight being 2.3 kg).

### Data Availability Statement

The data have been made publicly available on Gene Expression Omnibus (GSE223748) as part of the data release from the Mammalian Methylation Consortium (*18*).

The mammalian methylation array is available from the nonprofit Epigenetic Clock Development Foundation https://clockfoundation.org/MammalianMethylationConsortium/.

Further, data sets are available on Gene Expression Omnibus (GEO) accession numbers, GSE174758, GSE184211, GSE184213, GSE184215, GSE184216, GSE184218, GSE184220, GSE184221, GSE184224, GSE190660, GSE190661, GSE190662, GSE190663, GSE190664, GSE174544, GSE190665, GSE174767, GSE184222, GSE184223, GSE174777, GSE174778, GSE173330, GSE164127, GSE147002, GSE147003, GSE147004.

#### Software Availability Statement

DNAm Predictor R package available at Github: caeseriousli/MammalianMethylationPredictors)

## ACKNOWLEDGEMENTS and FUNDING

We acknowledge our co-authors from the Mammalian Methylation Consortium (*18*).

This work was mainly supported by the Paul G. Allen Frontiers Group (SH) and Open Philanthropy (SH), by grants from the National Institute on Aging to AS, VG, and VNG, by the Milky Way Foundation to VG and Michael Antonov Foundation to VNG and VG. Tissue samples were contributed by the Museum of Southwestern Biology (MSB), University of New Mexico, Museum of Vertebrate Zoology (MVZ) and UC Berkeley which are linked by the Arctos Museum database (https://arctosdb.org/). We acknowledge the MVZ and Chris J. Conroy from the University of California, Berkeley for their contributions of some tissue samples (sample IDs and catalog IDs listed in **table S11**).

The Framingham Heart Study is funded by National Institutes of Health contract N01-HC-25195 and HHSN268201500001I. The laboratory work for this investigation was funded by the Division of Intramural Research, National Heart, Lung, and Blood Institute, National Institutes of Health. The analytical component of this project was funded by the Division of Intramural Research, National Heart, Lung, and Blood Institute, and the Center for Information Technology, National Institutes of Health, Bethesda, MD.

The Women’s Health Initiative program is funded by the National Heart, Lung, and Blood Institute, National Institutes of Health, U.S. Department of Health and Human Services through contracts HHSN268201600018C, HHSN268201600001C, HHSN268201600002C, HHSN268201600003C, and HHSN268201600004C. The authors thank the WHI investigators and staff for their dedication, and the study participants for making the program possible. A full listing of WHI investigators can be found at: chrome-extension://efaidnbmnnnibpcajpcglclefindmkaj/https://www.whi.org/doc/WHI-Investigator-Long-List.pdf. The views expressed in this manuscript are those of the authors and do not necessarily represent the views of funding bodies such as the National Heart, Lung, and Blood Institute; the National Institutes of Health; or the U.S. Department of Health and Human Services.

## CONTRIBUTIONS

Caesar Li developed the multivariate predictors of life history traits clocks. Amin Haghani, Ake T Lu helped with additional bioinformatics analyses. CZL, AH, SH, KR, VG drafted the first version of the article. All authors helped with editing the article and data interpretation.

## COMPETING INTERESTS

The Regents of the University of California filed a patent application (publication number WO2020150705) related to the mammalian methylation array for which Steve Horvath, and Jason Ernst are named inventors. SH is a founder of the non-profit Epigenetic Clock Development Foundation, which has licensed several patents from UC Regents, and distributes the mammalian methylation array. The remaining authors declare no competing interests.

## Supplementary Information

**Fig. S1.**
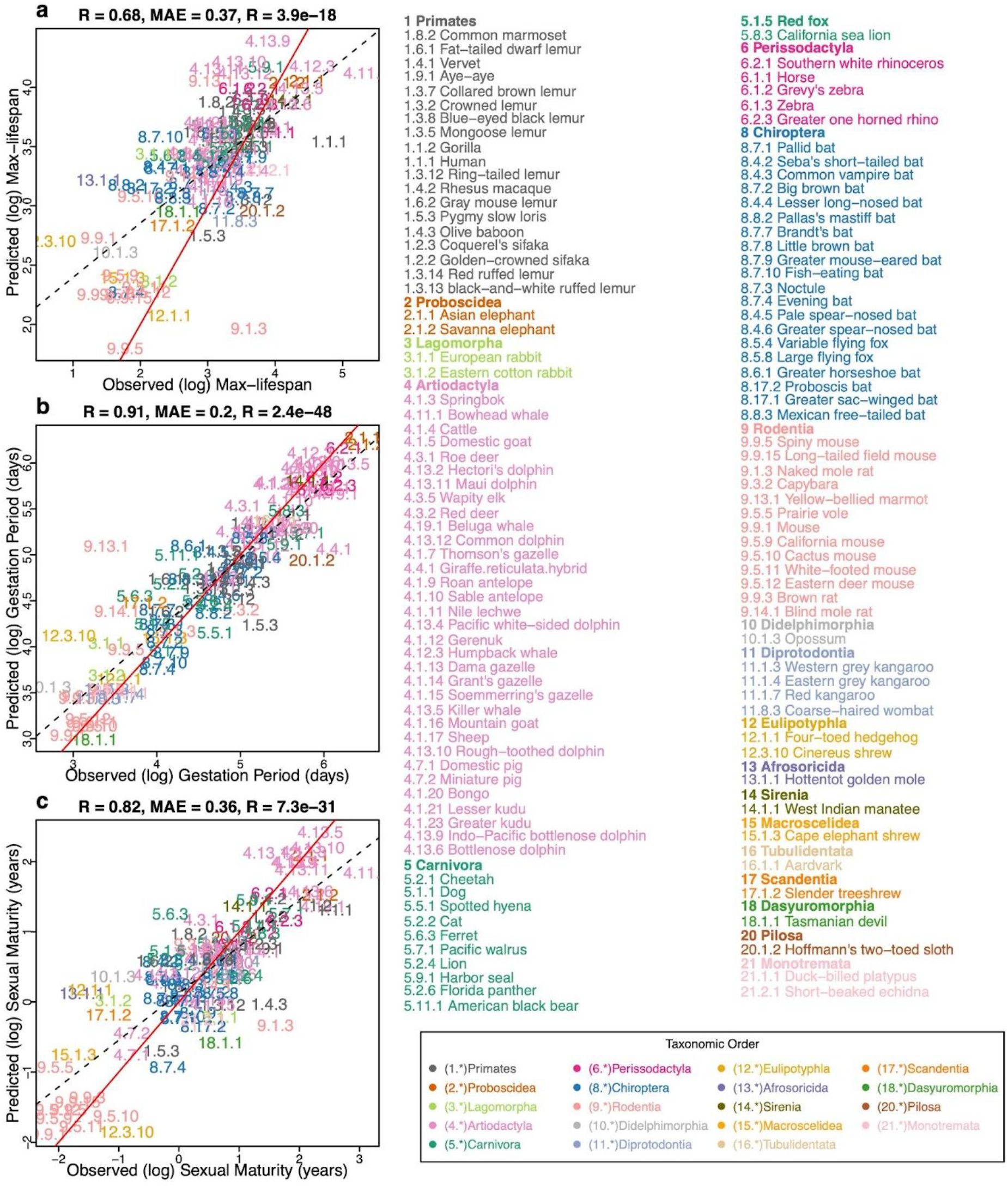
ElasticNet predictor based on young samples. Elastic Net Predictor, Leave-one-species-out analysis, fitted on a subset of all young samples (species n = 122). Young samples are defined as samples whose age is both younger than five years and less than the species’ average age at sexual maturation. Feature filtering and Elastic Net tuning parameter set-up is the same as those for **Fig. 1**. Three panels show predictors for **a,** log maximum lifespan (in log years), **b,** log-transformed gestation time (in log days), and **c,** log-transformed age at sexual maturity (in log years). As with the **Fig. 1**, species appear as designated numbers in scatter plot panels; the corresponding common names and phylogenetic orders are annotated in Figure legends; as indicated by the taxonomic order legend, the whole number (number before the decimal separator) part of each mammalian number is assigned in accordance with the corresponding taxonomic order. MAE abbreviates median absolute errors from the regression errors; r and p denote Pearson’s correlation and p-values, respectively. Numbers and colors are the mammalian species number and order annotation consistent with those of other Figures. Numeric values can be found in **table S1**. Red solid line represents the perfect prediction line, and the dotted line represents the fitted linear regression line.

**Fig. S2.**
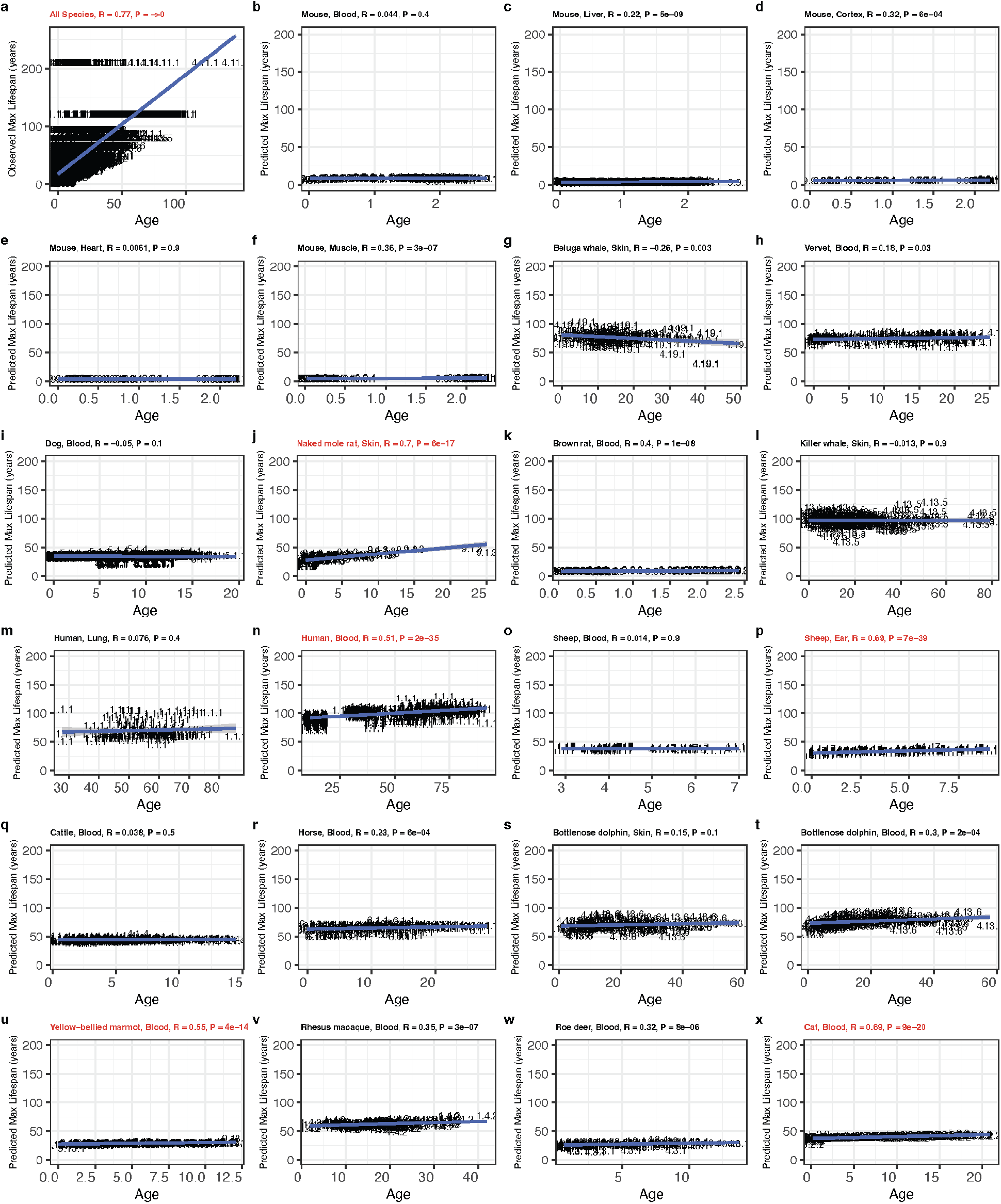
The maximum lifespan predictor is applied to individual samples in comparison to their chronological ages. Mammalian maximum lifespan predictor, based on averaged species methylation, was used to predict individual sample lifespans (in years scale). The predicted values are also stratified by species and tissues. Only species with >100 sample sizes are shown. To demonstrate natural relations between maximum lifespan and chronological age, panel **a** scatter plot shows association between observed maximum lifespan and chronological age of corresponding samples. Each of panels **b–x** shows scatter plots of predicted lifespans converted to original scales vs. chronological age in specific species/tissue combinations. Numbers are the mammalian species number consistent with those in **fig. S1**. Red font is used when the absolute value of the Pearson correlation exceeds 0.5. Numeric values can be found in **table S1**.

**Fig. S3.**
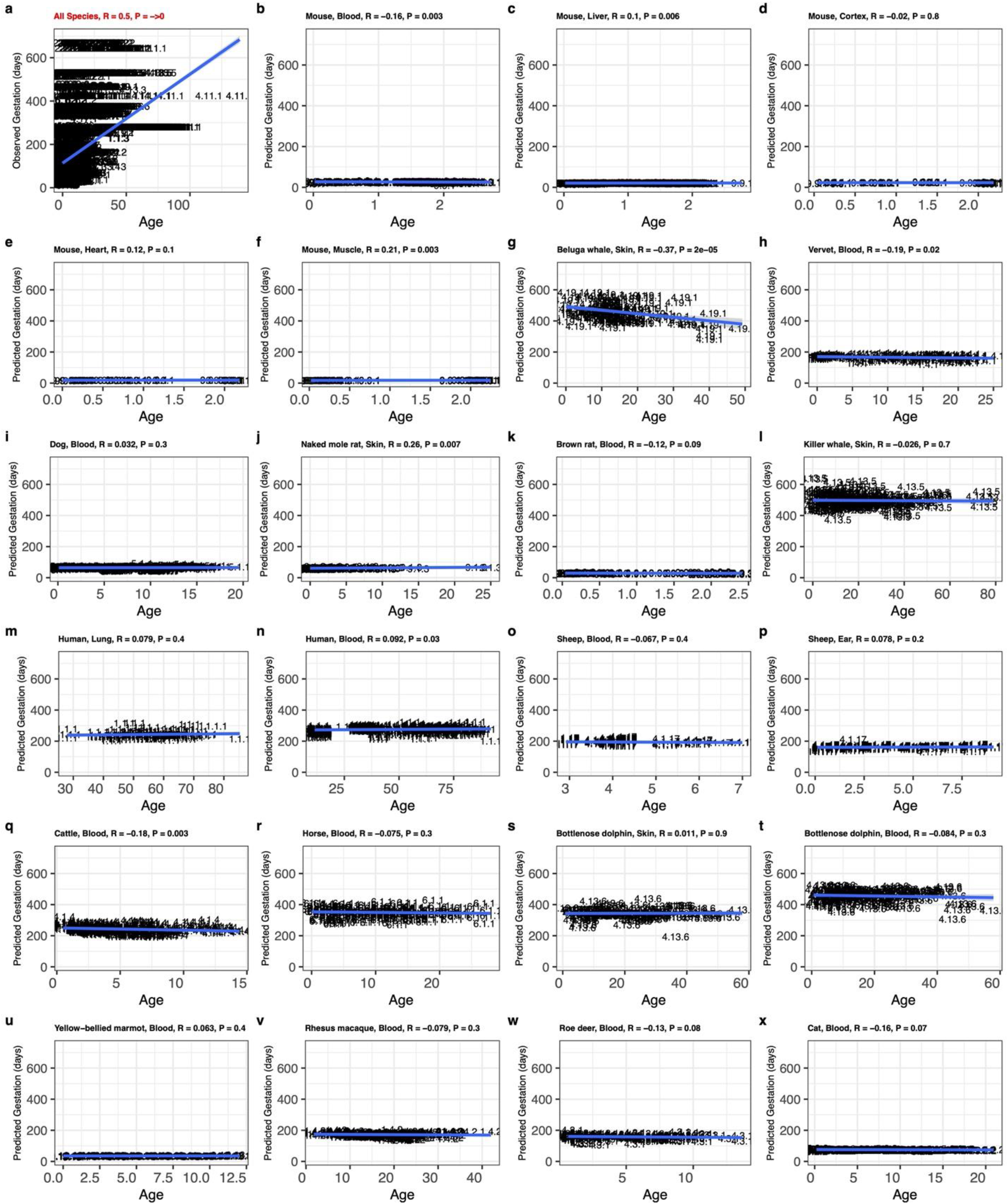
The gestation time predictor is applied to individual samples in comparison to their chronological ages. Gestation time predictor, based on averaged species methylation, was used to predict individual sample gestation time (in log days). The predicted values are also stratified by species and tissues. Only species with >100 sample sizes are shown. To demonstrate natural relations between gestation time (days) and chronological age, panel **a** scatter plot shows association between observed gestation time (days) and chronological age of corresponding samples. Each of panels **b–x** shows scatter plots of predicted gestation time in log-days converted back to days vs. chronological age in specific species. Numbers are the mammalian species number consistent with those in **fig. S1**. Numeric values can be found in **table S1.3**.

**Fig. S4.**
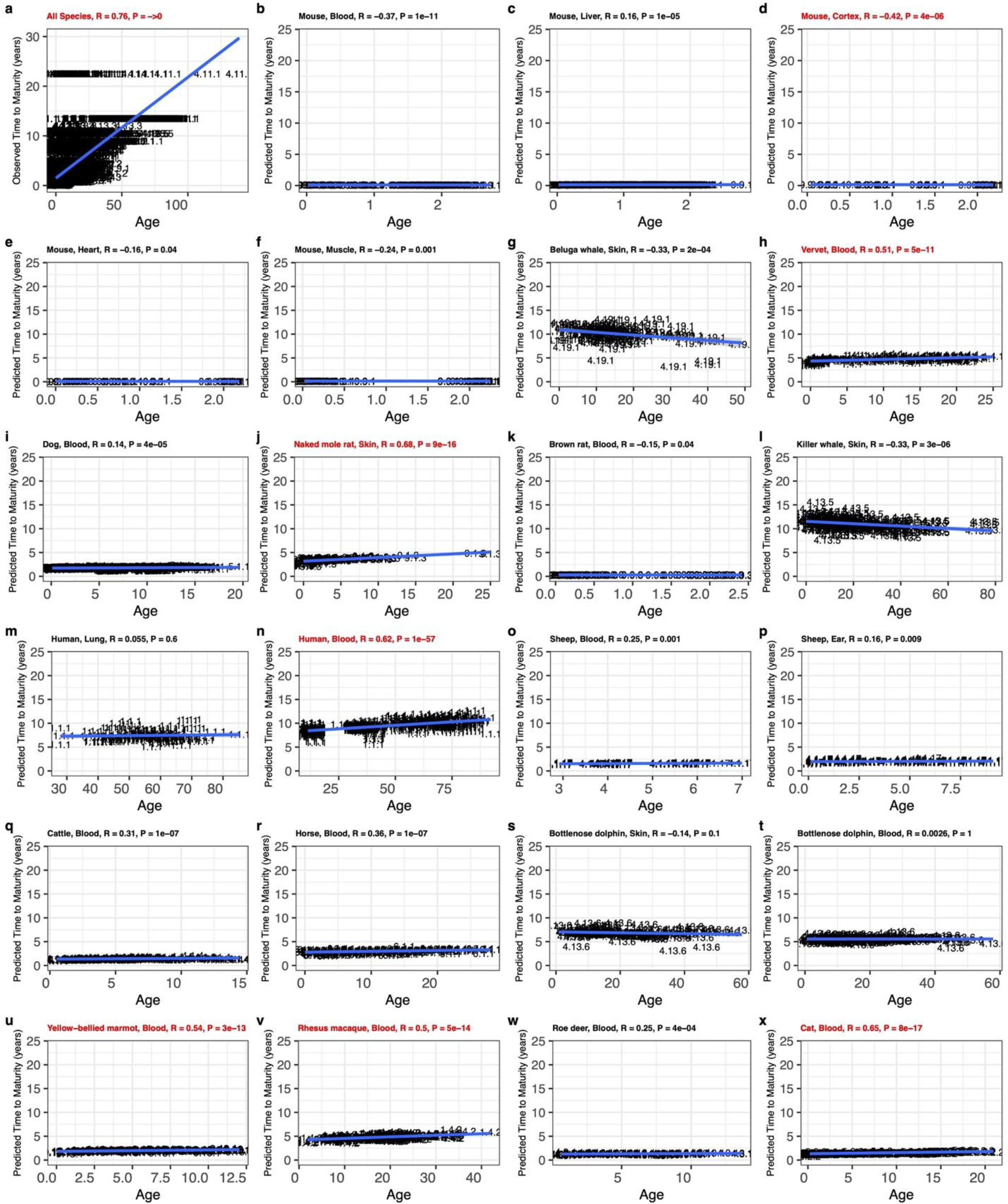
The time to sexual maturity predictor is applied to individual samples in comparison to their chronological ages. Time to sexual maturity predictor, based on averaged species methylation, was used to predict individual sample time to sexual maturity (in log years). The predicted values are also stratified by species and tissues. Only species with >100 sample sizes are shown. To demonstrate natural relations between time to sexual maturity and chronological age, panel **a** scatter plot shows association between time to sexual maturity (years) and chronological age of corresponding samples. **b–x,** scatter plots of predicted age at sexual maturity in log-years converted back to years vs. chronological age in specific species. Numbers are the mammalian species number consistent with those in **fig. S1**. Numeric values can be found in **table S1.3**.

**Fig. S5.**
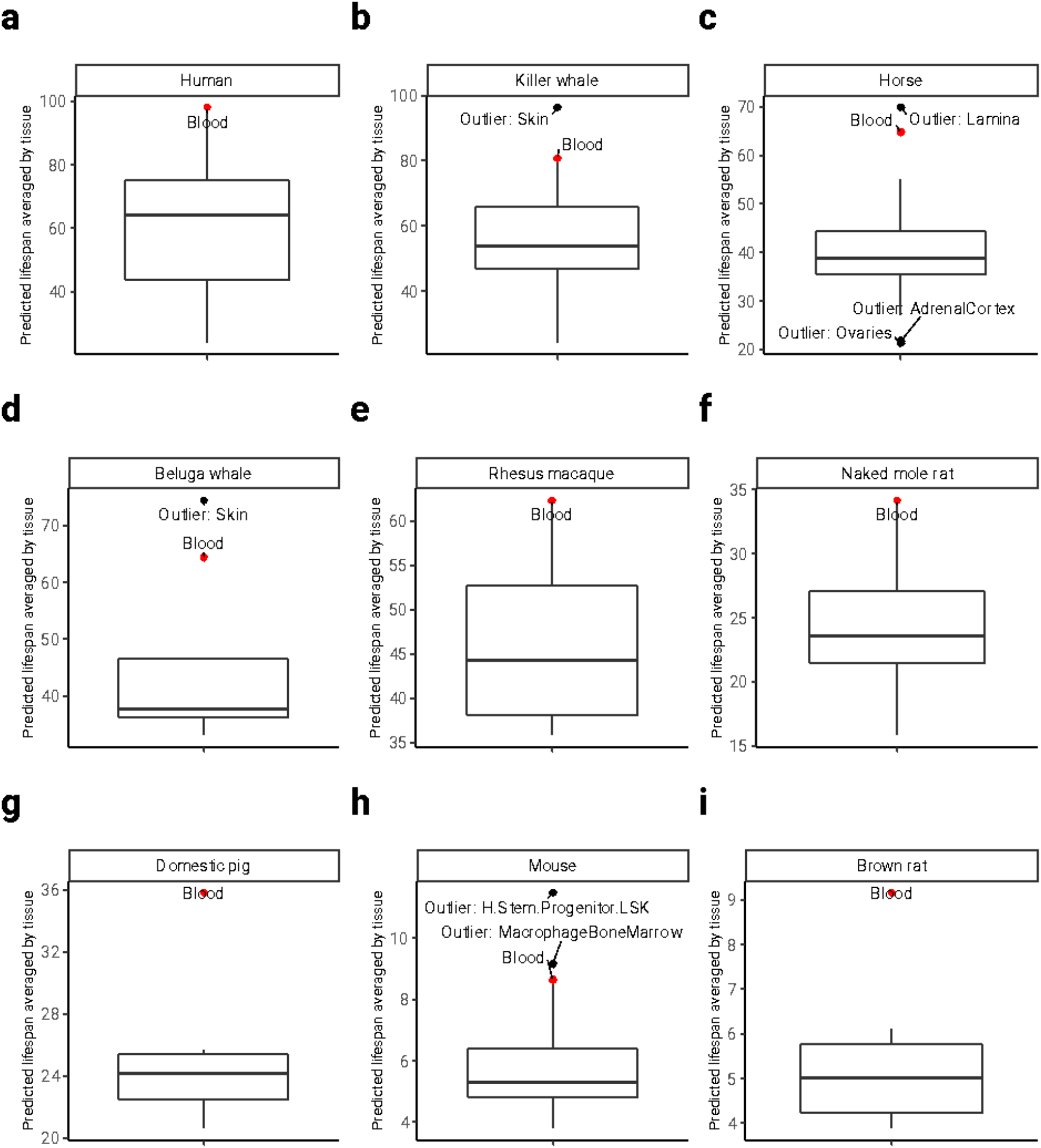
Tissue group differences in predicted mammalian maximum lifespan. Tissue-agnostic predictor of mammalian maximum lifespan, based on averaged species methylation, was used to predict individual maximum lifespan (in log years). The predicted values are aggregated by taking the mean lifespan predictions by tissue groups. Panels **a-i** convert log scale back to original units (lifespan in years); only species with more than 6 different tissue types are shown; mean tissue predicted value outliers are annotated; Tissue type “H.Stem.Progenitor.LSK” stands for “LSK Progenitor Hematopoietic Stem cells.” **c**, Apart from blood, laminae are an outlying tissue in horses. Laminae are interlocking leaf-like tissues that connect the inner surface of the horse’s hoof wall to the bone of the foot. The boxplot, as implemented in the R programming language, provides a visual summary of key statistics from a dataset: The median is represented by the horizontal line inside the box. The interquartile Range (IQR) encompasses the middle 50% of the data. The box’s upper boundary represents the 75th percentile, while the lower boundary represents the 25th percentile. The IQR is the difference between these two values. The whiskers extend to the most extreme data point which is no more than 1.5 times the interquartile range from the box.

**Fig. S6.**
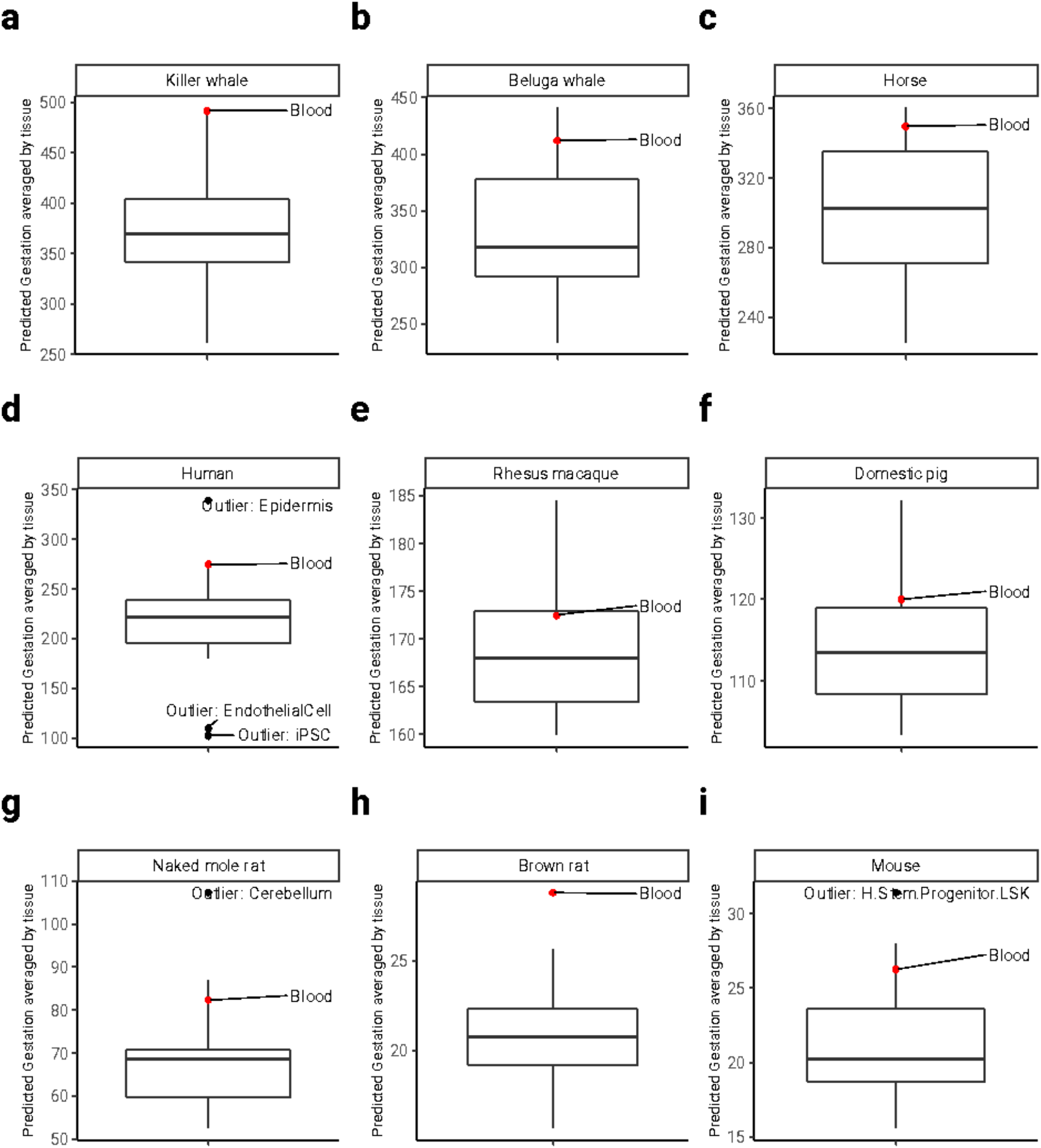
Tissue groups differences in predicted mammalian gestation time. Tissue-agnostic predictor of gestation time, based on averaged species methylation, was used to predict individual sample gestation time (in log days). The predicted values are aggregated by taking the mean gestation time predictions by tissue groups. Panels **a-i** convert log scale back to original units (gestation in days); only species with more than 6 different tissue types are shown; mean tissue predicted value outliers are annotated; Tissue type “H.Stem.Progenitor.LSK” stands for “LSK Progenitor Hematopoietic Stem cells.” The boxplot, as implemented in the R programming language, provides a visual summary of key statistics from a dataset: The median is represented by the horizontal line inside the box. The interquartile Range (IQR) encompasses the middle 50% of the data. The box’s upper boundary represents the 75th percentile, while the lower boundary represents the 25th percentile. The IQR is the difference between these two values. The whiskers extend to the most extreme data point which is no more than 1.5 times the interquartile range from the box.

**Fig. S7.**
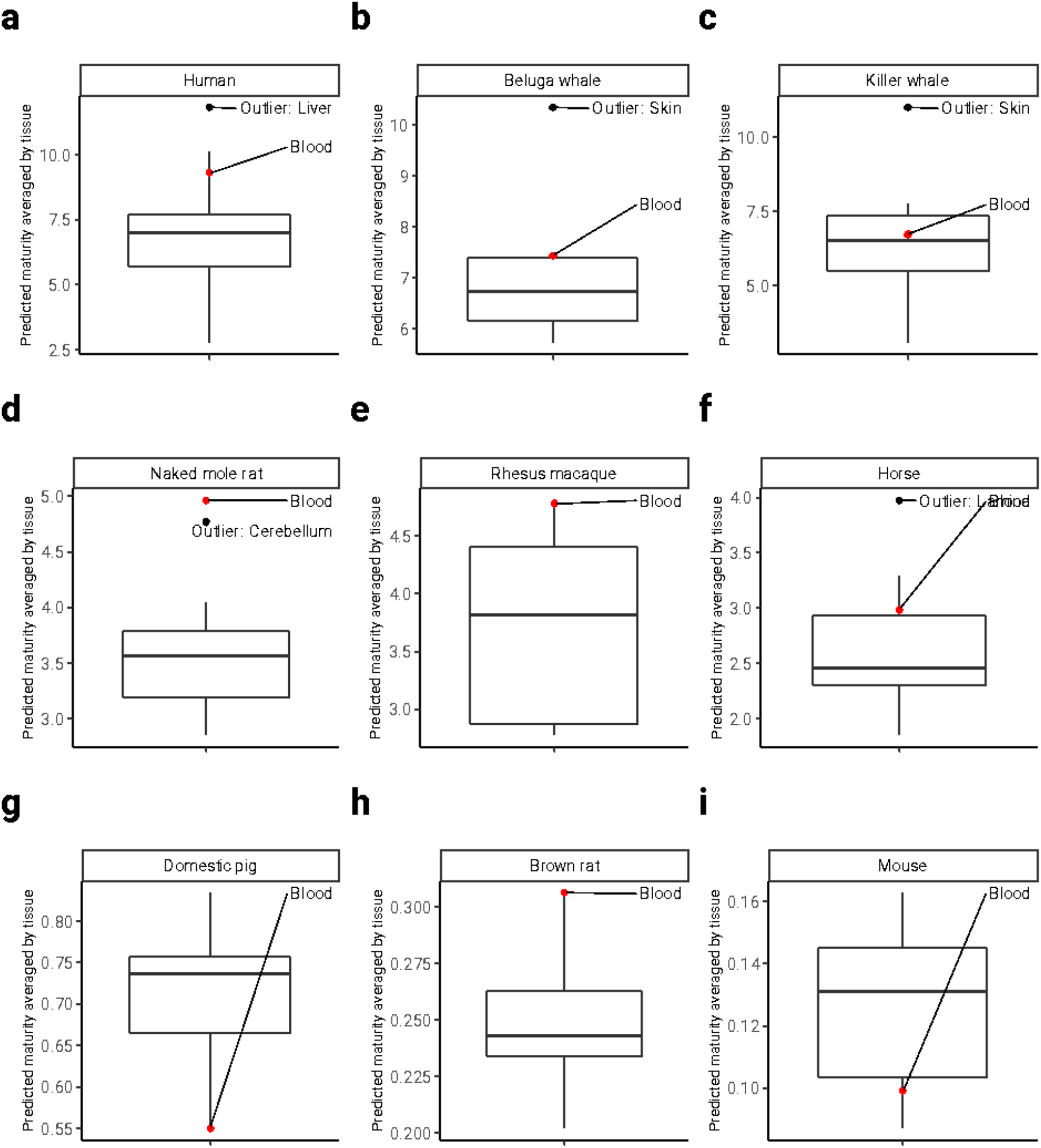
Predicted time to sexual maturity in select species for which multiple tissues were available. Tissue-agnostic predictor of time to sexual maturity. The boxplot shows median predicted values (short horizontal line) across tissue types. Significantly outlying tissues have been highlighted. The boxplot, as implemented in the R programming language, provides a visual summary of key statistics from a dataset: The median is represented by the horizontal line inside the box. The interquartile Range (IQR) encompasses the middle 50% of the data. The box’s upper boundary represents the 75th percentile, while the lower boundary represents the 25th percentile. The IQR is the difference between these two values. The whiskers extend to the most extreme data point which is no more than 1.5 times the interquartile range from the box. Tissue-agnostic predictor of time to sexual maturity predictor, based on averaged species methylation, was used to predict individual sample time to sexual maturity (in log years). The predicted values are aggregated by taking the mean lifespan predictions by tissue groups. Panels **a-i** convert log scale back to original units (age at sexual maturity in years); only species with more than 6 different tissue types are shown; mean tissue predicted value outliers are annotated; Tissue type “H.Stem.Progenitor.LSK” stands for “LSK Progenitor Hematopoietic Stem cells.”

**Fig. S8.**
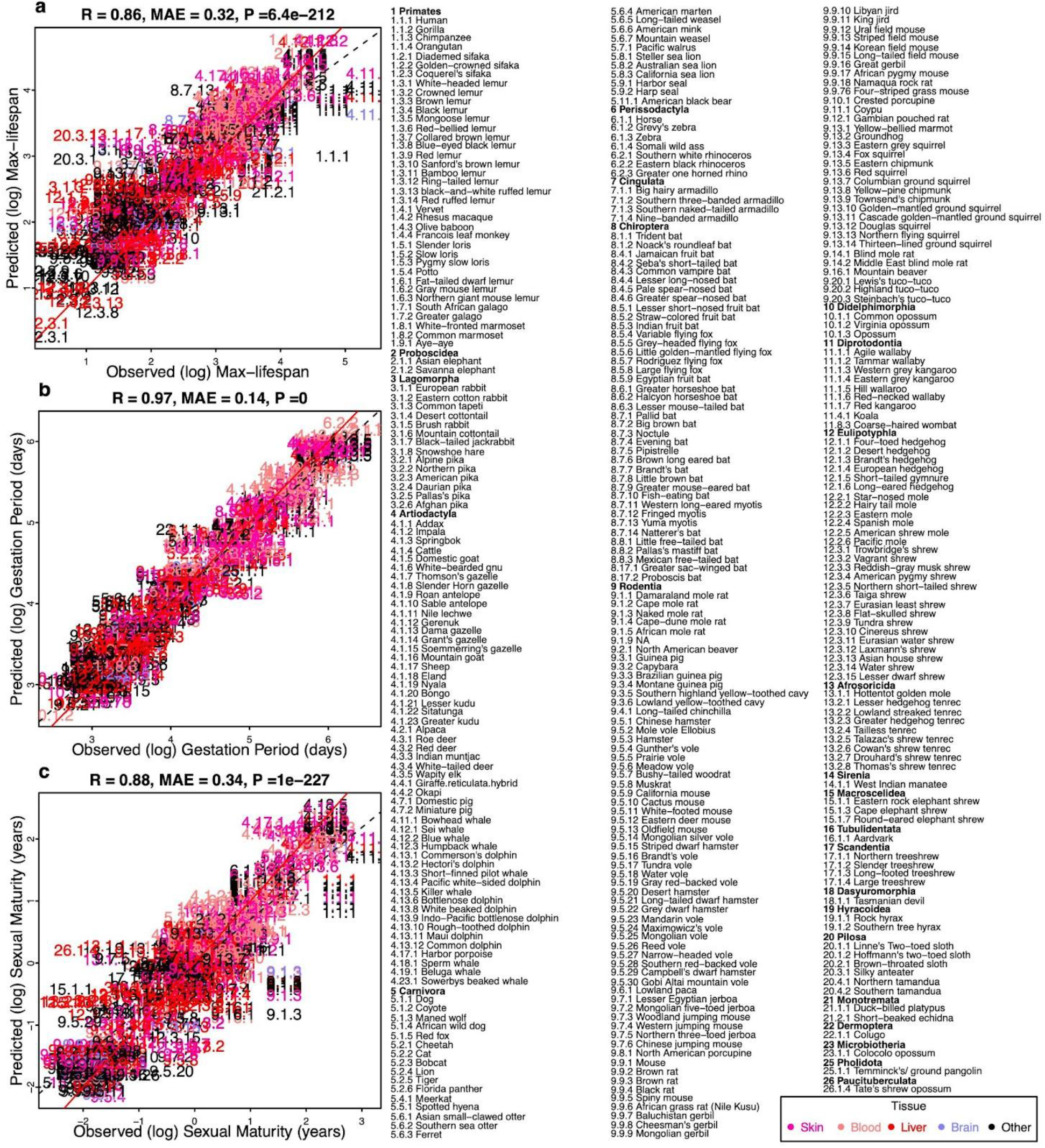
Tissue-aware predictors trained on species-tissue combinations. A penalized joint linear model used to predict species lifespan (Elastic Net). Same framework as that of **Fig. 1**, except that it distinguishes tissue types. CpG probes are averaged by each species-tissue combination. Different tissues within the same species share the same maximum lifespan but retain different methylation levels. Three panels show predictors for **a,** log maximum lifespan (in log years), **b,** log-transformed gestation time (in log days), and **c,** log-transformed age at sexual maturity (in log years). Designated Mammalian numbers in scatter plot panels and the Figure legend are the same as those of main **Fig. 1**. MAE abbreviates median absolute errors from the regression errors; r and p denote Pearson’s correlation and p-values, respectively. Numbers and colors are the mammalian species number and order annotation consistent with those of other Figures. Numeric values can be found in **table S3**. As with the **Fig. 1**, species appear as designated numbers in scatter plot panels; the corresponding common names and taxonomic orders are annotated in Figure legends; the whole number (number before the decimal separator) part of each mammalian number is assigned in accordance with the corresponding taxonomic order. Red solid line represents the perfect prediction line, and the dotted line represents the fitted linear regression line.

**Fig. S9.**
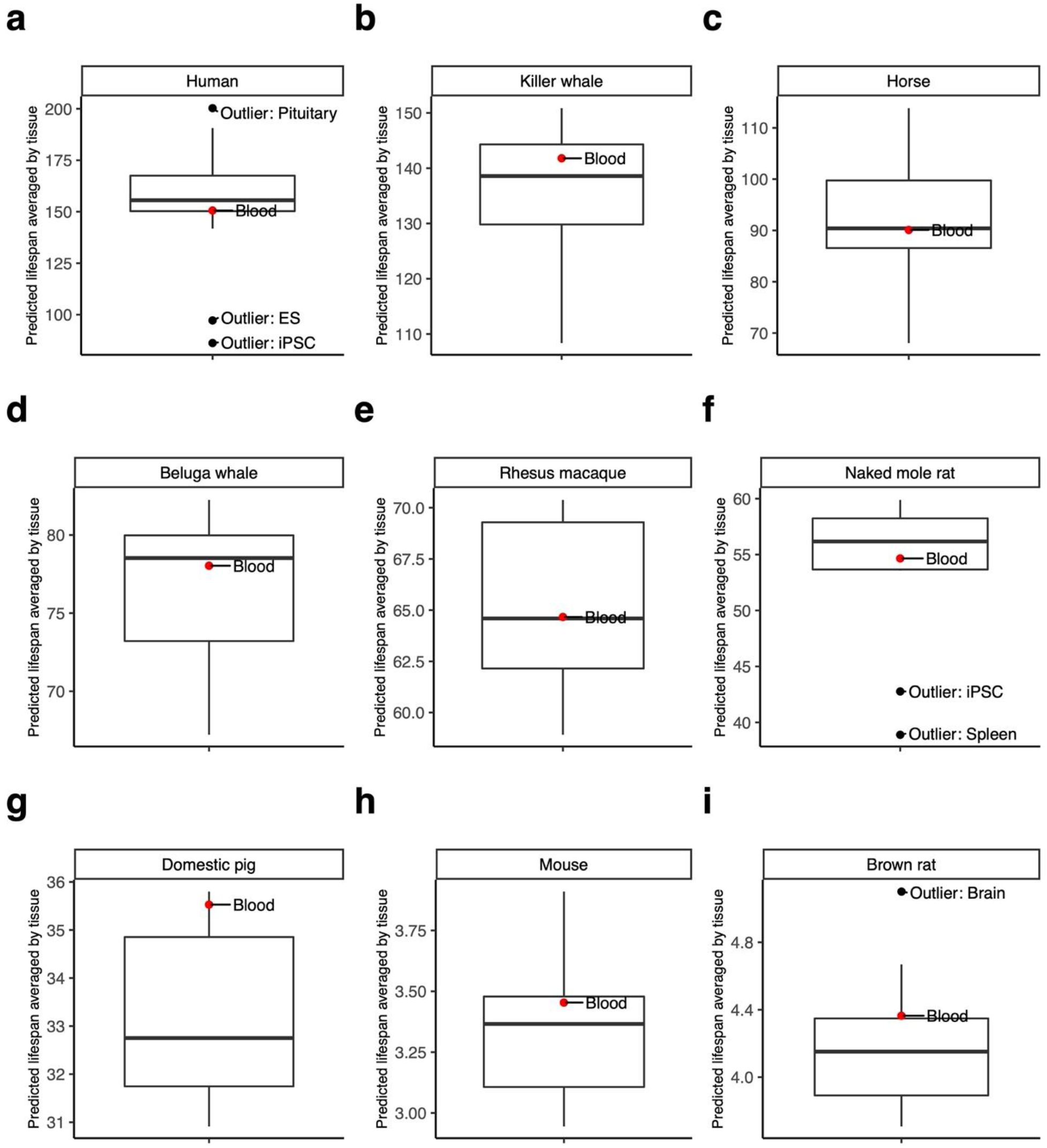
Tissue group differences in predicted mammalian maximum lifespan – Tissue-Aware. Tissue-aware predictor of mammalian lifespan, based on averaged species methylation, was used to predict individual sample lifespan (in log years). The predicted values are aggregated by taking the mean lifespan predictions by tissue groups. Panels **a-i** convert log scale back to original units (lifespan in years); only species with more than 6 different tissue types are shown; mean tissue predicted value outliers are annotated; Tissue type “H.Stem.Progenitor.LSK” stands for “LSK Progenitor Hematopoietic Stem cells.”

**Fig. S10.**
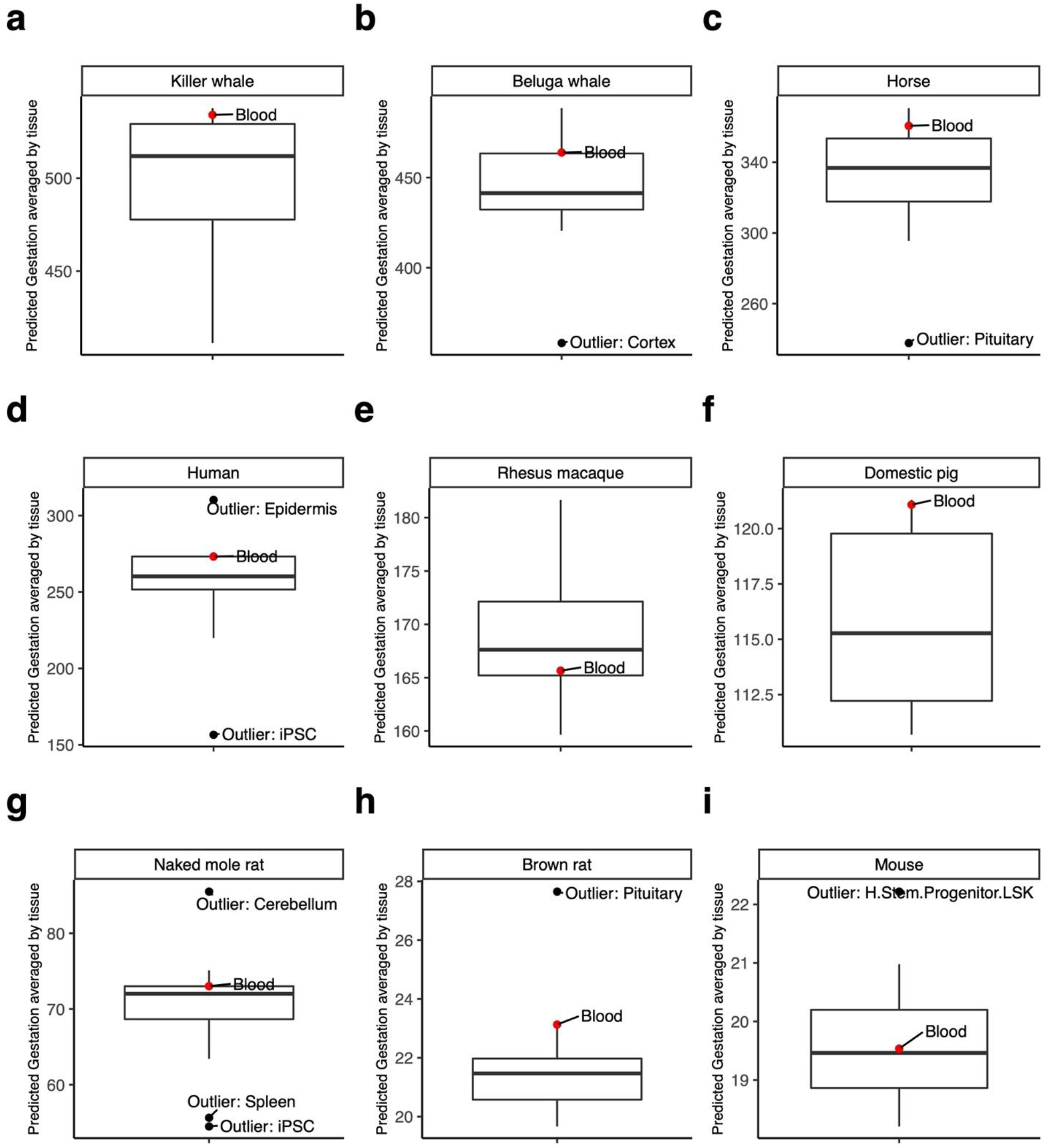
Tissue groups differences in predicted mammalian gestation time – Tissue-aware. Tissue-aware predictor of gestation time, based on averaged species methylation, was used to predict individual sample gestation time (in log days), trained on tissue-aware data. The predicted values are aggregated by taking the mean gestation time predictions by tissue groups. Panels **a-i** convert log scale back to original units (gestation in days); only species with more than 6 different tissue types are shown; mean tissue predicted value outliers are annotated; Tissue type “H.Stem.Progenitor.LSK” stands for “LSK Progenitor Hematopoietic Stem cells.”

**Fig. S11.**
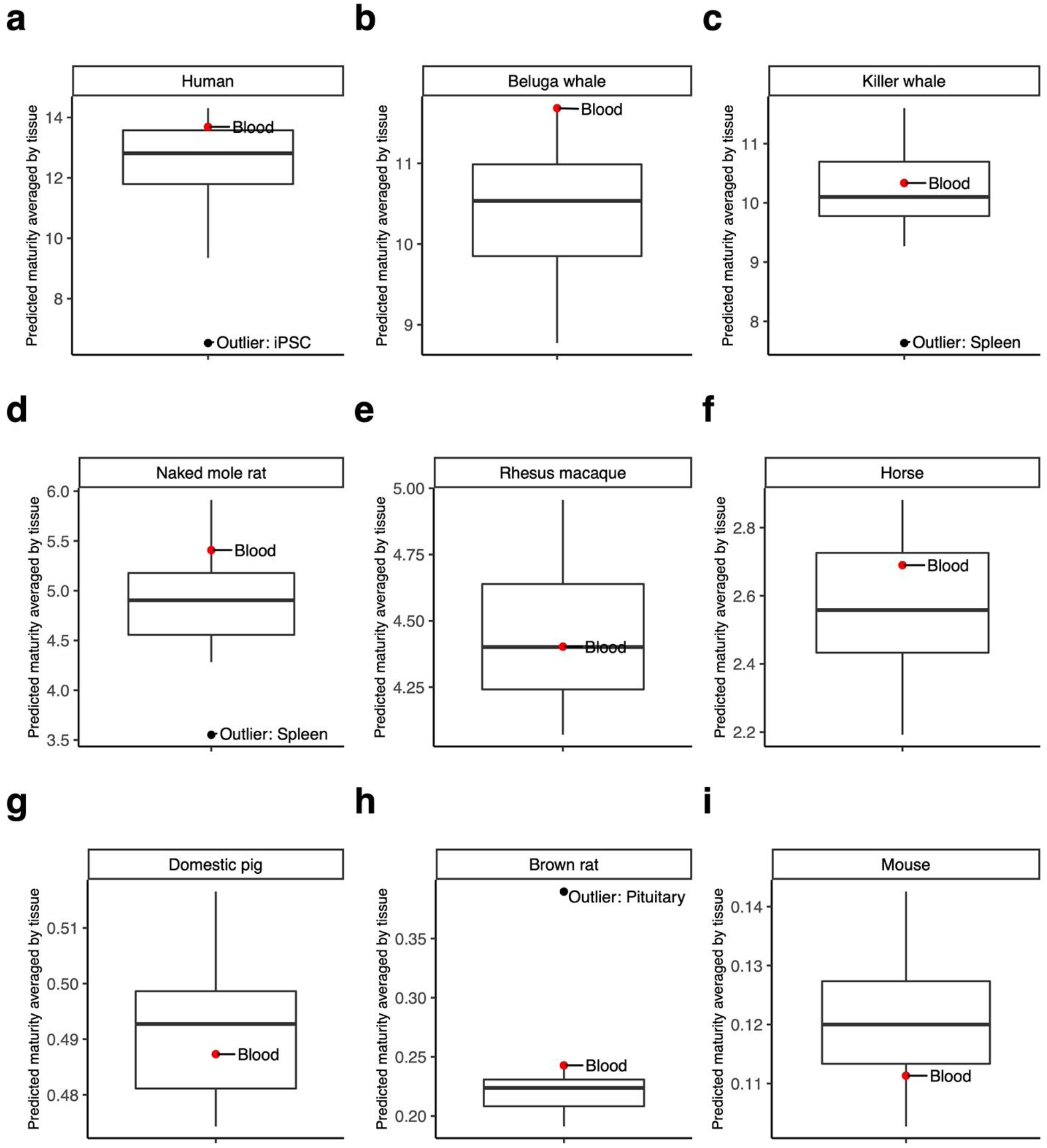
Tissue groups differences in predicted mammalian sexual maturity time – Tissue-aware. Mammalian times to sexual maturity predictor, based on averaged species methylation, was used to predict individual sample time to sexual maturity (in log years), trained on tissue-aware data. The predicted values are aggregated by taking the mean lifespan predictions by tissue groups. Panels **a-i** convert log scale back to original units (age at sexual maturity in years); only species with more than 6 different tissue types are shown; mean tissue predicted value outliers are annotated; Tissue type “H.Stem.Progenitor.LSK” stands for “LSK Progenitor Hematopoietic Stem cells.

**Fig. S12.**
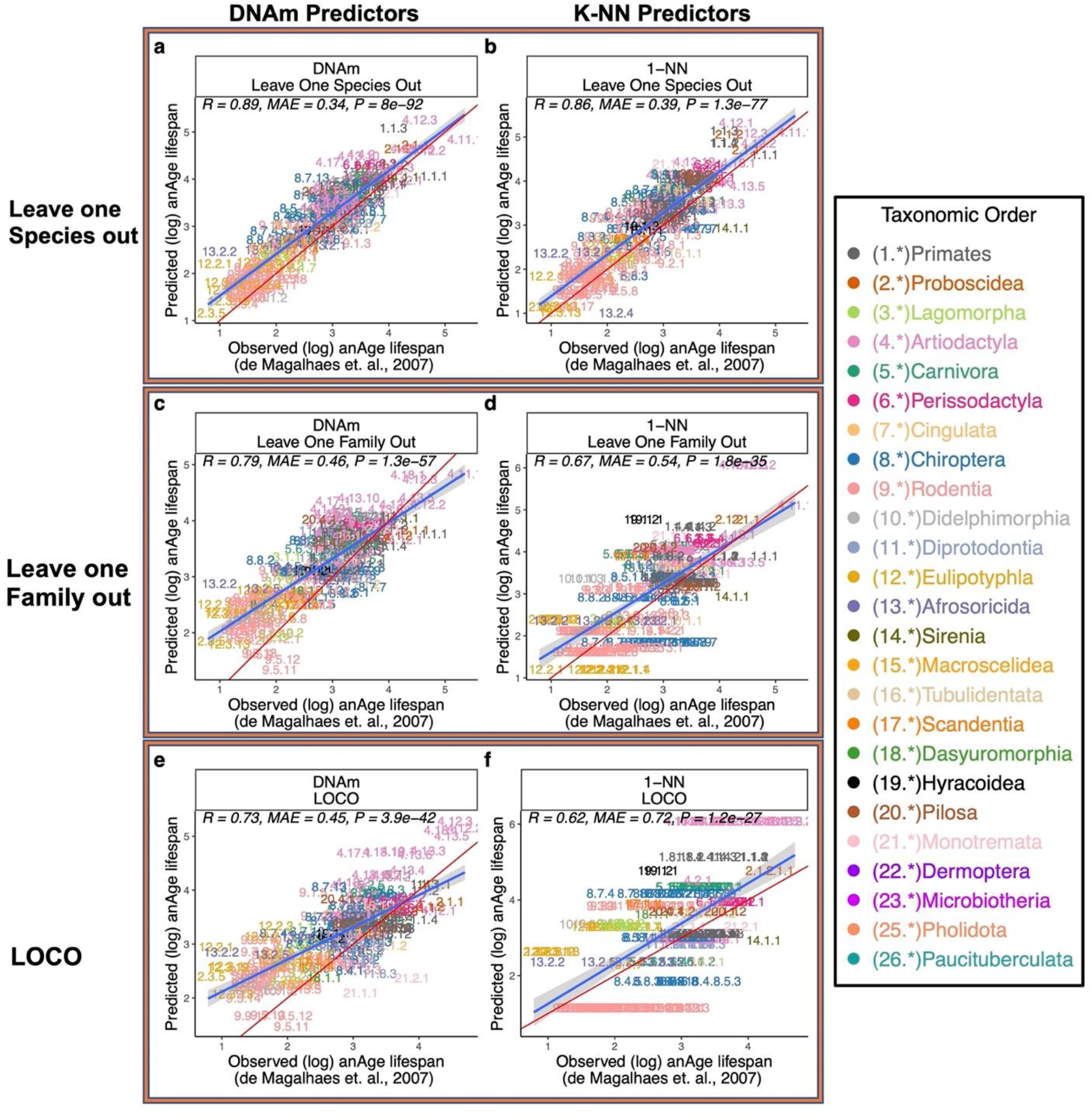
Overall comparisons between DNAm lifespan predictors and phylogeny-based predictors. Various training-test validation analyses of predictors of log (base e) transformed estimates of maximum lifespan. We compared prediction performance between DNAm elastic net predictors and 1-Nearest-Neighbor predictor (k-NN). 1-Nearest-Neighbor predictor utilizes distances from the Mammalian phylogenetic TimeTree (*54*). Results under different training-test separation methods are shown in panels **a**, **b**, DNAm and k-NN predictors test set predictions under leave-one-species-out (LOSO) training-test separation scheme; **c**, **d**, DNAm and k-NN predictors test set predictions under leave-one-family-out training-test separation; **e**, **f**, DNAm and k-NN predictors test set predictions under leave-one-clade-out (LOCO) training-test separation. LOCO (leave-one-clade-out) is defined as, for orders with more than 20 species (Rodentia, Artiodactyla, Chiroptera, Primates, Carnivora, and Eulipotyphla), leaving out all member species except the longest-living and shortest-living species. MAE abbreviates median absolute errors from the regression errors; r and p denote Pearson’s correlation and p-values, respectively. Numbers and colors are the mammalian species number and order annotation consistent with those of other Figures. Numeric values can be found in **table S1**. Shaded areas represent 95% confidence intervals of the simple linear regression line. E).

**Fig. S13.**
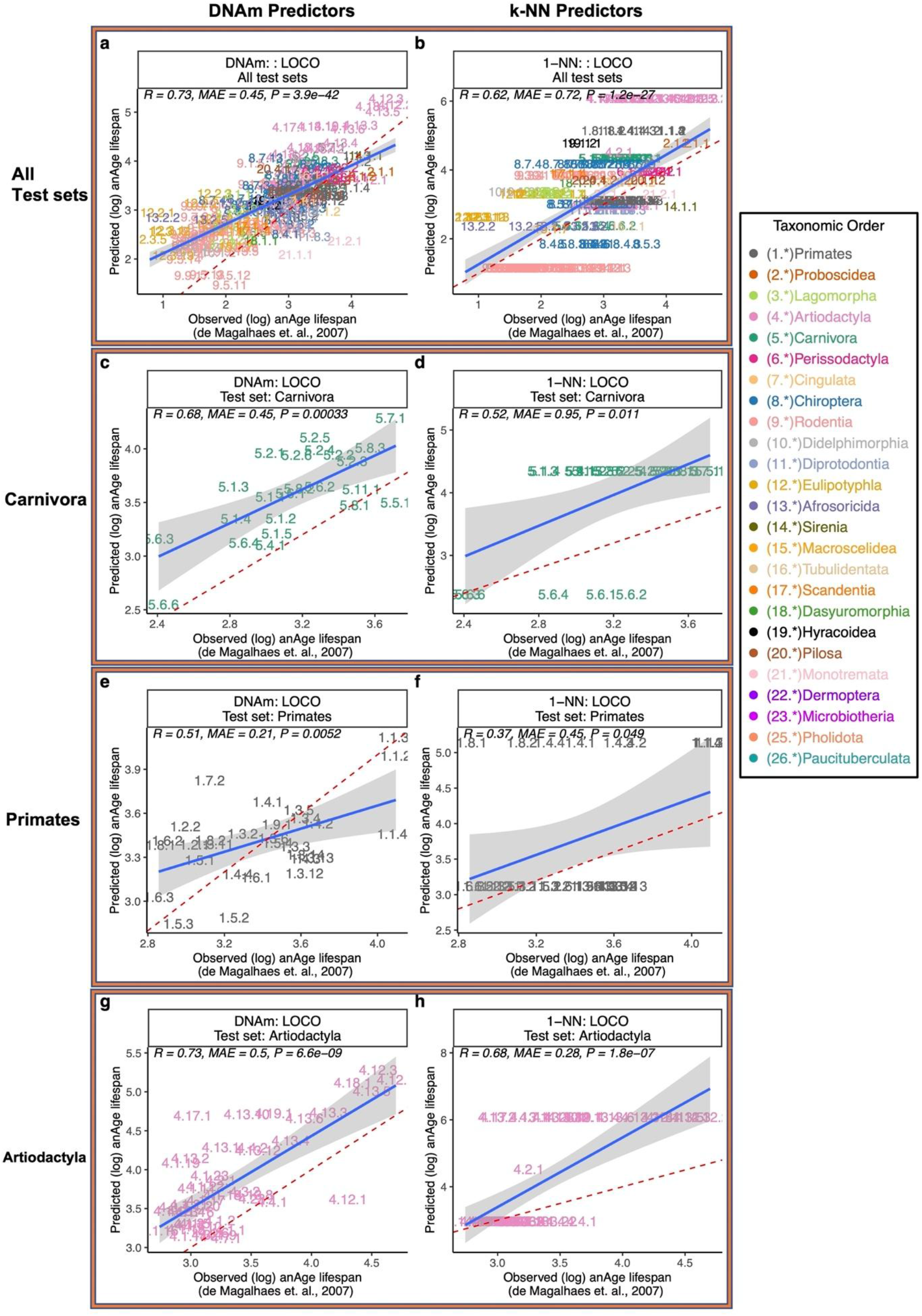
Taxonomic order breakdown of DNAm lifespan predictors and Phylogeny-based Predictors under LOCO. A breakdown of predictor performance in large taxonomic orders under LOCO. We compared prediction performance between DNAm elastic net predictors and 1-Nearest-Neighbor predictor (k-NN). 1-Nearest-Neighbor predictor utilizes distances from the Mammalian phylogenetic TimeTree (*54*). **a**, DNAm predictor’s test set predictions leave-one-clade-out (LOCO) training-test separation scheme; **b**, k-NN predictor’s test set predictions under LOCO; **c**, **d**, DNAm and k-NN predictors, respectively, test set predictions of lifespan for all species belonging to Carnivora under LOCO; **e**, **f**, DNAm and k-NN predictors, respectively, test set predictions of lifespan for all species belonging to Primates under LOCO; **g**, **h** DNAm and k-NN predictors, respectively, test set predictions of lifespan for all species belonging to Artiodactyla under LOCO. MAE abbreviates median absolute errors from the regression errors; r and p denote Pearson’s correlation and p-values, respectively. Numbers and colors are the mammalian species number and order annotation consistent with those of fig. S1. Numeric values can be found in **table S1**. Shaded areas represent 95% confidence intervals of the simple linear regression line. Panels **a** and **b** are analogous to those of **Fig. 2c,d**.

**Fig. S14.**
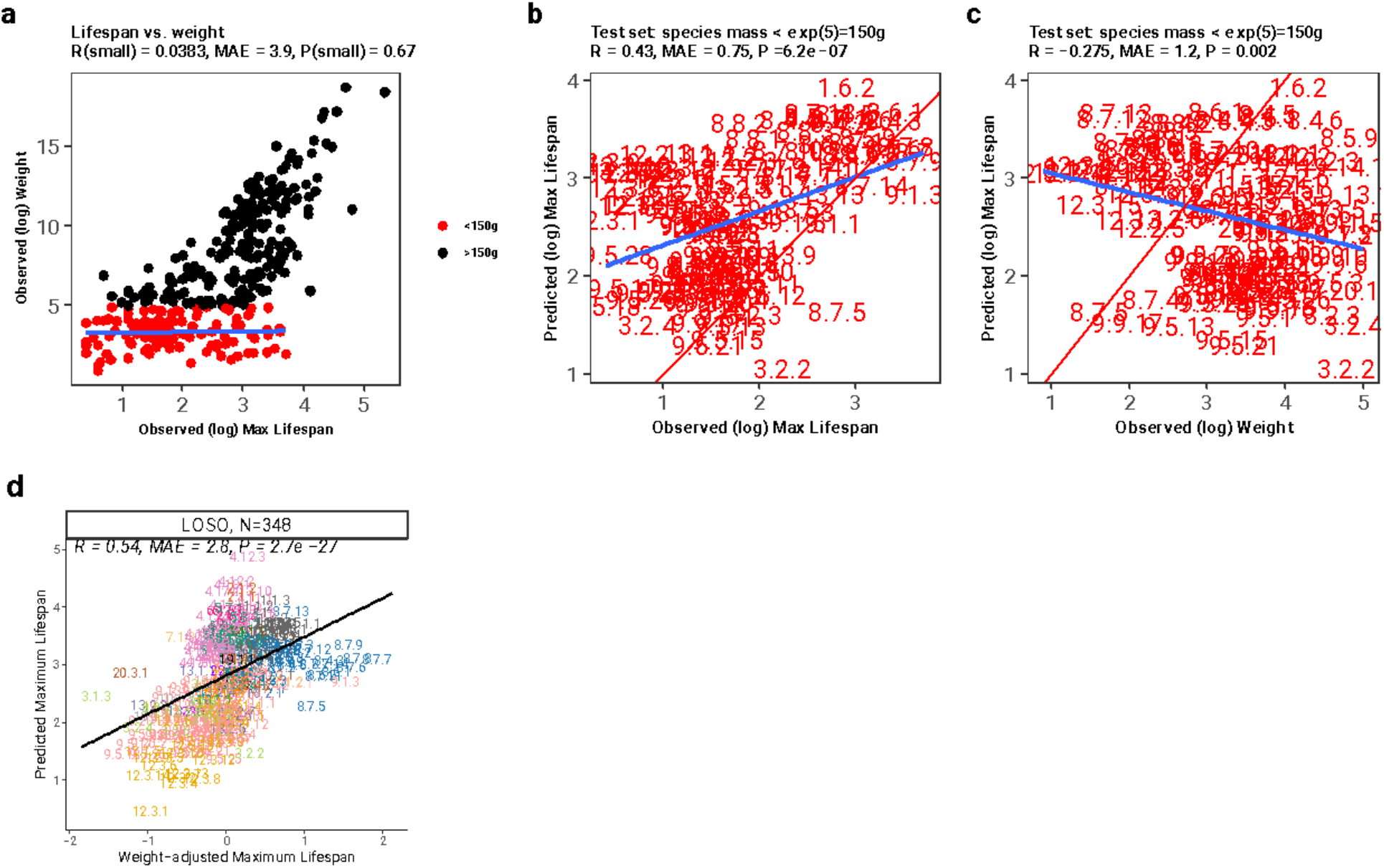
DNAm lifespan predictions do not reflect confounding by adult weight. **a-c**, Report results for a DNAm max lifespan predictor trained on mammal species with an average weight under 150 grams (small mammals). Panels **a**, observed (log) adult body weight vs. observed (log) maximum lifespan in all mammalian species within the data set, color-coded by small-size indicator (more than 150 grams); **b**, test set predictions for the maximum lifespan in small-sized (<150 grams) mammalian species vs. observed (log) maximum lifespan; **c**, test set predictions for the maximum lifespan in small-sized (<150 grams) mammalian species vs. observed (log) adult body weight. MAE abbreviates median absolute errors from the regression errors; r and p denote Pearson’s correlation and p-values, respectively. Numbers are the mammalian species number annotation consistent with those of other Figures. Numeric values can be found in **table S1**. Shaded areas represent 95% confidence intervals of the simple linear regression line. **d.** Results for the final version of the tissue-agnostic DNAm predictor of maximum lifespan. Predicted maximum lifespan (on the log scale, y-axis) versus the corresponding adult weight adjusted version (x-axis). Specifically, the weight adjusted version of log maximum lifespan was defined as raw residual resulting from regressing log transformed predicted maximum lifespan on the log transformed average adult weight of the species.

**Fig. S15.**
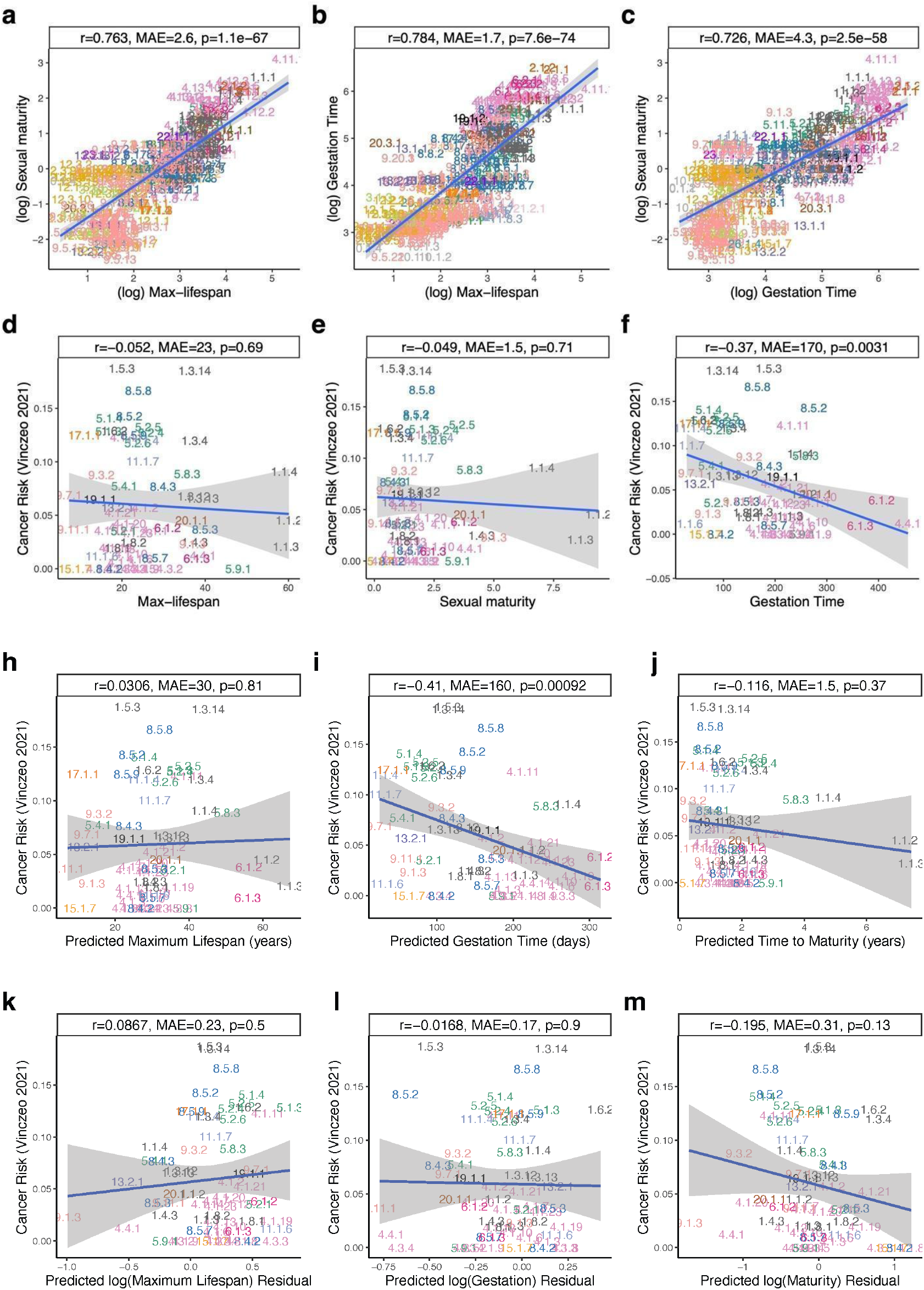
Relationships between observed and epigenetic estimates of mammalian life history traits, including mammalian cancer risk. **a-f**, Panels depict log-transformed relationships between observed variables: **a**. Age at sexual maturity and maximum lifespan **b**. Gestation time and maximum lifespan **c**. Sexual maturity time and gestation time **d**. Cancer risk and maximum lifespan **e**. Cancer risk and sexual maturity **f.** cancer risk and gestation time. **h-j**, estimates of mammalian cancer risk (Vinczeo 2021, y-axis) are plotted against their corresponding epigenetic estimates: **h.** Maximum lifespan **i**. Gestation time **j**. age at sexual maturity. **k-m**, this set is analogous to **h-j**, but the x-axis reports residuals derived from regressing the epigenetic estimate of the life history trait on its observed value (on the log scale): **k**. Log maximum lifespan **l**. Log gestation time **m**. Log-transformed age at sexual maturity. “MAE” represents median absolute errors from the regression errors, while “r” and “p” signify Pearson’s correlation and p-values, respectively. Numbering and colors correspond to the mammalian species number and order, consistent with those in **Fig. 1**. Shaded areas illustrate the 95% confidence intervals of the simple linear regression line. log denotes the natural logarithm, i.e., base e.

**Fig. S16.**
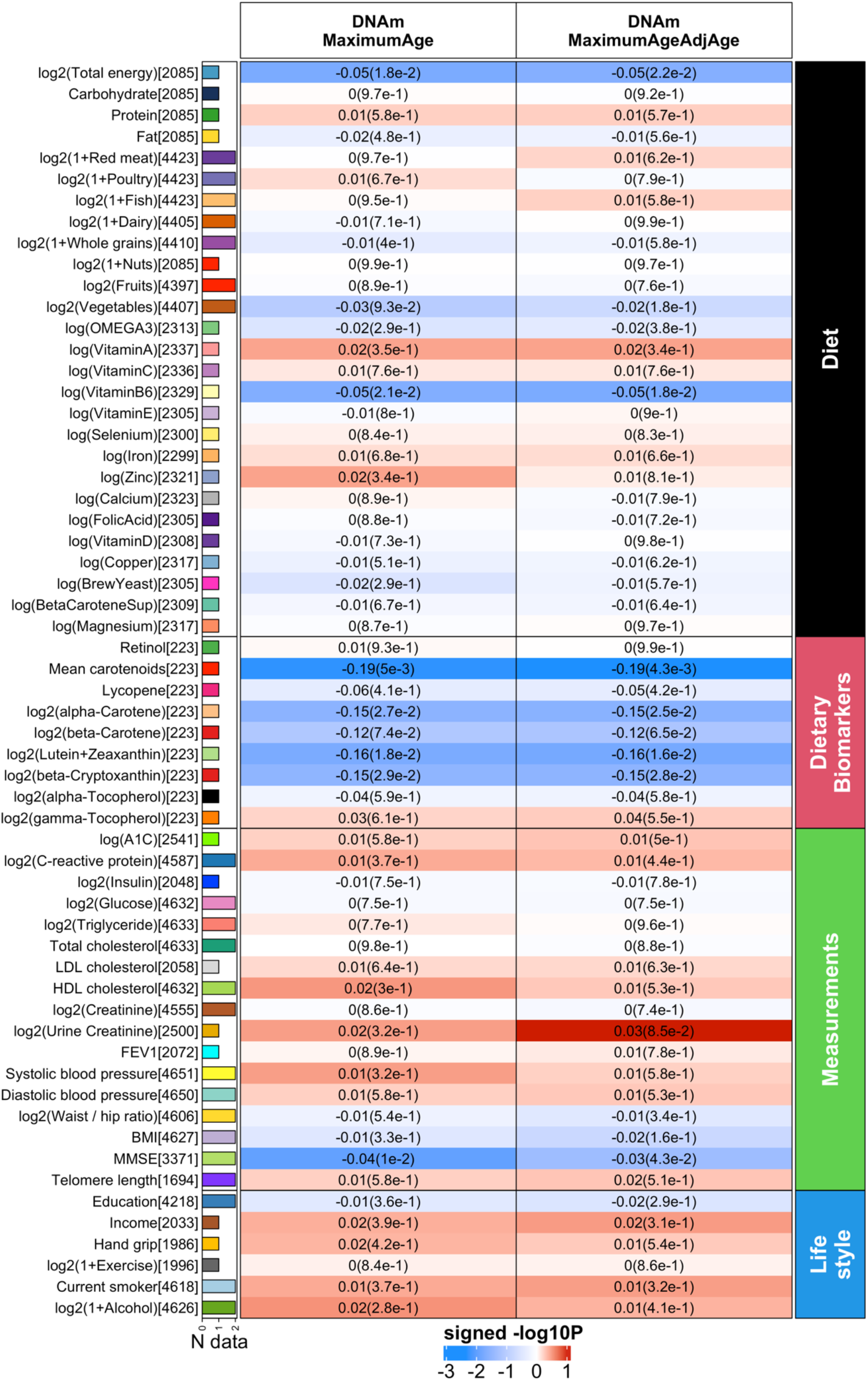
Human epidemiological cohort studies of diet and clinical biomarkers. We performed a correlation analysis between (1) our methylation-based estimator of maximum lifespan (first column) and its age adjusted version (second column) and (2) 59 variables spanning diet, clinically relevant measurements, and lifestyle factors. Comprehensive details of these variables can be found in (*18*). We conducted a robust correlation analysis (biweight midcorrelation, bicor) between (1) our methylation based measures (columns), and (2) 59 variables encompassing 27 self-reported dietary factors, 9 dietary biomarkers, 17 clinical measurements related to vital signs, metabolic traits, inflammatory markers, cognitive and lung function, central adiposity, leukocyte telomere length, and 6 lifestyle factors. This bicor analysis was applied to individuals from both the Framingham Heart Study (up to n=2544) and Women’s Health Initiative (up to n=2107), stratified by gender and ethnic category within each respective cohort. The results were consolidated using fixed-effects meta-analysis models, weighted by inverse variance, generating a meta-estimate of bicor and meta P-value. The clinical biomarkers in FHS offspring cohort were measured during the 8th examination aligned with the measures of DNA methylation profiles. The 9 dietary biomarkers, however, were only available in the WHI cohort, with measurements taken from fasting plasma collected at baseline. Food groups and nutrients considered were comprehensive, encompassing all types and preparation methods; for instance, folic acid included both synthetic and natural forms, and dairy encompassed cheese and all varieties of milk. Further details on the individual diet variables of the WHI cohort can be found in our previous study (*57*).

