## Supplemental Figures for "Epigenetic predictors of species maximum lifespan and other life history traits in mammals"

### Supplementary Figures

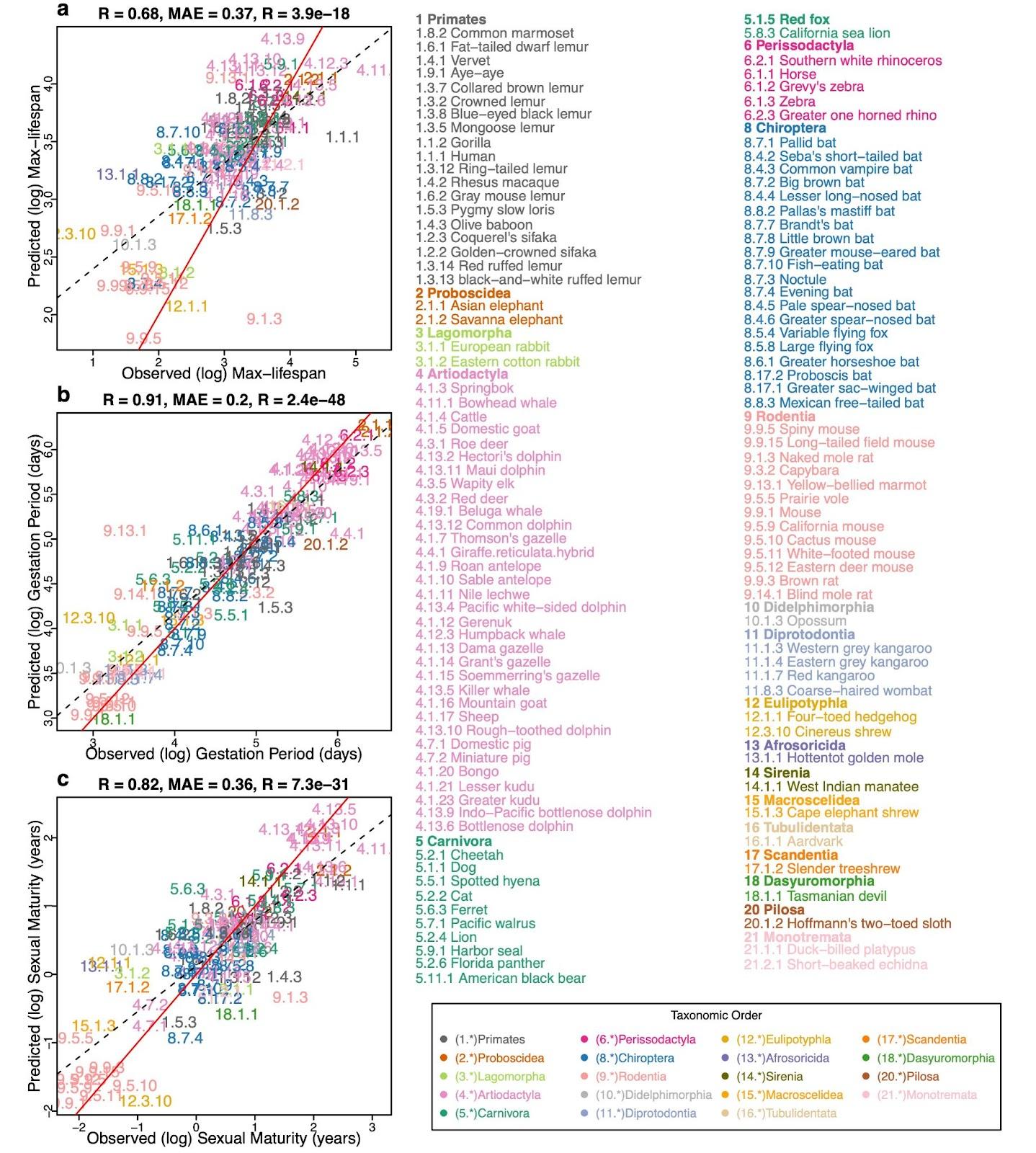

#### Fig. S1 | ElasticNet predictor based on young samples.

Elastic Net Predictor, Leave-one-species-out analysis, fitted on a subset of all young samples (species n = 122). Young samples are defined as samples whose age is both younger than five years and less than the species’ average age at sexual maturation. Feature filtering and Elastic Net tuning parameter set-up is the same as those for **Fig. 1**. Three panels show predictors for **a,** log maximum lifespan (in log years), **b,** log-transformed gestation time (in log days), and **c,** log-transformed age at sexual maturity (in log years). As with the **Fig. 1**, species appear as designated numbers in scatter plot panels; the corresponding common names and phylogenetic orders are annotated in Figure legends; as indicated by the taxonomic order legend, the whole number (number before the decimal separator) part of each mammalian number is assigned in accordance with the corresponding taxonomic order. MAE abbreviates median absolute errors from the regression errors; r and p denote Pearson’s correlation and p-values, respectively. Numbers and colors are the mammalian species number and order annotation consistent with those of other Figures. Numeric values can be found in **table S1.3**. Red solid line represents the perfect prediction line, and the dotted line represents the fitted linear regression line.

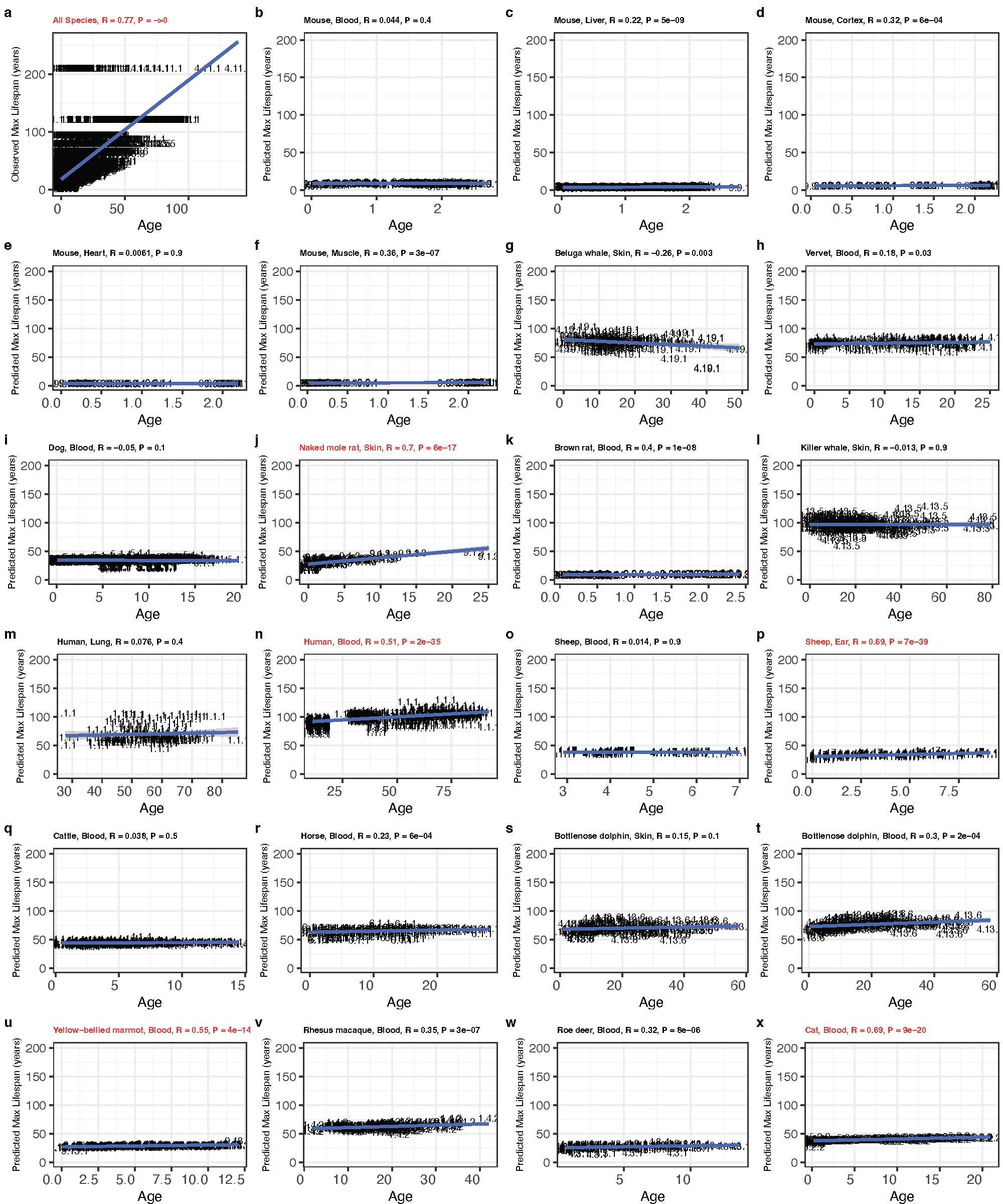

#### Fig. S2 | The maximum lifespan predictor is applied to individual samples in comparison to their chronological ages.

Mammalian maximum lifespan predictor, based on averaged species methylation, was used to predict individual sample lifespans (in years scale). The predicted values are also stratified by species and tissues. Only species with >100 sample sizes are shown. To demonstrate natural relations between maximum lifespan and chronological age, panel **a** scatter plot shows association between observed maximum lifespan and chronological age of corresponding samples. Each of panels **b–x** shows scatter plots of predicted lifespans converted to original scales vs. chronological age in specific species/tissue combinations. Numbers are the mammalian species number consistent with those in **fig. S1**. Red font is used when the absolute value of the Pearson correlation exceeds 0.5. Numeric values can be found in **table S1**.

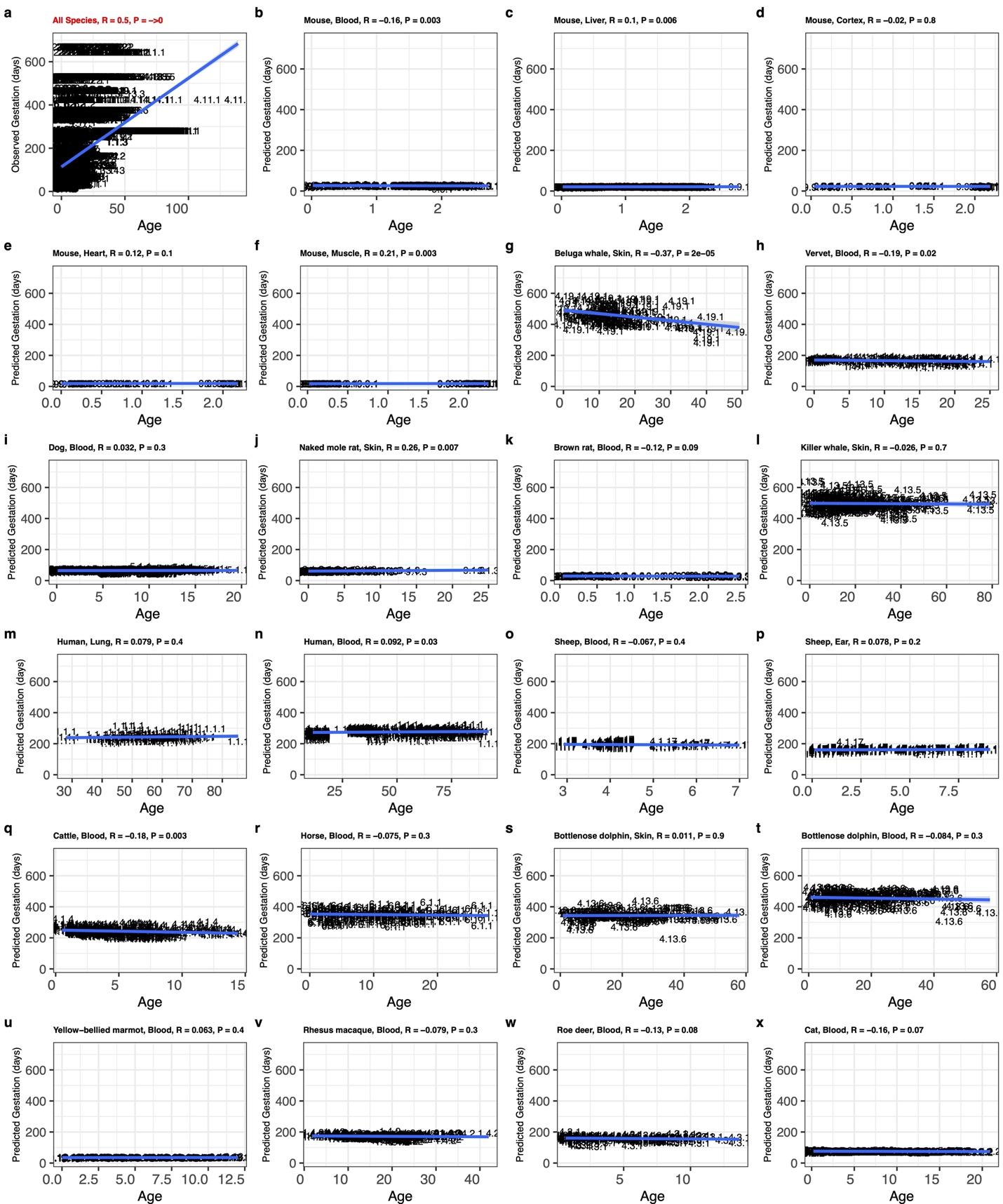

#### Fig. S3 | The gestation time predictor is applied to individual samples in comparison to their chronological ages.

Gestation time predictor, based on averaged species methylation, was used to predict individual sample gestation time (in log days). The predicted values are also stratified by species and tissues. Only species with >100 sample sizes are shown. To demonstrate natural relations between gestation time (days) and chronological age, panel **a** scatter plot shows association between observed gestation time (days) and chronological age of corresponding samples. Each of panels **b–x** shows scatter plots of predicted gestation time in log-days converted back to days vs. chronological age in specific species. Numbers are the mammalian species number consistent with those in **fig. S1**. Numeric values can be found in **table S1.3**.
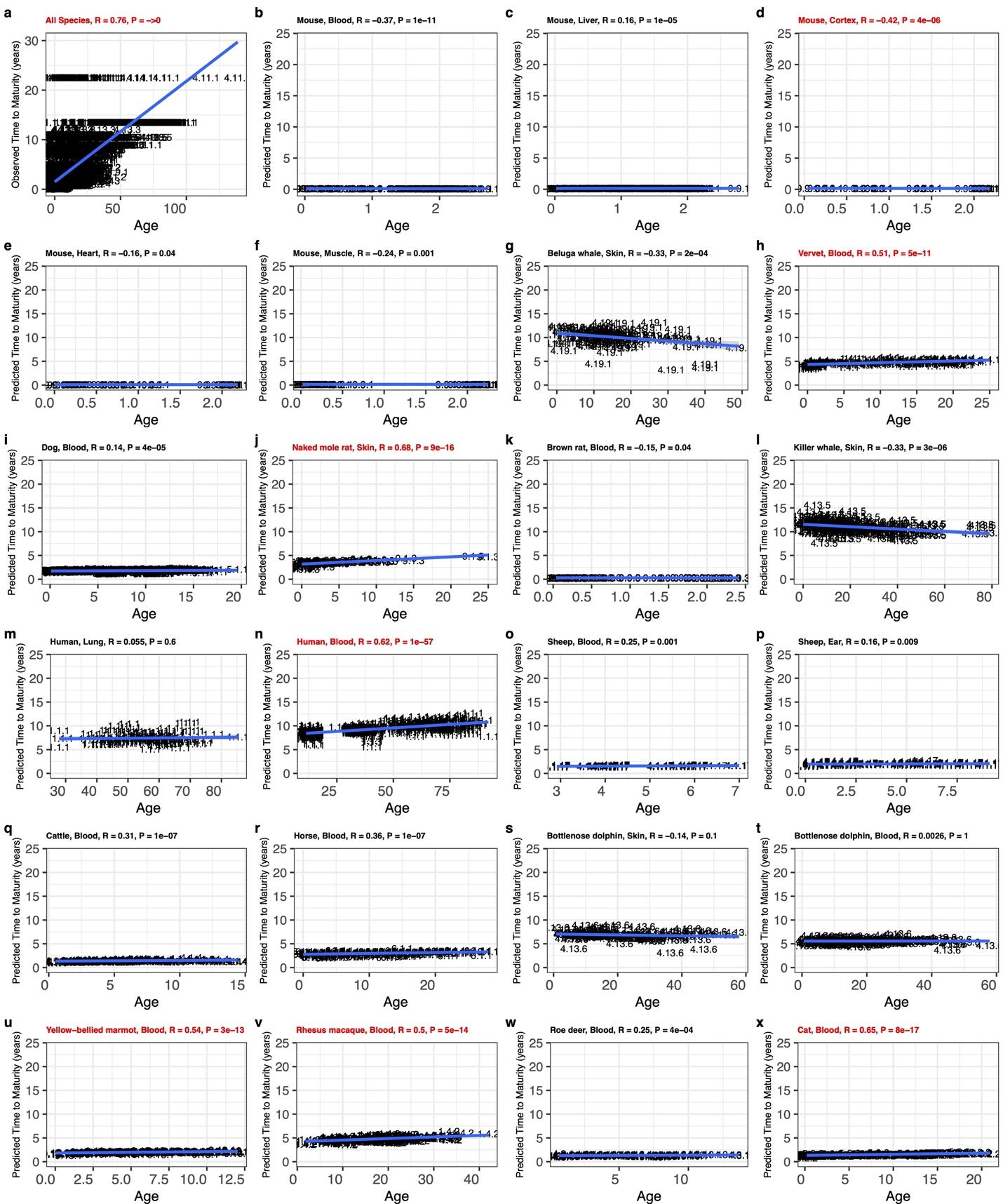

#### Fig. S4 | The time to sexual maturity predictor is applied to individual samples in comparison to their chronological ages.

Time to sexual maturity predictor, based on averaged species methylation, was used to predict individual sample time to sexual maturity (in log years). The predicted values are also stratified by species and tissues. Only species with >100 sample sizes are shown. To demonstrate natural relations between time to sexual maturity and chronological age, panel **a** scatter plot shows association between time to sexual maturity (years) and chronological age of corresponding samples. **b–x,** scatter plots of predicted age at sexual maturity in log-years converted back to years vs. chronological age in specific species. Numbers are the mammalian species number consistent with those in **fig. S1**. Numeric values can be found in **table S1.3**.

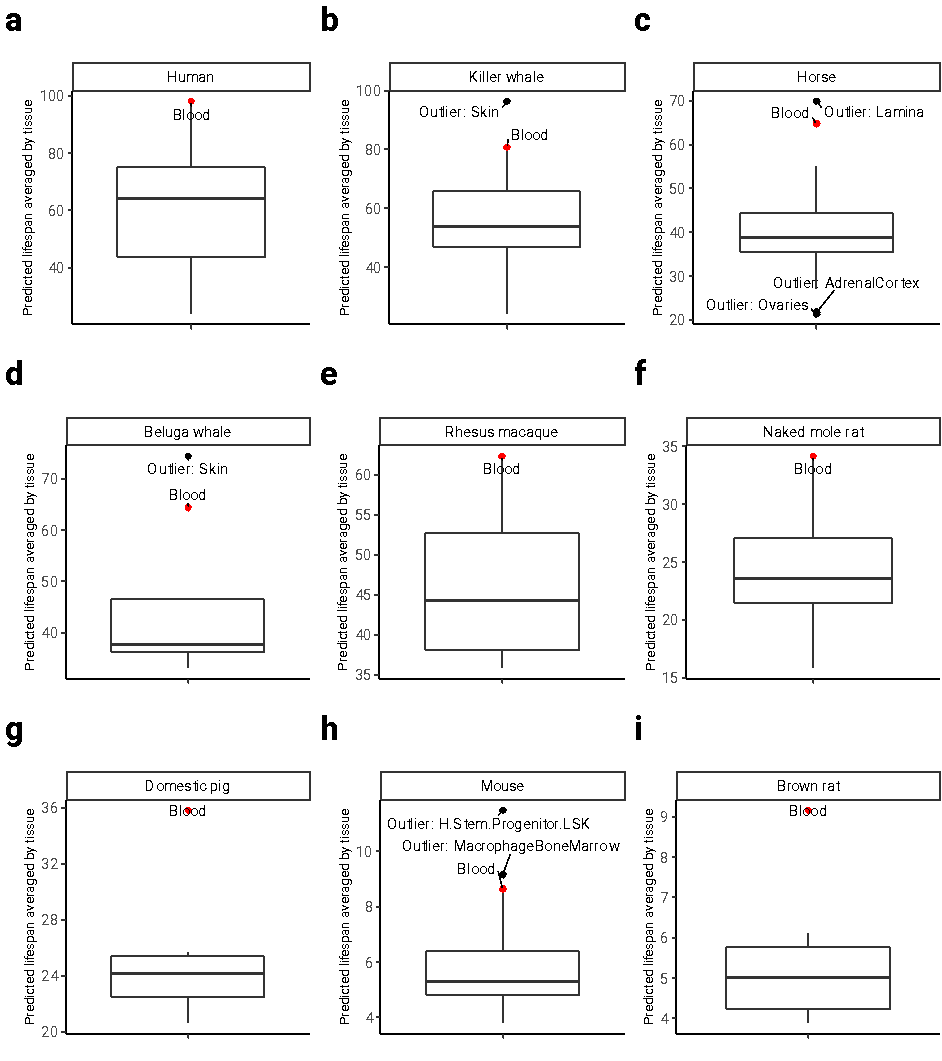

#### Fig. S5 | Tissue group differences in predicted mammalian maximum lifespan.

Tissue-agnostic predictor of mammalian maximum lifespan, based on averaged species methylation, was used to predict individual maximum lifespan (in log years). The predicted values are aggregated by taking the mean lifespan predictions by tissue groups. Panels **a-i** convert log scale back to original units (lifespan in years); only species with more than 6 different tissue types are shown; mean tissue predicted value outliers are annotated; Tissue type “H.Stem.Progenitor.LSK” stands for “LSK Progenitor Hematopoietic Stem cells.” **c**, Apart from blood, laminae are an outlying tissue in horses. Laminae are interlocking leaf-like tissues that connect the inner surface of the horse's hoof wall to the bone of the foot. The boxplot, as implemented in the R programming language, provides a visual summary of key statistics from a dataset: The median is represented by the horizontal line inside the box. The interquartile Range (IQR) encompasses the middle 50% of the data. The box's upper boundary represents the 75th percentile, while the lower boundary represents the 25th percentile. The IQR is the difference between these two values. The whiskers extend to the most extreme data point which is no more than 1.5 times the interquartile range from the box.

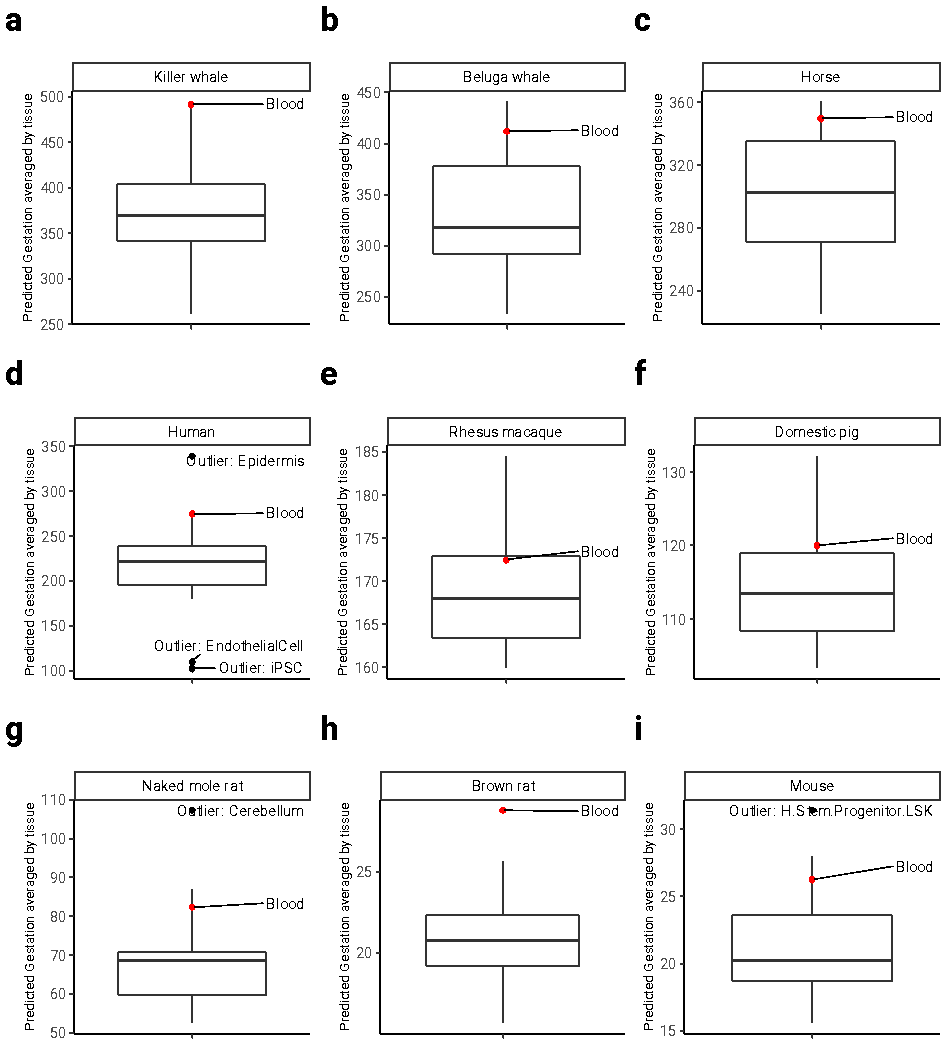

#### Fig. S6 | Tissue groups differences in predicted mammalian gestation time.

Tissue-agnostic predictor of gestation time, based on averaged species methylation, was used to predict individual sample gestation time (in log days). The predicted values are aggregated by taking the mean gestation time predictions by tissue groups. Panels **a-i** convert log scale back to original units (gestation in days); only species with more than 6 different tissue types are shown; mean tissue predicted value outliers are annotated; Tissue type “H.Stem.Progenitor.LSK” stands for “LSK Progenitor Hematopoietic Stem cells.” The boxplot, as implemented in the R programming language, provides a visual summary of key statistics from a dataset: The median is represented by the horizontal line inside the box. The interquartile Range (IQR) encompasses the middle 50% of the data. The box's upper boundary represents the 75th percentile, while the lower boundary represents the 25th percentile. The IQR is the difference between these two values. The whiskers extend to the most extreme data point which is no more than 1.5 times the interquartile range from the box.

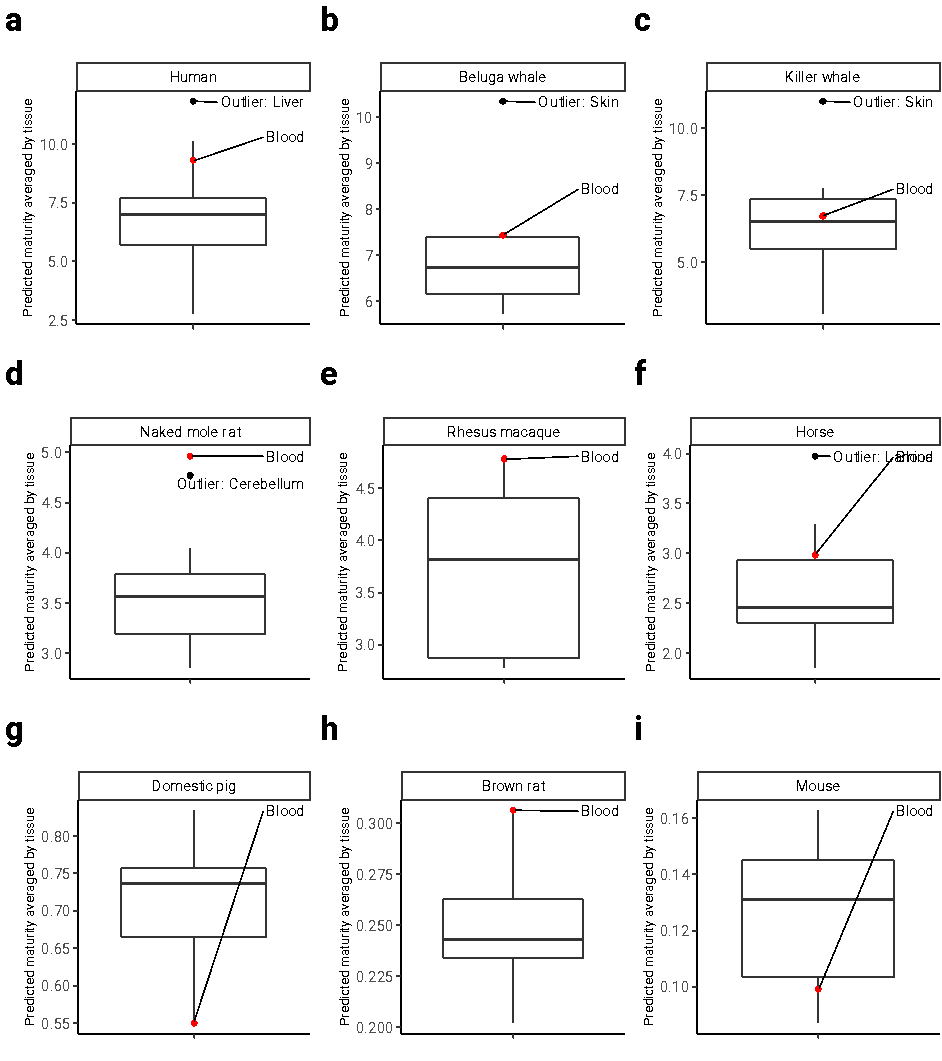

#### Fig. S7. Predicted time to sexual maturity in select species for which multiple tissues were available.

Tissue-agnostic predictor of time to sexual maturity. The boxplot shows median predicted values (short horizontal line) across tissue types. Significantly outlying tissues have been highlighted. The boxplot, as implemented in the R programming language, provides a visual summary of key statistics from a dataset: The median is represented by the horizontal line inside the box. The interquartile Range (IQR) encompasses the middle 50% of the data. The box's upper boundary represents the 75th percentile, while the lower boundary represents the 25th percentile. The IQR is the difference between these two values. The whiskers extend to the most extreme data point which is no more than 1.5 times the interquartile range from the box. Tissue-agnostic predictor of time to sexual maturity predictor, based on averaged species methylation, was used to predict individual sample time to sexual maturity (in log years). The predicted values are aggregated by taking the mean lifespan predictions by tissue groups. Panels **a-i** convert log scale back to original units (age at sexual maturity in years); only species with more than 6 different tissue types are shown; mean tissue predicted value outliers are annotated; Tissue type “H.Stem.Progenitor.LSK” stands for “LSK Progenitor Hematopoietic Stem cells.”

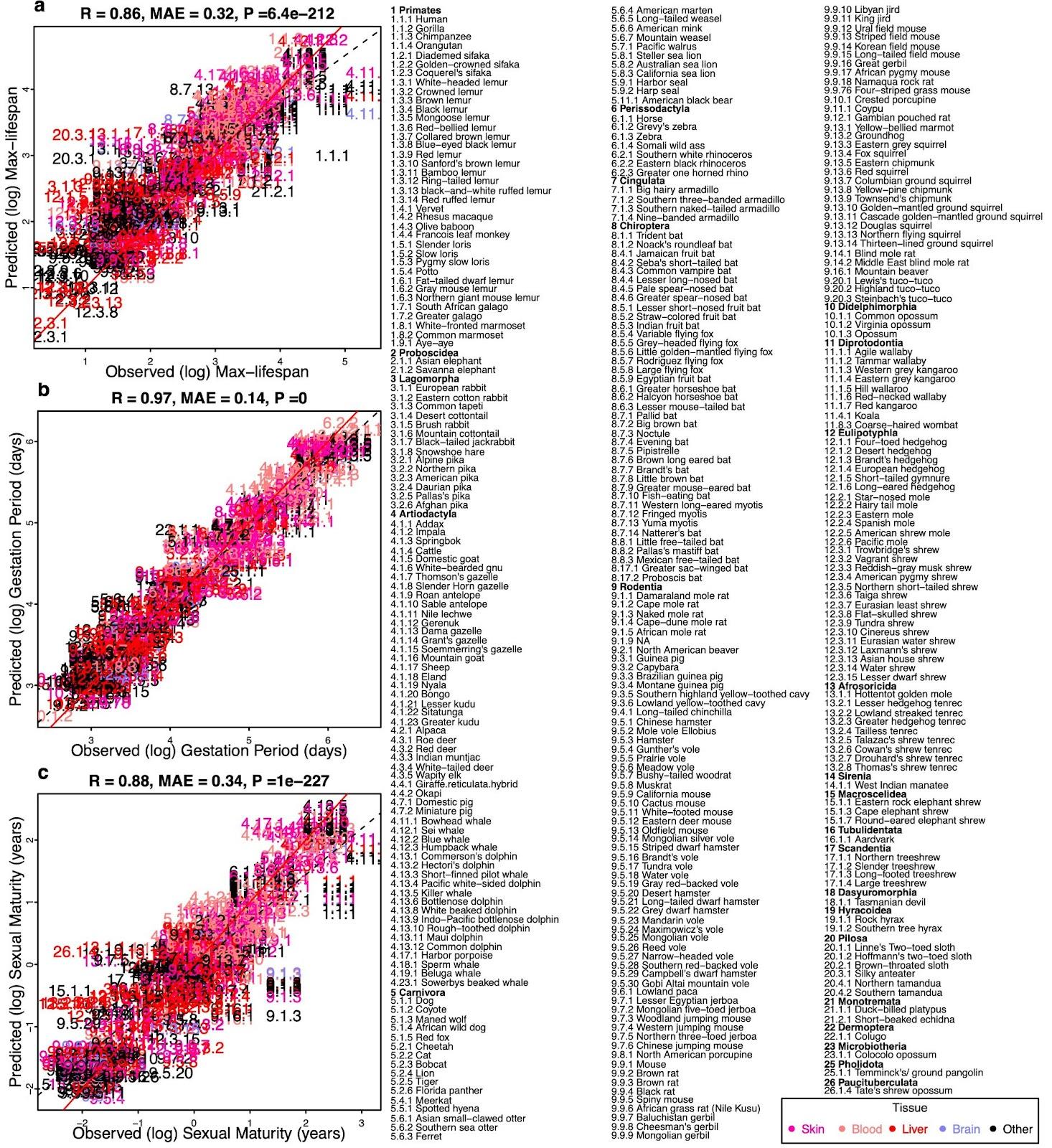

#### Fig. S8 | Tissue-aware predictors trained on species-tissue combinations.

A penalized joint linear model used to predict species lifespan (Elastic Net). Same framework as that of **Fig. 1**, except that it distinguishes tissue types. CpG probes are averaged by each species-tissue combination. Different tissues within the same species share the same maximum lifespan but retain different methylation levels. Three panels show predictors for **a,** log maximum lifespan (in log years), **b,** log-transformed gestation time (in log days), and **c,** log-transformed age at sexual maturity (in log years). Designated Mammalian numbers in scatter plot panels and the Figure legend are the same as those of main **Fig. 1.** MAE abbreviates median absolute errors from the regression errors; r and p denote Pearson’s correlation and p-values, respectively. Numbers and colors are the mammalian species number and order annotation consistent with those of other Figures. Numeric values can be found in **table S3**. As with the **Fig. 1**, species appear as designated numbers in scatter plot panels; the corresponding common names and taxonomic orders are annotated in Figure legends; the whole number (number before the decimal separator) part of each mammalian number is assigned in accordance with the corresponding taxonomic order. Red solid line represents the perfect prediction line, and the dotted line represents the fitted linear regression line.

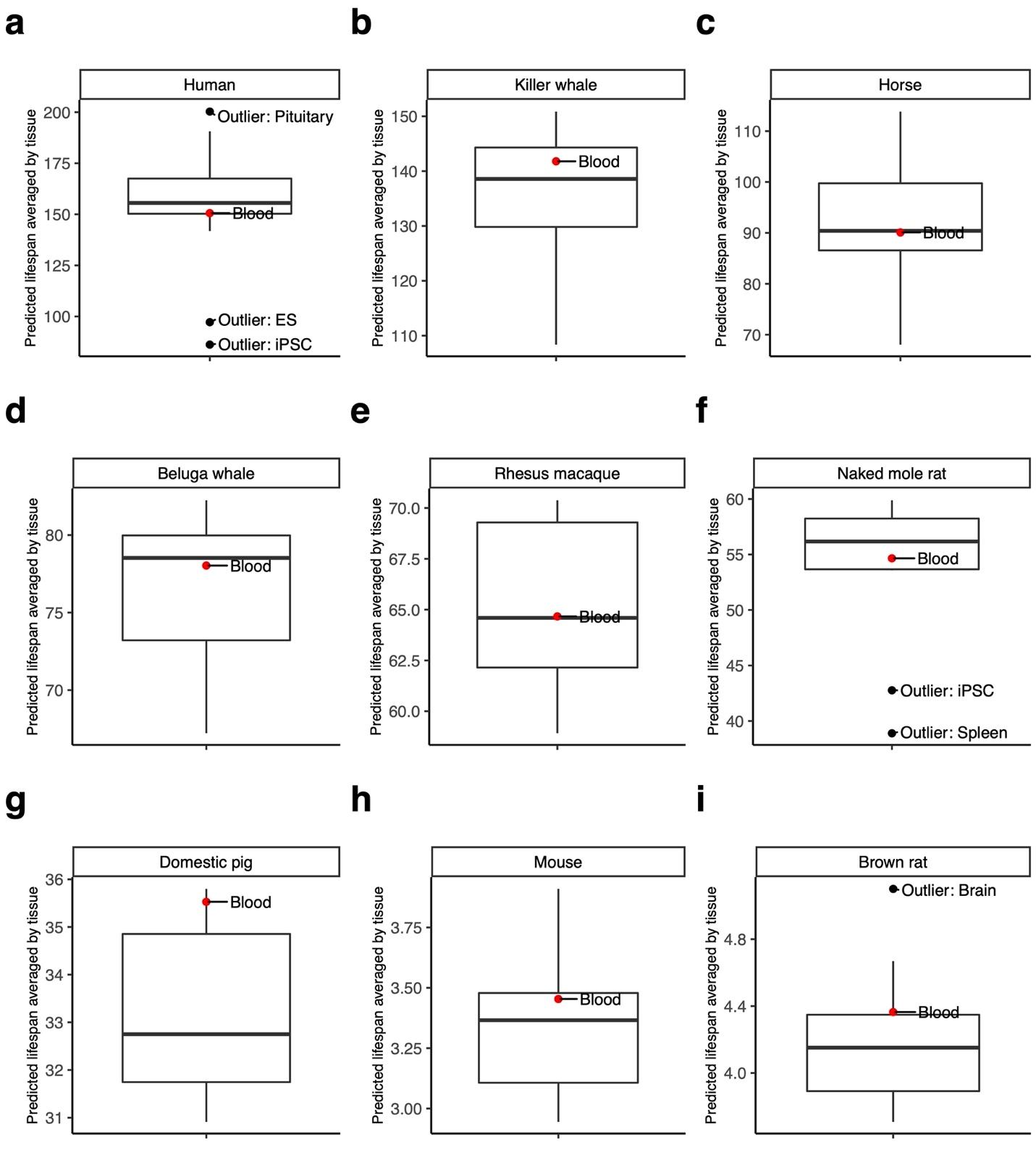

#### Fig. S9 | Tissue group differences in predicted mammalian maximum lifespan – Tissue-Aware

Tissue-aware predictor of mammalian lifespan, based on averaged species methylation, was used to predict individual sample lifespan (in log years). The predicted values are aggregated by taking the mean lifespan predictions by tissue groups. Panels **a-i** convert log scale back to original units (lifespan in years); only species with more than 6 different tissue types are shown; mean tissue predicted value outliers are annotated; Tissue type “H.Stem.Progenitor.LSK” stands for “LSK Progenitor Hematopoietic Stem cells.”

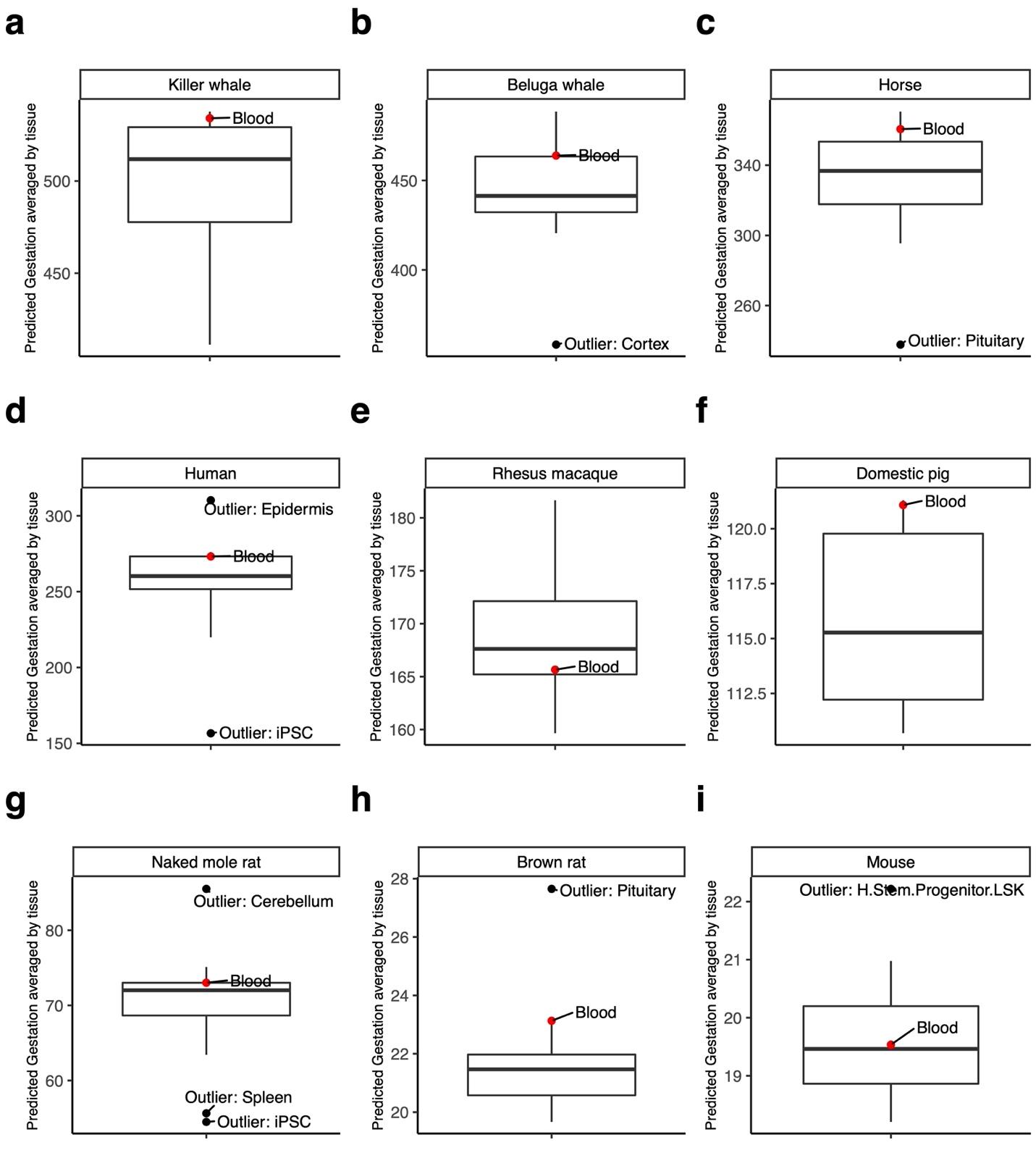

#### Fig. S10 | Tissue groups differences in predicted mammalian gestation time – Tissue-aware.

Tissue-aware predictor of gestation time, based on averaged species methylation, was used to predict individual sample gestation time (in log days), trained on tissue-aware data. The predicted values are aggregated by taking the mean gestation time predictions by tissue groups. Panels **a-i** convert log scale back to original units (gestation in days); only species with more than 6 different tissue types are shown; mean tissue predicted value outliers are annotated; Tissue type “H.Stem.Progenitor.LSK” stands for “LSK Progenitor Hematopoietic Stem cells.”

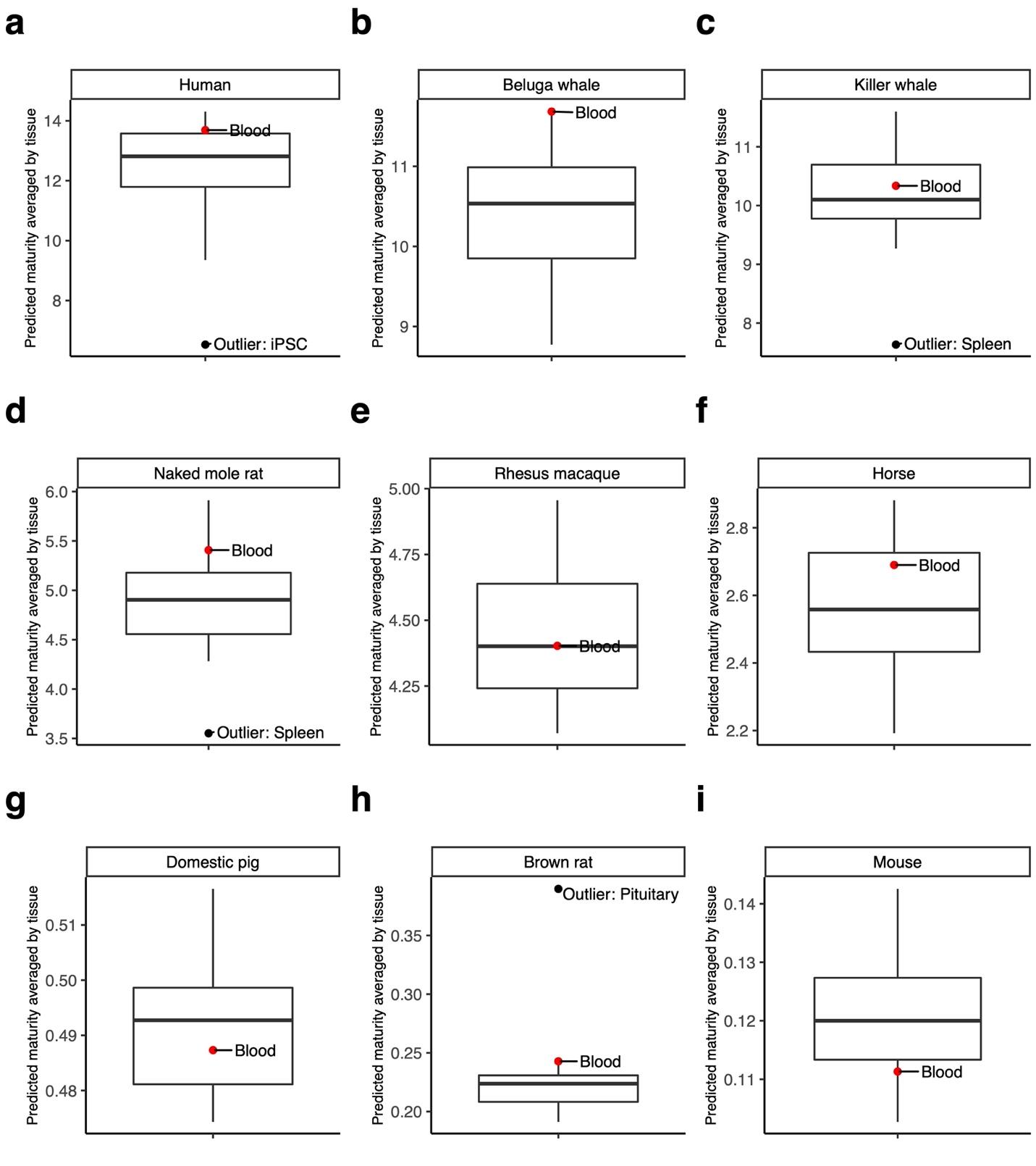

#### Fig. S11 | Tissue groups differences in predicted mammalian sexual maturity time – Tissue-aware.

Mammalian times to sexual maturity predictor, based on averaged species methylation, was used to predict individual sample time to sexual maturity (in log years), trained on tissue-aware data. The predicted values are aggregated by taking the mean lifespan predictions by tissue groups. Panels **a-i** convert log scale back to original units (age at sexual maturity in years); only species with more than 6 different tissue types are shown; mean tissue predicted value outliers are annotated; Tissue type “H.Stem.Progenitor.LSK” stands for “LSK Progenitor Hematopoietic Stem cells.

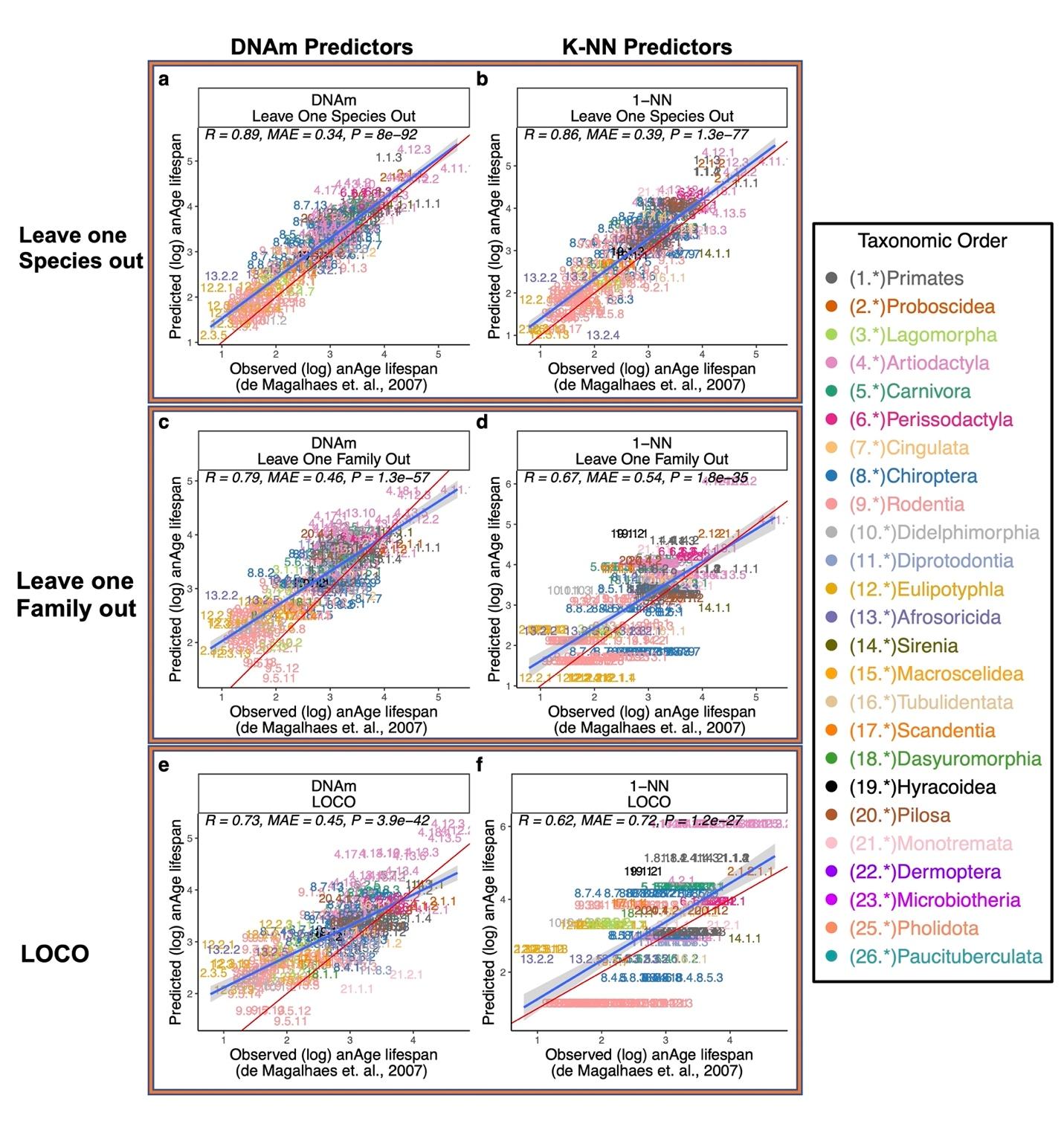

#### Fig. S12 | Overall comparisons between DNAm lifespan predictors and phylogeny-based predictors.

Various training-test validation analyses of predictors of log (base e) transformed estimates of maximum lifespan. We compared prediction performance between DNAm elastic net predictors and 1-Nearest-Neighbor predictor (k-NN). 1-Nearest-Neighbor predictor utilizes distances from the Mammalian phylogenetic TimeTree (*54*). Results under different training-test separation methods are shown in panels **a**, **b**, DNAm and k-NN predictors test set predictions under leave-one-species-out (LOSO) training-test separation scheme; **c**, **d**, DNAm and k-NN predictors test set predictions under leave-one-family-out training-test separation; **e**, **f**, DNAm and k-NN predictors test set predictions under leave-one-clade-out (LOCO) training-test separation. LOCO (leave-one-clade-out) is defined as, for orders with more than 20 species (Rodentia, Artiodactyla, Chiroptera, Primates, Carnivora, and Eulipotyphla), leaving out all member species except the longest-living and shortest-living species. MAE abbreviates median absolute errors from the regression errors; r and p denote Pearson’s correlation and p-values, respectively. Numbers and colors are the mammalian species number and order annotation consistent with those of other Figures. Numeric values can be found in **table S1**. Shaded areas represent 95% confidence intervals of the simple linear regression line. E).

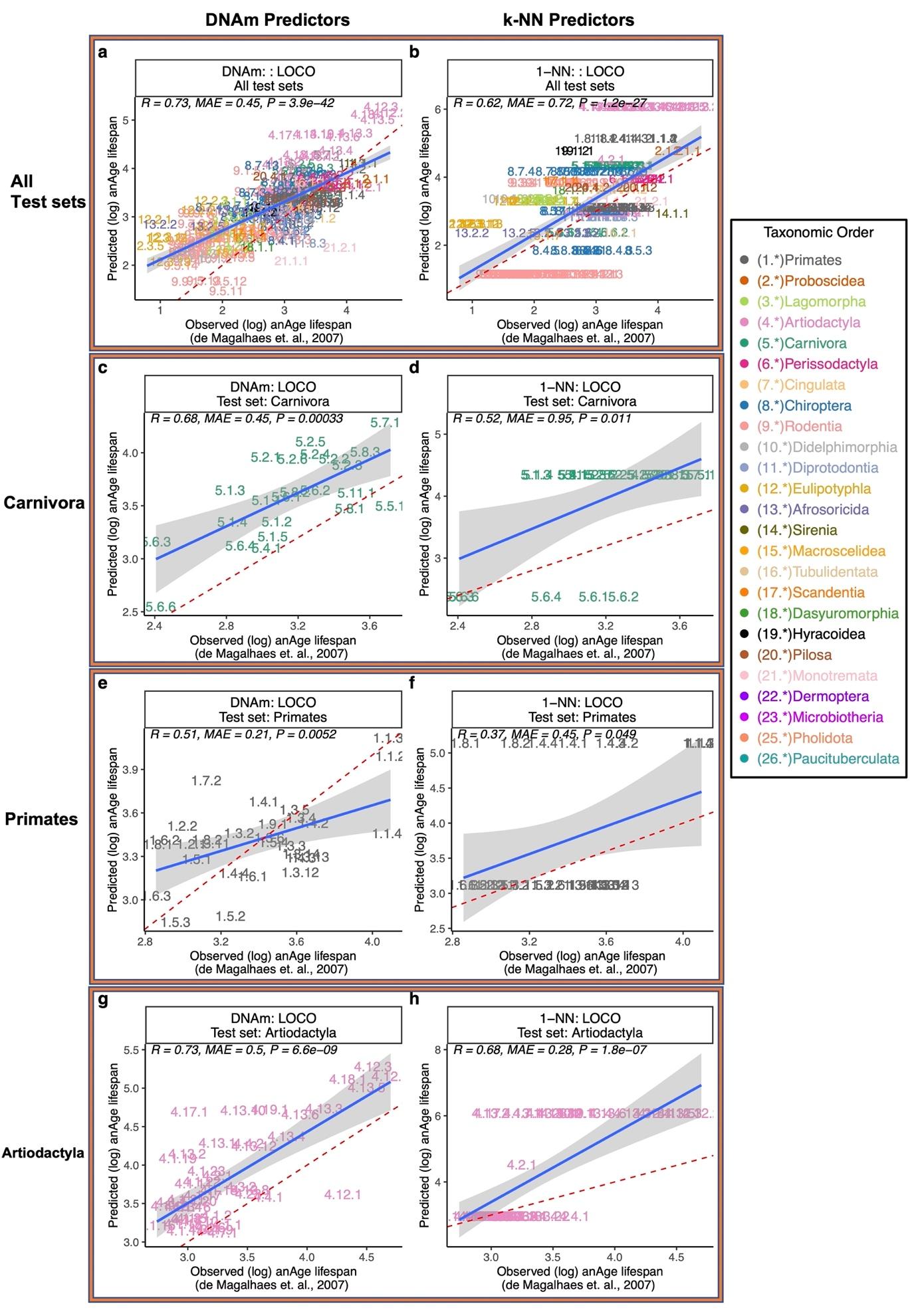

#### Fig. S13 | Taxonomic order breakdown of DNAm lifespan predictors and Phylogeny-based Predictors under LOCO.

A breakdown of predictor performance in large taxonomic orders under LOCO. We compared prediction performance between DNAm elastic net predictors and 1-Nearest-Neighbor predictor (k-NN). 1-Nearest-Neighbor predictor utilizes distances from the Mammalian phylogenetic TimeTree (*54*). **a**, DNAm predictor’s test set predictions leave-one-clade-out (LOCO) training-test separation scheme; **b**, k-NN predictor’s test set predictions under LOCO; **c**, **d**, DNAm and k-NN predictors, respectively, test set predictions of lifespan for all species belonging to Carnivora under LOCO; **e**, **f**, DNAm and k-NN predictors, respectively, test set predictions of lifespan for all species belonging to Primates under LOCO; **g**, **h** DNAm and k-NN predictors, respectively, test set predictions of lifespan for all species belonging to Artiodactyla under LOCO. MAE abbreviates median absolute errors from the regression errors; r and p denote Pearson’s correlation and p-values, respectively. Numbers and colors are the mammalian species number and order annotation consistent with those of fig. S1. Numeric values can be found in **table S1**. Shaded areas represent 95% confidence intervals of the simple linear regression line. Panels **a** and **b** are analogous to those of **Fig. 2c,d**.

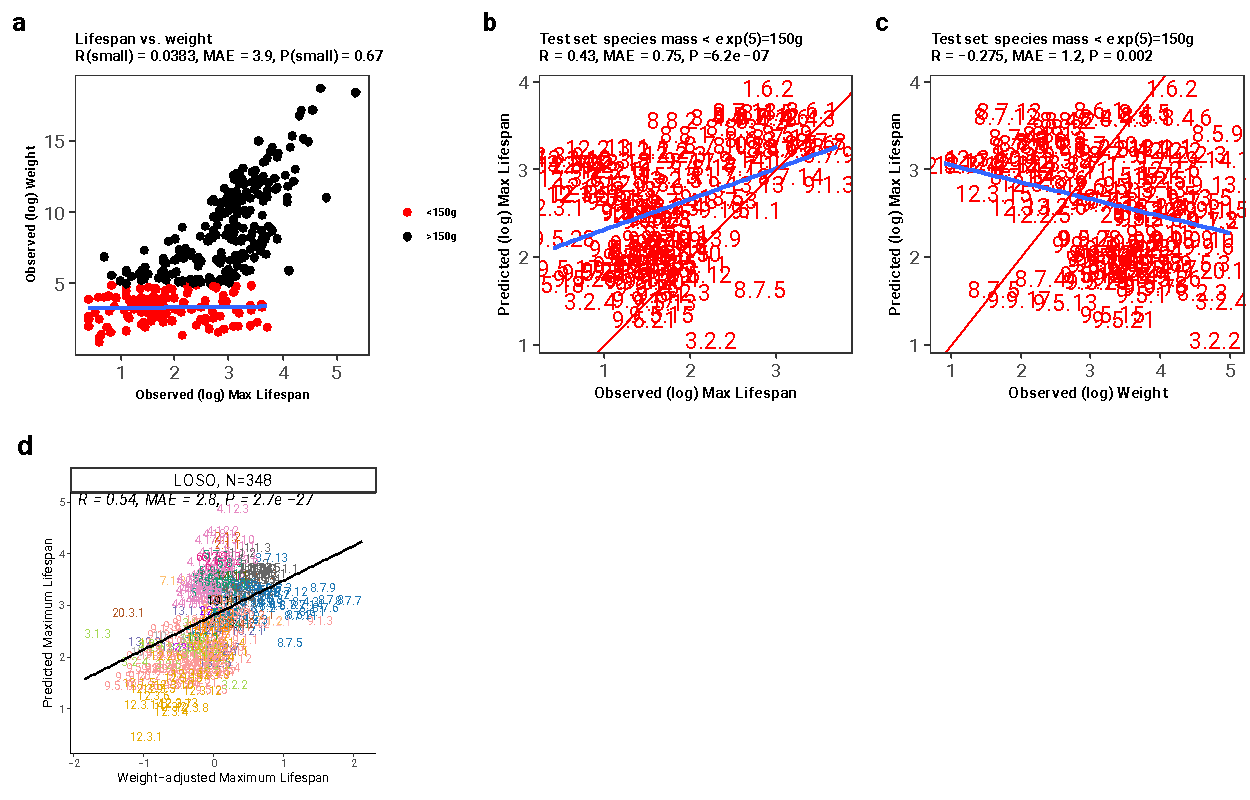

#### Fig. S14 | DNAm lifespan predictions do not reflect confounding by adult weight.

**a-c**, Report results for a DNAm max lifespan predictor trained on mammal species with an average weight under 150 grams (small mammals). Panels **a**, observed (log) adult body weight vs. observed (log) maximum lifespan in all mammalian species within the data set, color-coded by small-size indicator (more than 150 grams); **b**, test set predictions for the maximum lifespan in small-sized (<150 grams) mammalian species vs. observed (log) maximum lifespan; **c**, test set predictions for the maximum lifespan in small-sized (<150 grams) mammalian species vs. observed (log) adult body weight. MAE abbreviates median absolute errors from the regression errors; r and p denote Pearson’s correlation and p-values, respectively. Numbers are the mammalian species number annotation consistent with those of other Figures. Numeric values can be found in **table S1**. Shaded areas represent 95% confidence intervals of the simple linear regression line.

**d.** Results for the final version of the tissue-agnostic DNAm predictor of maximum lifespan. Predicted maximum lifespan (on the log scale, y-axis) versus the corresponding adult weight adjusted version (x-axis). Specifically, the weight adjusted version of log maximum lifespan was defined as raw residual resulting from regressing log transformed predicted maximum lifespan on the log transformed average adult weight of the species.

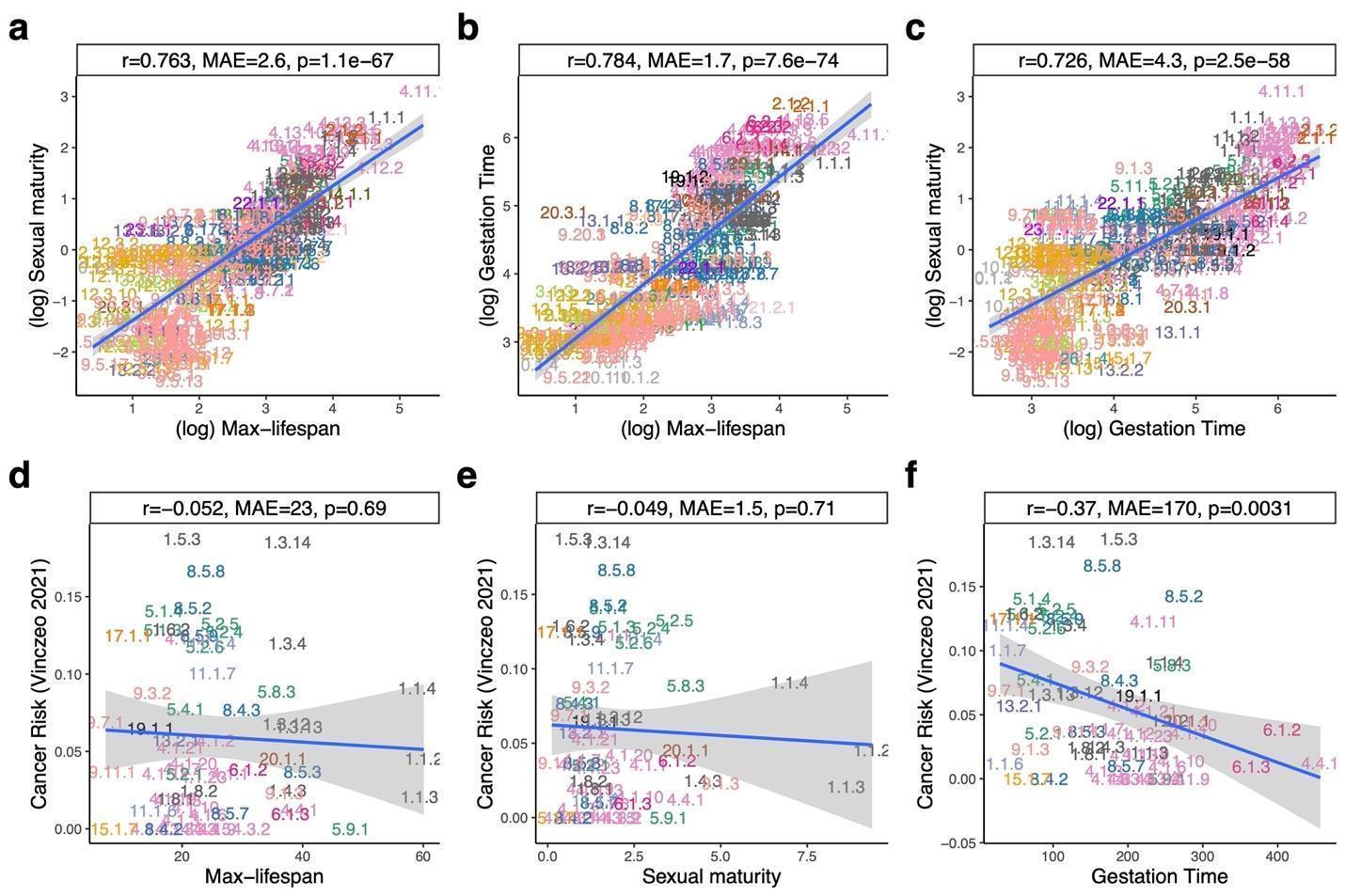

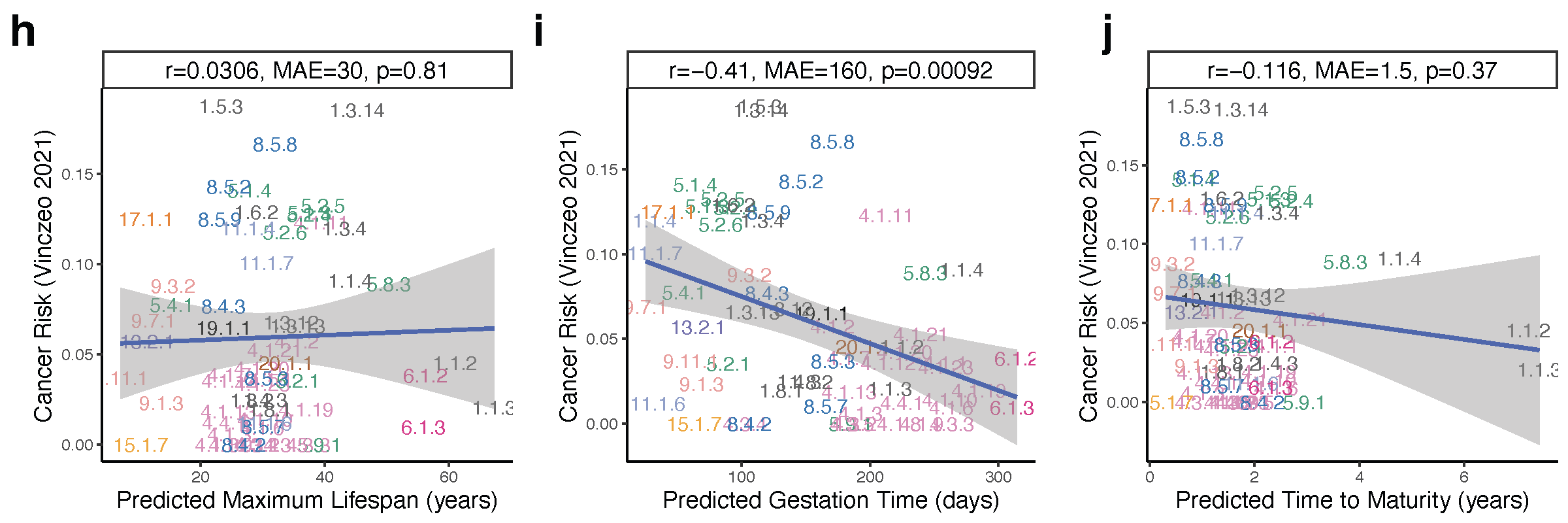

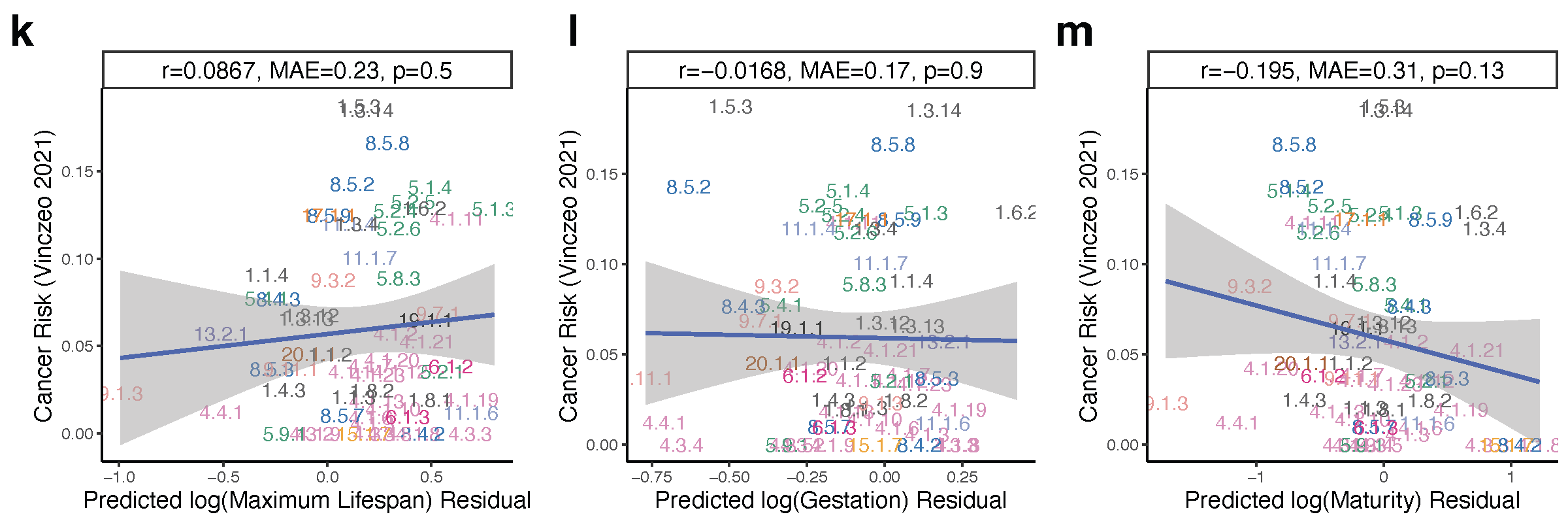

#### Fig. S15 | Relationships between observed and epigenetic estimates of mammalian life history traits, including mammalian cancer risk.

**a-f**, Panels depict log-transformed relationships between observed variables: **a**. Age at sexual maturity and maximum lifespan **b**. Gestation time and maximum lifespan **c**. Sexual maturity time and gestation time **d**. Cancer risk and maximum lifespan **e**. Cancer risk and sexual maturity **f.** cancer risk and gestation time. **h-j**, estimates of mammalian cancer risk (Vinczeo 2021, y-axis) are plotted against their corresponding epigenetic estimates: **h.** Maximum lifespan **i**. Gestation time **j**. age at sexual maturity. **k-m**, this set is analogous to **h-j**, but the x-axis reports residuals derived from regressing the epigenetic estimate of the life history trait on its observed value (on the log scale): **k**. Log maximum lifespan **l**. Log gestation time **m**. Log-transformed age at sexual maturity. "MAE" represents median absolute errors from the regression errors, while "r" and "p" signify Pearson’s correlation and p-values, respectively. Numbering and colors correspond to the mammalian species number and order, consistent with those in **Fig. 1**. Shaded areas illustrate the 95% confidence intervals of the simple linear regression line. log denotes the natural logarithm, i.e., base e.

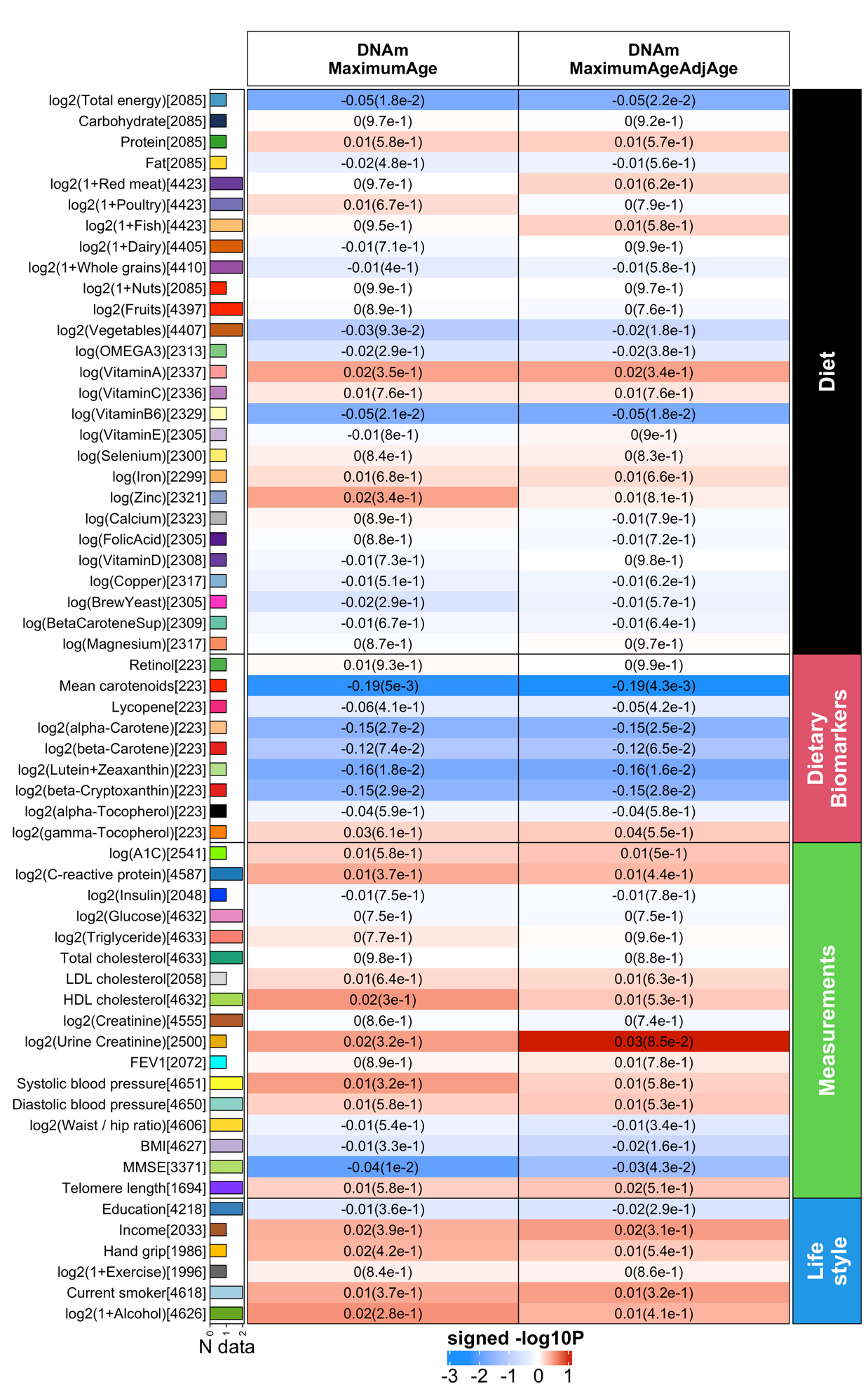

#### Fig. S16. Human epidemiological cohort studies of diet and clinical biomarkers.

We performed a correlation analysis between (1) our methylation-based estimator of maximum lifespan (first column) and its age adjusted version (second column) and (2) 59 variables spanning diet, clinically relevant measurements, and lifestyle factors. Comprehensive details of these variables can be found in (*18*). We conducted a robust correlation analysis (biweight midcorrelation, bicor) between (1) our methylation based measures (columns), and (2) 59 variables encompassing 27 self-reported dietary factors, 9 dietary biomarkers, 17 clinical measurements related to vital signs, metabolic traits, inflammatory markers, cognitive and lung function, central adiposity, leukocyte telomere length, and 6 lifestyle factors. This bicor analysis was applied to individuals from both the Framingham Heart Study (up to n=2544) and Women's Health Initiative (up to n=2107), stratified by gender and ethnic category within each respective cohort. The results were consolidated using fixed-effects meta-analysis models, weighted by inverse variance, generating a meta-estimate of bicor and meta P-value. The clinical biomarkers in FHS offspring cohort were measured during the 8th examination aligned with the measures of DNA methylation profiles. The 9 dietary biomarkers, however, were only available in the WHI cohort, with measurements taken from fasting plasma collected at baseline. Food groups and nutrients considered were comprehensive, encompassing all types and preparation methods; for instance, folic acid included both synthetic and natural forms, and dairy encompassed cheese and all varieties of milk. Further details on the individual diet variables of the WHI cohort can be found in our previous study (*57*).
